# Detailed profiling with MaChIAto reveals various genomic and epigenomic features affecting the efficacy of knock-out, short homology-based knock-in and Prime Editing

**DOI:** 10.1101/2022.06.27.496697

**Authors:** Kazuki Nakamae, Mitsumasa Takenaga, Shota Nakade, Akinori Awazu, Naoaki Sakamoto, Takashi Yamamoto, Tetsushi Sakuma

## Abstract

Highly efficient gene knock-out and knock-in have been achieved by harnessing CRISPR-Cas9 and its advanced technologies such as Prime Editor. In addition, various bioinformatics resources have become available to quantify and qualify the efficiency and accuracy of CRISPR edits, which significantly increased the user-friendliness of the general next-generation sequencing (NGS) analysis in the context of genome editing. However, there is no specialized and integrated software for investigating the preference in the genomic context involved in the efficiency and accuracy of genome editing using CRISPR-Cas9 and beyond. Here, we address this issue by establishing a novel analysis platform of NGS data for profiling the outcome of template-free knock- out and short homology-based editing, named MaChIAto (**M**icrohomology- **a**ssociated **Ch**romosomal **I**ntegration/editing **A**nalysis **to**ols) (https://github.com/KazukiNakamae/MaChIAto). MaChIAto accommodates the classification and profiling of the NGS reads to uncover the tendency of the corresponding method of genome editing. In the profiling function, MaChIAto can summarize the mutation patterns along with the editing efficiency, and > 70 kinds of feature analysis, e.g., correlation analysis with thermodynamics and secondary structure parameters, are available. Additionally, the classifying function of MaChIAto is based on, but much stricter than, that of the existing tool, which is achieved by implementing a novel method of parameter adaptation utilizing Bayesian optimization. To demonstrate the functionality of MaChIAto, we analyzed the NGS data of knock- out, short homology-based knock-in, and Prime Editing. We confirmed that some features of (epi-)genomic context affected the efficiency and accuracy. These results show that MaChIAto is a helpful tool for understanding the best design for CRISPR edits. More importantly, it is the first tool for discovering features in the short homology-based knock-in outcomes. MaChIAto would help researchers profile editing data and generate prediction models for CRISPR edits, further contributing to revealing a “black box” process to produce a variety of CRISPR and Prime Editing outcomes.

---

CRISPR gene editing has recently been applied in numerous studies. Bioinformatics tools are required for quantifying edited sequence reads from targeted amplicon sequencing analysis (TASA). One of the representative TASA tools^1–4^ is CRISPResso^1, 2^, which has been adopted in various CRISPR experiments (Supplementary Note 1). However, CRISPResso and the other tools have never dealt with the following two issues. First, the tools do not function enough to obtain a comprehensive understanding of the various tendencies on editing efficiency such as the relationship with thermodynamics and secondary structure features. The analysis requires the calculation pipeline that estimates the values from the input sequence information and then aggregates the variable results obtained from multiple samples. The second issue is the instability of the application performance depending on parameter settings. Researchers who use the TASA tools need to adjust the analysis parameter of the TASA tools to stably estimate the *bona fide* editing rate, although the process is difficult because the quality of the sequencing data varies per experiment.

To address these issues, we have developed MaChIAto (**M**icrohomology-**a**ssociated **Ch**romosomal **I**ntegration/editing **A**nalysis **to**ols); a comprehensive analysis software that can precisely classify, deeply analyze, correctly align, and thoroughly review the TASA data obtained by various CRISPR experiments, including template-free gene knock-out, short homology-based gene knock-in, and even a new-class CRISPR methodology, Prime Editing^5^ (Fig. 1a). MaChIAto harnesses the variants table from CRISPResso and re-classifies sequencing reads based on, but much stricter than, that of CRISPResso2 (Fig. 1b, Supplementary Note 2, 3, 5, and 9), which is achieved by implementing a novel parameter adaptation method utilizing Bayesian optimization (Methods and Supplementary Note 2). Thus, MaChIAto obtains an optimized accurate classification using the auto-adjustment of the parameters per input. Furthermore, MaChIAto can estimate the > 70 feature values such as secondary structures and profile mutation patterns along with editing efficiency to reach a deeper understanding of various CRISPR edits (Fig. 1c–e).

**Figure 1.**
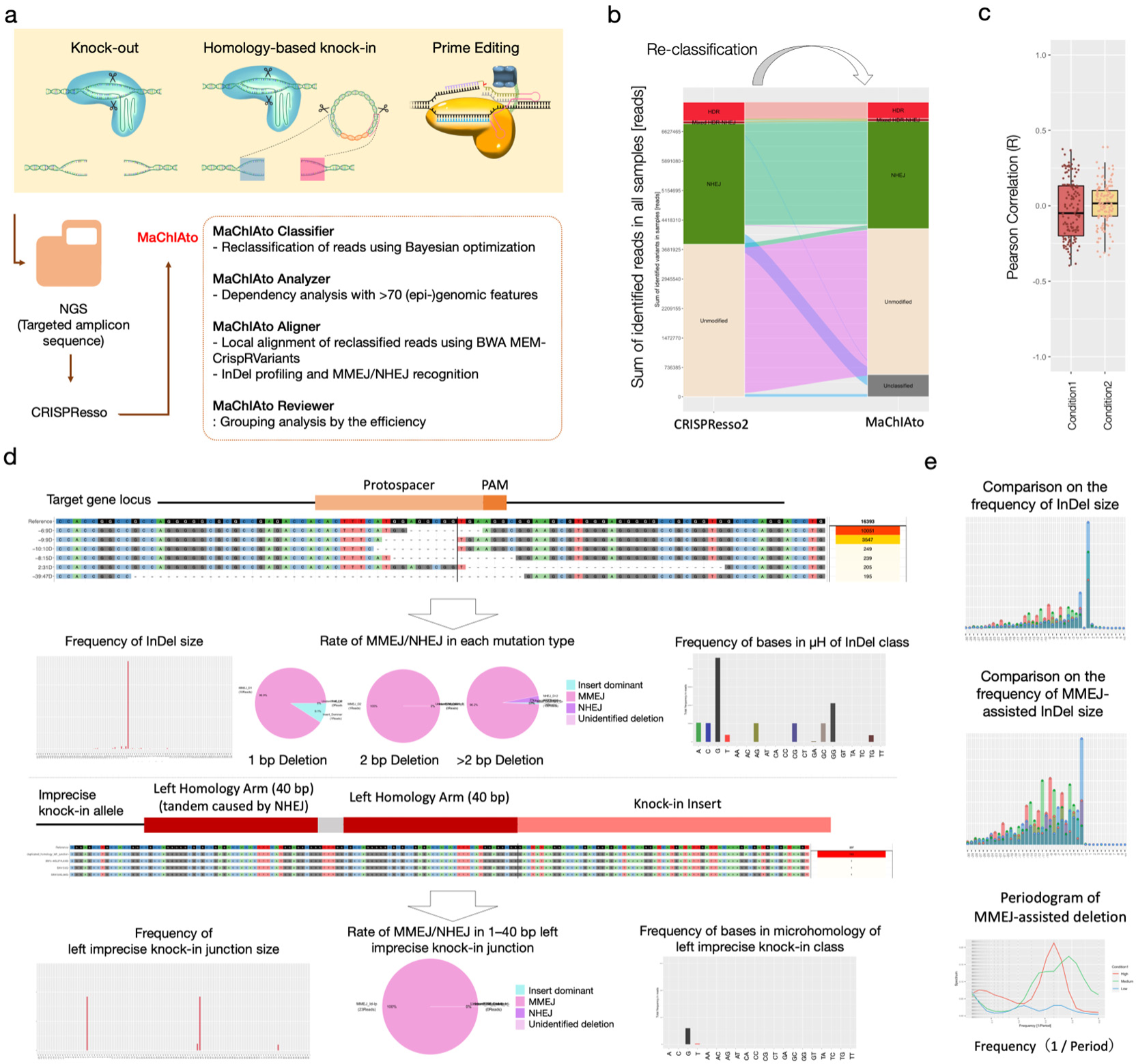
Overview of MaChIAto. a. Overview of the process in the CRISPResso-MaChIAto pipeline to profile the amplicon sequencing data from next-generation sequencing (NGS). The pipeline receives the amplicon sequencing result of CRISPR-based knock-out and knock-in, and Prime Editing as input. MaChIAto utilizes the output of CRISPResso, Alleles_frequency_table.txt (or .zip), to execute the following four programs. (1) MaChIAto Classifier optimizes parameters for better classification and then performs stricter re-classification. Next, (2) MaChIAto Analyzer runs a dependency analysis between each classification outcome and >70 biological features. (3) MaChIAto Aligner makes local alignment and variants distribution on the target site and knock-in junction using BWA MEM-CrispRVariants. Additionally, the program qualifies MMEJ-assisted InDels. (4) MaChIAto Reviewer uncovers a tendency of the efficient and inefficient editing group by aggregating and plotting values that MaChIAto Aligner mainly generates. b. Re-classification result of the MaChIAto Classifier. MaChIAto Classifier rigorously and comprehensively re-classifies the result of CRISPResso version1 and CRISPResso version2 (CRISPResso2) (Methods and Supplementary Note 2). The alluvial diagram represents changes in re-classification. The left stacked bar plot shows the result of the original classification of CRISPResso2, and the right one shows the result of re-classification of the MaChIAto Classifier. White, green, dark red, and red boxes indicate Unmodified, NHEJ (includes MMEJ-assisted InDels in this section, also called InDels class), Mixed HDR-NHEJ (represents imprecise editing class), HDR (represents precise editing class) classes, respectively. Variants in NHEJ class have InDels but no knock-in insert. The definition of the classes follows the way of CRISPResso. The grey box represents Unclassified class, in which ambiguous sequence is classified (Methods and Supplementary Note 9), which is originally accommodated in MaChIAto. The detailed explanation of MaChIAto Classifier is described in the Supplementary Information and online manual (https://machiatopage.github.io). c. Example of the result output by MaChIAto Analyzer. MaChIAto Analyzer estimates Pearson correlations between >70 biological features and the efficacy, such as the editing efficiency. The box plot is a summary of the correlations. The left box shows the case of condition1 (control) result, and the right one shows the case of condition2 result. The correlation estimation is with statistic tests. The detailed figures are shown in Supplementary Note 6. The detailed explanation of MaChIAto Analyzer is described in the Supplementary Information and online manual (https://machiatopage.github.io). d. Example of the result output by MaChIAto Aligner. MaChIAto Aligner identifies variants based on the local alignment of BWA MEM-CrispRVariants pipeline using unmodified and expected knock-in sequences as the reference sequence, and MaChIAto Aligner summarizes and visualizes the result by InDel position, size, and kinds (MMEJ and NHEJ) of base. The number of output figures can be >100 items. The detailed explanation of MaChIAto Aligner is described in the Supplementary Information and online manual (https://machiatopage.github.io). e. Example of the result output by MaChIAto Reviewer. MaChIAto Reviewer summarizes and visualizes the result by InDel position, size, type of repair system (MMEJ and NHEJ), and kinds of base that MaChIAto aligner identifies. The number of output figures can be >100 items. The detailed explanation of MaChIAto Reviewer is described in the Supplementary Information and online manual (https://machiatopage.github.io).

MaChIAto introduces the five novel functions for profiling: first, alignment of the target site and knock-in junction (Fig. 1d, alignment, Supplementary Note 6); second, insertion and/or deletion (InDel) profiling (Fig. 1d, left and right bar plots, Supplementary Note 6); third, microhomology-mediated end-joining (MMEJ)/nonhomologous end-joining (NHEJ) recognition (Fig. 1d, center pie charts, Supplementary Note 6); forth, dependency analysis between the sequencing outcome and > 70 (epi-)genomic features (Fig. 1c, Supplementary Note 6); and fifth, grouping analysis by the editing efficiency (Fig. 1e, Supplementary Note 6).

The applicability and superiority of MaChIAto in analyzing the CRISPR editing data were confirmed for knock-out, knock-in, and Prime Editing. We performed the experiments and newly obtained the TASA data for a series of knock-out and short homology-based knock-in^6–8^ targeting 40 gene loci in HEK293T cells (Supplementary Note 4). We analyzed the data with MaChIAto and obtained better classification results than those of CRISPResso (Supplementary Note 5). Next, MaChIAto investigated dependency between 130 features and the editing efficiency of knock-out and knock- in. As a result, some (epi-)genomic features such as DNaseI accessibility and propeller twist of homology arm were revealed to affect the accuracy of knock-in. In the group analysis, MaChIAto identified that one base pair insertion was a crucial factor in the knock-in efficiency (Supplementary Note 6). Next, we applied MaChIAto to profile the publicly available Prime Editing data^5^ (Supplementary Note 9). Our re-classification pipeline successfully enabled to reduce the percentages of the reads erroneously classified as precisely edited alleles from 8.6% to 0% in the best case. Furthermore, MaChIAto uncovered the difference between the groups with high and low efficiency in the distribution of deletion positions and predicted values such as the energy in tetranucleotides of the RT template.

In conclusion, we developed a new integrative TASA tool named MaChIAto and successfully demonstrated its utility to clear up the sequential details and the involvement of (epi-)genomic contexts in template-free knock-out, short homology- based knock-in, and Prime Editing. MaChIAto offers the first bioinformatics pipeline for discovering *a priori* sequence and structural features involved in a “black box” process to produce a variety of gene editing outcomes. By modifying MaChIAto, it would also be applied for other approaches such as base editing^9^ and ssODN knock- in^10^. In the future, MaChIAto would help researchers to profile the editing data and find the characteristics that help an efficiency prediction (Supplementary Note 6–9). Our analysis using MaChIAto is easily reproducible without specialized equipment (Supplementary Note 10).

## Acknowledgments

We would like to acknowledge the technical assistance of Tomoko Kurisu (Hiroshima University) for plasmid construction, and Yusaku Wada and Keita Tsukahara (FASMAC Co. Ltd.) for next-generation sequencing. We also thank Feng Zhang (Broad Institute) for the provision of reagents through Addgene and David R. Liu (Merkin Institute and Broad Institute) for the provision of Prime Editing data through SRA. We are also grateful for previous works: especially, Luca Pinello (Massachusetts General Hospital and Harvard Medical School) who built CRISPResso^1, 2^, and Mark D. Robinson (University of Zurich) who established CrispRVariants^3^, which enabled to develop MaChIAto.

## Author Contributions

K.N. and T.S. conceived and designed the study. K.N. developed MaChIAto software. K.N., M.T., S.N, A.A., and T.S, contributed to the design and preparation of knock-out and knock-in experiments. K.N performed the knock-out and knock-in experiments.

K.N. analyzed the amplicon sequencing data of knock-out, knock-in, and Prime Editing.

S.N. provided instructions on the experiment. K.N. and N.S. contributed to an online release of MaChIAto. T.Y. and T.S. supervised the study. K.N. and T.S. wrote the manuscript with feedback from all authors.

## Supplementary Information

### Code availability

MaChIAto source code is available online at https://github.com/KazukiNakamae/MaChIAto. The code used for the analyses in this paper and the additional manuscripts are available at https://machiatopage.github.io.

### Data availability

The data used in Supplementary Note 3 was obtained from the additional manuscript of ref. 3 (https://github.com/markrobinsonuzh/CrispRvariants_manuscript) and SRR1769728. The danRer7 reference genome was obtained from UCSC Genome Browser database (http://hgdownload.cse.ucsc.edu/goldenPath/danRer7/bigZips/danRer7.fa.gz). We downloaded the call sets from the ENCODE portal (https://www.encodeproject.org/) for making the extra table in MaChIAto Analyzer. The example files for MaChIAto are available at our Source Forge (https://sourceforge.net/projects/machiato-example-files/). The raw NGS amplicon sequencing data generated in Supplementary Note 4 can be provided if there is a request to the authors. We performed the meta-analysis of Supplementary Note 9 using the high-throughput sequencing data at the NCBI Sequence Read Archive database under accession PRJNA565979.

## Methods

### Implementation of MaChIAto (**M**icrohomology-**a**ssociated **Ch**romosomal **I**ntegration/editing **A**nalysis **to**ols)

See the online manual (https://machiatopage.github.io) for the detailed usage and limitation of MaChIAto.

MaChIAto is a third-party tool of the CRISPResso series (CRISPResso and CRISPResso2), which was built with Python and R. MaChIAto consists of four components; “MaChIAto Classifier,” “MaChIAto Analyzer,” “MaChIAto Aligner,” and “MaChIAto Reviewer.”

MaChIAto Classifier can re-classify CRISPResso and CRISPResso2 results to generate stricter classifications. The classes are “Unmodified,” “Deletion,” “Insertion,” “Frequent Substitution,” “Imprecise Editing,” and “Precise Editing.” The tool can also calculate values such as “Editing Efficiency” and “Precise Knock-in Efficiency” based on the MaChIAto result. MaChIAto Classifier is one of the core components of MaChIAto and is located as the initial step in MaChIAto because it generates data that is used as input for MaChIAto Analyzer, MaChIAto Aligner, and MaChIAto Reviewer.

MaChIAto Classifier accepts the output file: “Alleles_frequency_table.txt” and some parameters of the CRISPResso series. If knock-in analysis is performed, each homology arm length is also required as input. In the initial MaChIAto Classifier process, random sequences are temporally recognized as homology arms and indicators. The definition of indicators is similar to that in Cas-Analyzer^11^. Four indicators of MaChIAto Classifier are used for the recognition of the following sequences: on-target sequence, knock-in sequence, and the sequences that are located around the cutting site and on knock-in donor sequence (Supplementary Fig. 3, callout). To decrease false- negative detection, MaChIAto Classifier automatically chooses the proper length of indicators and alignment thresholds to detect more on-target sequences and knock-in sequences in the total reads. Specifically, MaChIAto Classifier runs four-step Bayesian optimization: 1) optimizing out-out indicators, 2) optimizing out-out alignment thresholds, 3) optimizing in-out indicators and 4) optimizing in-out alignment thresholds. Each set of information, which are the purpose, adjustable parameters, and calculation, is explained below. The purpose of the “optimizing out-out indicators” step is to maximize the rate of detected variants (RDV) in each class of CRISPResso: RDVHDR, RDVMixed, RDVNHEJ, and RDVUnmodified.

**Supplementary Figure 1.**
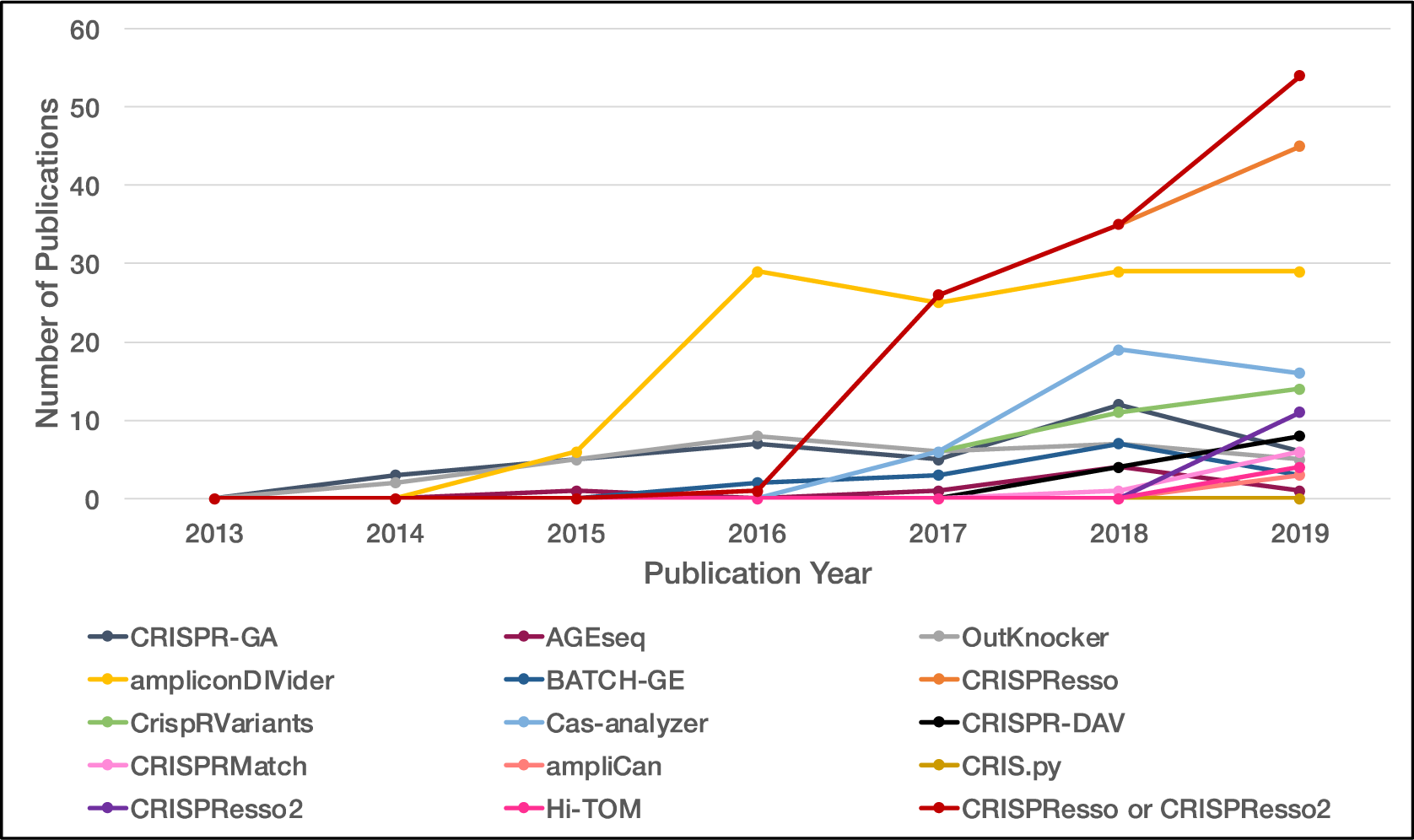
The trend in analysis tools for amplicon sequencing in the context of genome editing (2020/05/24 in PubMed). The lines indicate changes in the numbers of citations, and their colors represent the kinds of tools that are shown in the bottom legend.

**Supplementary Figure 2.**
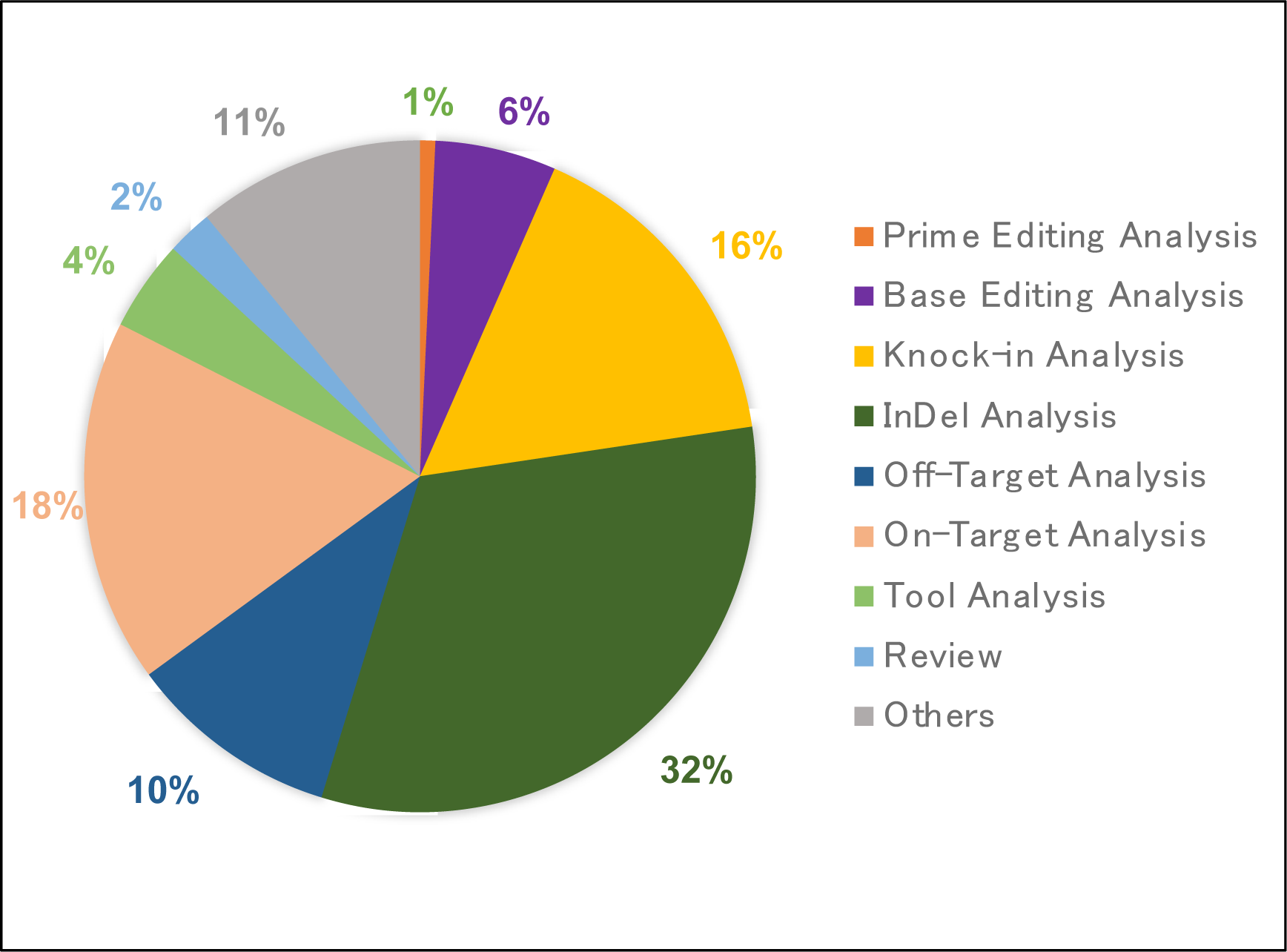
Type of publications among papers citing CRISPResso or CRISPResso2 (n=137, 2016–2020). The figure shows the percentages of papers citing CRISPResso or CRISPResso2 per the type of usage. “Prime Editing Analysis” and “Base Editing Analysis” includes the research articles or protocols that analyzed the variants generated by Prime Editing^5^ and Base Editing^9^, respectively. “Knock-in Analysis” includes the research articles or protocols that analyzed the insertion variants generated by the donor-mediated knock-in systems. “InDel Analysis” includes the research articles or protocols that analyzed the InDel patterns and showed the figure(s) of the read alignment or the InDel distribution in the context of knock-out study. “Off- Target Analysis” includes the research articles or protocols that analyzed the off-target regions. “On-Target Analysis” includes the research articles or protocols that analyzed the InDel frequency or checked the mosaicism. “Review” indicates a review article. “Tool Analysis” includes the research articles comparing a novel tool with CRISPResso or CRISPResso2. “Others” includes the research articles that did not use CRISPResso or CRISPResso2 but mentioned them. When the papers are applicable in two or more classifications, it was classified into one of the nine types according to the following priority: Prime Editing Analysis→Base Editing Analysis→Knock-in Analysis→InDel Analysis→Off-Target Analysis→On-Target Analysis→Tool Analysis→Review→Others.

**Supplementary Figure 3.**
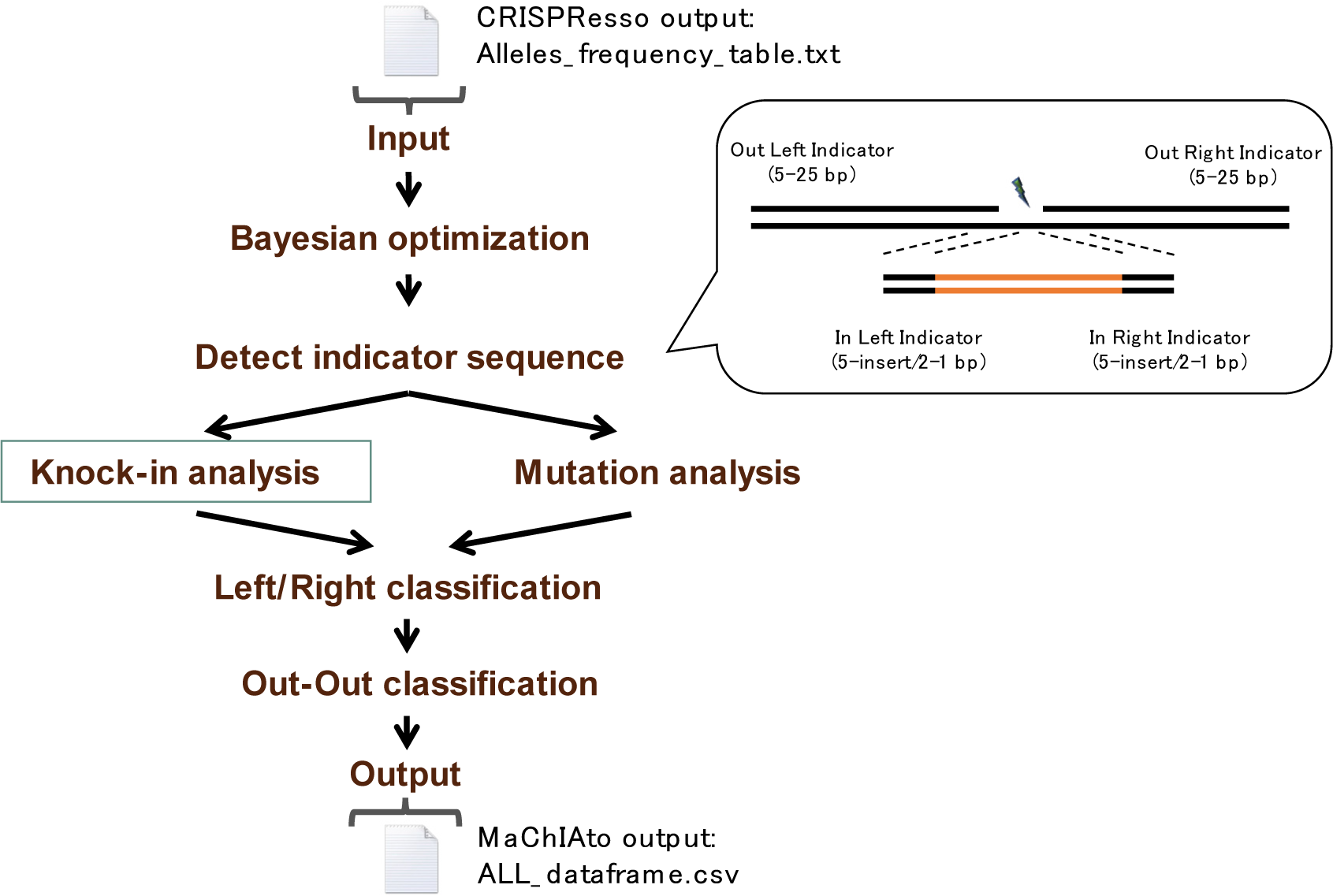
The schematic overview showing a process in MaChIAto Classifier to re-classify data from CRISPResso series. At first, MaChIAto Classifier performs Bayesian optimization using the data of “Alleles_frequency_table.txt” to adjust parameters. Next, MaChIAto Classifier confirms the positional relationships by detecting indicator sequences. After that, MaChIAto Classifier checks whether the reads are knock-in reads or not. The left/right junctions of knock-in reads are analyzed in Knock-in analysis, whereas the cut site of non-knock-in reads is validated in Mutation analysis. After Left/Right classification and Out-Out classification, the reads are re-classified based on the result of the total process.

To do that, the lengths of the Left out indicator, the Right out indicator, the left junction sequence, and the right junction sequence are changed repeatedly unless the minimum value of the following acquisition function is found.

The acquisition function is f [g (RDVHDR, RDVMixed, RDVNHEJ, RDVUnmodified)]

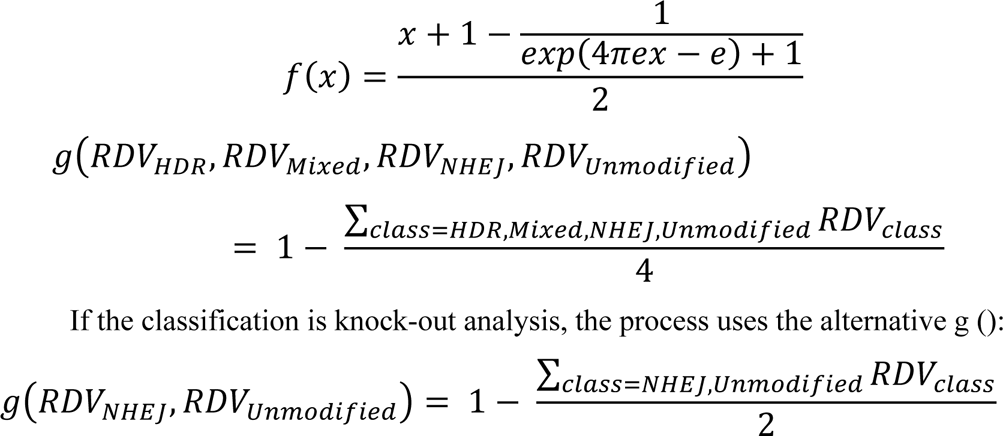

The purpose of the “out-out alignment thresholds” step is to adjust the alignment thresholds for the alignment between the Left out indicator and the Right out indicator. It is also called out-out alignment, so that it can distinguish unmodified and HDR reads from others. The process uses the **r**ate of **v**ariants which gets “is_**u**ntreated” flag (RVU) and “is_**p**erfect_outout_ki” flag (RVP) in some classes of CRISPResso: RVU NHEJ, RVUUnmodified, RVPHDR, and RVPMixed.

To do that, the threshold for left bit-by-bit alignment (Supplementary Note 2) in out-out alignment, and the threshold for right bit-by-bit alignment in out-out alignment are changed repeatedly unless the minimum value of the following acquisition function is found.

The used acquisition function is f

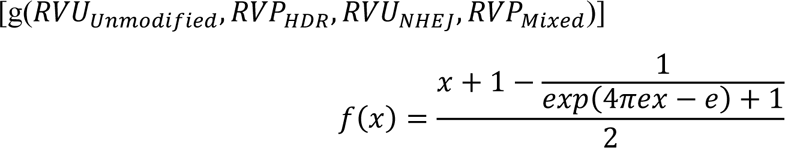

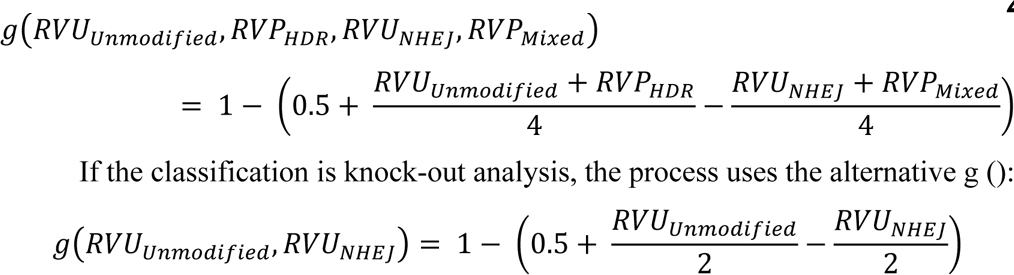

The purpose of “optimizing in-out indicators” is to adjust the indicators so that the only variants in HDR and Mixed HDR-NHEJ classes can be identified as knock-in variants. The process uses the **a**verage of **v**ariants which “has_**l**eftinout_indicators” flag and “has_**r**ightinout_**i**ndicators” flag (AVLRI) in some classes of CRISPResso: AVLRINHEJ, AVLRIHDR, and AVLRIMixed.

To do that, length of left in indicator and length of right in indicator are changed repeatedly unless the minimum value of the following acquisition function is found.

The used acquisition function is f [g (AVLRINHEJ, AVLRIHDR, AVLRIMixed)]

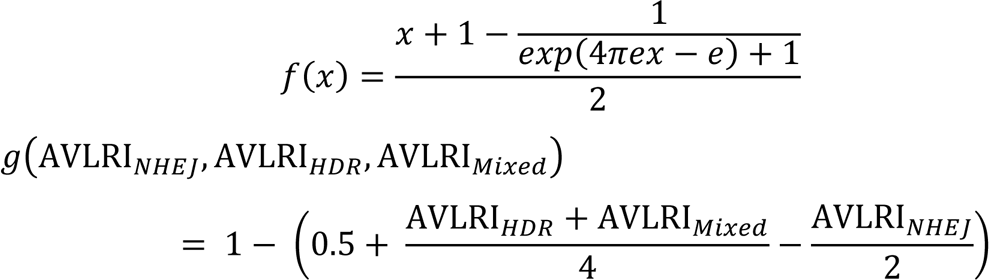

If the classification is knock-out analysis, the parameters are {(the length of donor)/2 – 1, (the length of donor)/2 – 1}. An optimization process is not performed.

The purpose of “optimizing in-out alignment thresholds” is to adjust the alignment thresholds for the alignment between “left/right out indicator” and “left/right in indicator” (left/right junction alignment), so that the only variants in the HDR class can be identified as precise knock-in variants. The process uses the **r**ate of **v**ariants which gets “is_**n**early_**l**eftinout_ki” flag and “is_nearly_**r**ightinout_ki” flag (RVNLR) in some classes of CRISPResso: RVNLRNHEJ, RVNLRHDR, and RVNLRMixed.

To do that, threshold for right bit-by-bit alignment in left junction alignment, and threshold for left bit-by-bit alignment in left junction alignment are changed repeatedly unless the minimum value of the following acquisition function is found.

The used acquisition function is f [g(RVNLRNHEJ, RVNLRHDR, RVNLRMixed)]

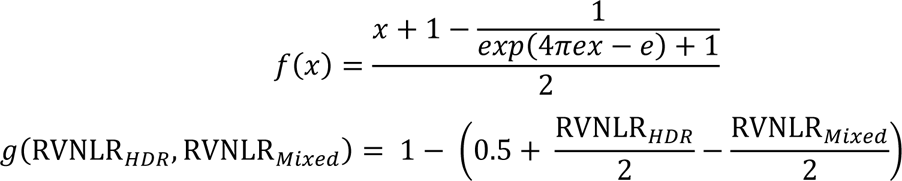

If the classification is knock-out analysis, the parameters are {1, 1}. An optimization process is not performed.

After all the optimization processes, MaChIAto Classifier applies knock-in analysis and mutation analysis to each “Aligned_Sequence” of the “Alleles_frequency_table.txt.” Through this process, MaChIAto Classifier checks 33 items for “Aligned_Sequence” and records 18 kinds of value used in alignment, and then MaChIAto Classifier extracts five sequences caught between indicators from “Aligned_Sequence” to use in the following alignment analysis.

Next, MaChIAto Classifier performs left/right classification for evaluation of the knock-in junction and out-out classification for evaluation of on-target and precise knock-in. The table “Alleles_frequency_table.txt” is given labels of “global_left_classification_label,” “global_right_classification_label” and “global_outout_classification_label.”

Finally, the table is given “CRISPResso_reclassification_labels,” which represent the result of reclassification. The name is the same as the original CRISPResso classification, in which the definition of NHEJ, Mixed HDR-NHEJ, and HDR classes mean exact InDels except for knock-in, imprecise knock-in, and precise knock-in, respectively. Furthermore, MaChIAto provides a unique class, named Unclassified (UC), which is given to a sequence that does not have Out-Right/Left indicators pair and a non-knock-in sequence with >30 bp InDels in the either of both sides. The previous report confirmed that ≤ 30 bp InDel represents high reproducibility in various genomic loci^12^. Therefore, our analysis focuses on InDel distribution up to 30 bp to help users investigate the preference for CRISPR edits. The table with all variants is saved as “ALL_dataframe.csv,” and the partial table is saved as “CRISPResso_<CLASS_NAME>_dataframe.csv” per class of CRISPResso classification (Supplementary Table 1). Moreover, the MaChIAto classification is plotted as pie charts, and then the figures are saved as MaChIAto_alignment_pie_chart.png and .pdf file, and MaChIAto_alignment_pie_chart [without UC(Other)].png and .pdf file In these figures, Ambiguous Editing label is separated from other UC reads. Ambiguous Editing read has the complete donor sequence and homology arms but does not have the Out-Right/Left indicator pair. Therefore, it is not confirmed whether the Ambiguous Editing read is a perfect editing sequence or an imperfect one. Such a read might be a chimeric read.

MaChIAto Analyzer can be used for the dependency analysis between > 70 main genomic features on homology arms and the summary of re-classification such as precise knock-in efficiency and accuracy. The tool requires a summary directory of all results from MaChIAto Classifier, and extra tables (CSV format) of name and pathname. The summary directory can be generated using “collect_MaChIAto_data.py” based on the output of MaChIAto Classifier, and it contains the summary of re- classification, sequence information per target locus and sample type. “collect_MaChIAto_data.py” requires the choice of calculation targets {“Leftside homology arm (lmh),” “Rightside homology arm (rmh),” “Bothside homology arm (bmh),” “Leftside edited homology arm (elmh),” “Rightside edited homology arm (ermh),” “Bothside edited homology arm (ebmh),” “protospacer”}, and optionally imports the list of an ignored target set (CSV file) that includes targets that users want to ignore in MaChIAto Analyzer. The script also requires labels that indicate sample type {“untreated,” “knock-out,” “knock-in”}, and then recognizes the analysis type of MaChIAto Analyzer {“Double knock-in analysis,” “Single knock-in analysis,” “Simple knock-in analysis,” “Double knock-out analysis,” “Single knock-out analysis”}.

Most main genomic features are defined in previous reports. For example, frequencies of mono and dinucleotides (Freq), DNA secondary structures based on dinucleotide and tetranucleotide properties (Packer), physiochemical properties (Phychem), pseudo K-tuple nucleotide composition (PseKNC), and thermodynamics and secondary structure properties (Thermo) were used in the previous studies on sgRNA activity^13^. In MaChIAto Analyzer, the values are calculated based on the sequences of the calculation target. The process is performed using the following R script: “CalcNNucFreq-class.R,” “CalcPacker-class.R,” “CalcPhyChem-class.R,” “CalcPseKNC-class.R,” and “CalcThermo-class.R.”

“MP” features, which is one of the main genomic features, are previously defined microhomology score features^19^. They include microhomology scores at knock-in junctions. These are calculated using CalcMHScore-class.R, which is a wrapper program of the python script used in a previous report^14^.

“SgRNA.Align” features, which are also one of the main genomic features, are the features of the alignment score between the target site and sgRNA including the scaffold sequence. The alignment is performed between the 3′ end of the left homology arm and the 5′ end of the *k* nt sgRNA sequence, or the 5′ end of right homology arm and the 3′ end of *k* nt sgRNA sequence. If the calculation target is “protospacer,” 3′ end of the protospacer and the 5′ end of the *k* nt sgRNA sequence. The *k* can increase from 20 nt to the maximum length of sgRNA sequence per 20 nt.

Furthermore, users can freely add the extra (epi-)genomic features for the dependency analysis by writing that into the extra tables.

We showed the example of the extra (epi-)genomic feature tables (“Genome.Property,” “inDelphi,” “Accessibility”) that were used in a series of knock-in targeting 40 gene loci (Supplementary Note 4).

“Genome.Property” feature table contains “Chromosome_size,” “ExonNumber,” “protein_length.aa.,” “Position.relative.,” “CNV_HMM_293T,” “Expression,” and “Editing.Efficiency.”

“Chromosome_size” is the whole size of the chromosome to which the target belongs. It is taken from the hg19 genome assembly.

“ExonNumber” is the number of an exon to which the target belongs. It is taken from the UCSC Genome Browser^15^ and a previous report regarding a CRISPR KO screen^16^.

“protein_length.aa.” and “Position.relative.” are the length (amino acids) and the relative position of the target amino acid to the whole length of the protein encoded by the target, respectively. It is taken from the UCSC Genome Browser^15^.

“CNV_HMM_293T”^17^ is the value of the Complete Genomics CNV by the HMM algorithm. The value is taken from the Integrative Genome Viewer version- 2.3.9716^18^.

“Expression” is RNA-seq data of the target gene. It is taken from GTEx Analysis V6p. The original data is “Gene read count” (GTEx_Analysis_v6p_RNA- seq_RNA-SeQCv1.1.8_gene_reads.gct.gz). The data of the tissue to which the sample cells belong is extracted from the read count data, and then the values are normalized using the trimmed mean of M values (TMM)^19^ to generate the value of “Expression.”

“Editing.Efficiency” is the value of non-knock-in (knock-out) sample calculated by MaChIAto Classifier. This feature is automatically retrieved by

MaChIAto Analyzer if the analysis type is “Double knock-in analysis” or “Single knock-in analysis.” “inDelphi” is the feature of InDel tendency predicted using a machine learning algorithm^20^. It is taken from the inDelphi website (https://indelphi.giffordlab.mit.edu). Left or Right Accessibility is the accessibility in which a microhomology sequence and the flanking sequence lie. “Left” and “Right” indicate the extending direction. When there is no “Left” and “Right” in the name, the extending direction is both directions. It is taken from “Base Overlap Signal” in “Duke DNaseI HS,” ENCODE^21^ (http://hgsv.washington.edu/cgi-bin/hgTrackUi?g=wgEncodeOpenChromDnase). The calculation is performed using Blat^36^ and homemade scripts. The detailed script is available on our GitHub page (https://machiatopage.github.io/2022/06/25/How-to-calculate-the-value-of-accessibility/#more).

MaChIAto Analyzer first performed several filtration steps to exclude the data of failed experiments. First, the targeting loci with the low number reads (< 1,000) are removed to guarantee resolution of outcome; second, the targeting loci with the low number of the classified reads (< 1,000) are removed to guarantee quality of classification; third, the targeting loci with the low rate unmodified reads in the untreated sample (the rate of unmodified reads < 80%) are removed to avoid noise from unusual sequencing results such as a high rate of amplification-error and cross-contamination; fourth, the targeting loci where the untreated sample and the knock- out_1 sample represent no significant difference (chi-squared test, *p*-value ≥ 0.001), are removed because the result does not make sense in the context of genome editing if the analysis type is not the Simple knock-in analysis; fifth, in the case of the Double knock-in analysis, the targeting loci, where the knock-in_1 sample or the knock-in_2 sample have no precise knock-in, are removed because the result does not make sense in the context of knock-in experiments. In the case of the Double knock-out analysis, the locus, where the knock-out_1 sample or the knock-out_2 sample has no InDel, is also removed. After the filtering process, MaChIAto Analyzer calculates Pearson correlation between the knock-in result and the features of target loci that have all features, and MaChIAto Analyzer also estimates the correlation between enhancement from condition 1 (knock-in_1 / knock-out_1) to condition 2 (knock-in_1 / knock-out_1) in the raw scale or the log scale. Finally, MaChIAto Analyzer summarizes the correlation value as a box plot and then performs a paired *t*-test and a Kolmogorov-Smirnov test between condition 1 to condition 2 (Fig. 1c). If the sequence of calculation target repeats in the sample set, the tool calculates average values per the same sequence, and then uses them as the representative value in the above correlation analysis.

MaChIAto Aligner creates sequence alignment per the class of CRISPResso and MaChIAto, and also visualizes the distribution of InDel size and the ratio of NHEJ/MMEJ. The tool requires the output of MaChIAto Classifier and optionally left and right extra sequences as input. The left and right extra sequences are fragments flanking homology arms in the knock-in donor, which result from CRISPR cutting. The example of left and right extra sequences is described in the online manual. Initially, MaChIAto Aligner reads sequence information on the output. Next, the “OutOut_Aligned_Sequence,” “Leftside_InOut_Aligned_Sequence,” “Rightside_InOut_Aligned_Sequence,” “RC_Leftside_InOut_Aligned_Sequence” and “RC_Rightside_InOut_Aligned_Sequence” of “ALL_dataframe.csv” are loaded as regions of the “cut site,” “left junction (in the case of knock-in with correct direction),” “right junction (in the case of knock-in with correct direction),” “left junction (in the case of knock-in with reverse-complement direction)” and “right junction (in the case of knock-in with reverse-complement direction),” respectively. These five sequences are aligned using BWA MEM^22^ and CrispRVariants^3^, when unmodified and the expected knock-in sequence is used as a reference sequence. After that, MaChIAto Aligner performs the distribution analysis of InDel size and position, and also performs MMEJ recognition based on the variant calling of CrispRVariants. The alignment result is saved as “aligned.variants.pdf.” Subsequently, the quantification and visualization are carried out using merged data of a variants list which is made of output “variantCounts” and “consensusAlleles” function of CrispRVariants. The variants list, named “variants.count.total.list.rds,” is saved with “aligned.variants.pdf” in each directory of output. The variants list contains the information on MMEJ/NHEJ, microhomology and intervening sequence. The detailed structure of the variants list and MMEJ detection method is described in the online manual.

MaChIAto Reviewer requires the output of MaChIAto Classifier and MaChIAto Aligner, and the summary directory used in MaChIAto Analyzer as input. Initially, MaChIAto Reviewer executes the same filtering process as MaChIAto Analyzer described above. Next, MaChIAto Reviewer evaluates the difference between CRISPResso series and MaChIAto (Fig. 1b and Supplementary Fig. 2), and also visualizes and summarizes the efficiency such as “Precise Knock-in Efficiency” and “Indels Editing Efficiency,” and the level of the enhancement. MaChIAto Reviewer makes three groups, called “High,” “Medium,” and “Low,” according to the efficiency and enhancement levels so that the members can be split into groups of almost equal size (Fig. 1e, top). MaChIAto Reviewer calculates the cumulative ratio of outputs of MaChIAto Aligner by the groups, and then the ratio is plotted as an overlapping bar or multiple bar plot. MaChIAto Reviewer also makes a pie chart of max and total frequent categories regarding InDel size and MMEJ/NHEJ. The visualization is carried out in a group set of the efficiency of condition 1, the efficiency of condition 2, and the enhancement level from condition 1 to condition 2.

### Plasmid construction and oligonucleotide preparation

The all-in-one CRISPR-Cas9 vectors expressing Cas9 nuclease and multiple sgRNAs were constructed using the Multiplex CRISPR/Cas9 Assembly System Kit (Kit #1000000055, Addgene), as described previously^23^ with some modifications. For the addition of MS2 stem loops to the sgRNA scaffold, the sgRNA expression cassette of the pX330A and pX330S vectors was replaced with that of the sgRNA (MS2) cloning backbone vector (Plasmid #61424, Addgene)^24^ as previously described^8^. The oligonucleotides used for the sgRNA template are listed in Supplementary Table 2. The CtIP effector vectors expressing the MS2 coat protein were described previously^8^. The PITCh donor vectors carrying the 3× Flag tag and microhomologies were constructed by PCR and Gibson Assembly using an In-Fusion® HD Cloning Kit (TaKaRa) with the primer sets in Supplementary Table 2. PCR and TA-Cloning were performed to add the target sequences of universal sgRNA (previously described as PITCh-gRNA #1)^7^ using TArget Clone -Plus- (TOYOBO). The primers used are listed in Supplementary Table 2.

### Cell culture and transfection

HEK293T cells were maintained in Dulbecco’s modified Eagle’s medium (DMEM) supplemented with 10% fetal bovine serum (FBS). Lipofectamine LTX (Life Technologies) and serum-free DMEM were used to transfect cells with plasmids according to the supplier’s protocol. For the genomic PCR and NGS analysis of all loci in the cells knocked-in with 3× Flag tag, 200 ng of each plasmid (PITCh donor vector, all-in-one CRISPR-Cas9 vector, and MCP-CtIP vector) was transfected into 4.5 × 10^4^ cells using a 96-well plate. At 24 and 48 h post-transfection, 70 µl of DMEM supplemented with 10% FBS was added to each well. There was no drug or fluorescence selection applied during cell culture.

### Genomic DNA extraction, PCR, Amplicon sequencing and data analysis

Transfected cells were collected using TrypLE Express (GIBCO) and PBS (-) at 72 h post-transfection. Genomic DNA was extracted from the collected cells using a Wizard® SV 96 Genomic DNA Purification System (Promega). Genomic PCR was performed using PrimeSTAR Max DNA Polymerase (TaKaRa). Preparation of libraries and NGS were performed by FASMAC. Briefly, NGS libraries were prepared via three-step or two-step PCR using the primers listed in Supplementary Table 2. PCR amplification was performed using Ex Taq Hot Start Version (TaKaRa) or KAPA HiFi HS ReadyMix (KARABIOSYSTEMS), and the PCR products were purified using Agencourt AMPure XP (Beckman Coulter). Note that the PCR products of *C8orf86*, *DAAM1*, *FOXF1*, *RAPGEF3*, and *WFDC2* loci included bands of unexpected size; in these cases, the correct bands were excised from the gels and DNAs were purified using a FastGene Gel/PCR Extraction Kit (NIPPON Genetics). Amplicon sequencing was performed using a MiSeq sequencer (Illumina). The genomic PCR products were also analyzed by microchip electrophoresis using MultiNA (Shimadzu) and LabChip GX Touch HT (Perkin Elmer) to confirm the edited bands.

The NGS data were analyzed using CRISPResso2^2^ and MaChIAto. The parameters used for these analyses are described in Supplementary Table 4. In a series of knock-in targeting several gene loci (Supplementary Note 4), 50 genes were filtered to 40 genes because the gene targeting data of bad quality were removed by MaChIAto.

#### Supplementary Note 1 - Usage of analysis tools for amplicon sequencing in the context of genome editing

Many computational tools have been developed to evaluate the amplicon sequencing data which come from genome-editing experiments. For example, CRISPR-GA^25^, AGEseq^26^, OutKnocker^27^, ampliconDIVider^28^, BATCH-GE^29^, CRISPResso^1^, CrispRVariants^3^, Cas-analyzer^11^, CRISPR-DAV^30^, CRISPRMatch^31^, ampliCan^4^, CRIS.py^32^, CRISPResso2^2^, and Hi-TOM^33^ have been used. We counted the number of the papers cited each tool per year as of 24th May 2020 (Supplementary Table 3 and Supplementary Fig. 1).

The total number of publications citing these tools has reached 428 papers within less than seven years, which implies the high importance of these softwares for the advancing field of genome editing. Notably, CRISPResso has recently become one of the most cited tools, although it was published later than the pioneer tools such as ampliconDIVider. CRISPRessso offers a wide range of usages in genome editing. We summarized the usages of CRISPResso in previous studies (Supplementary Table 3 and Supplementary Fig. 2). The percentages of “Knock-In Analysis” and “InDel Analysis” occupied approximately half of all usages (48%). We suppose that this result shows the researchers’ demand for the amplicon sequencing has focused on a more in-depth and complex analysis of amplicon reads such as the identification of end-joining and accurate classification of the integration reads.

#### Supplementary Note 2 - Details of the MaChIAto Classifier pipeline

MaChIAto receives “Alleles_frequency_table.txt” as input and the data are processed through the following pipeline (Supplementary Fig. 3). MaChIAto Classifier applies novel 5 techniques: 1) Bayesian optimization for the optimal parameters, 2) multiple indicators, 3) expansion alignment, 4) reverse knock-in detection using indicator, and 5) “degeneracy” classification with “strict” labeling. Detailed explanations of each technique are described below along with the motivations for the implementation.

1. The parameter adjustment is essential for the analysis of NGS reads because amplicon reads can have a lot of PCR errors and/or sequence errors which depend on the sample condition and sequence context. Most tools accommodate the adjustable parameter to get *bona fide* mutations by eliminating the noise. However, in most cases, the process is conducted manually and there is no standard. Therefore, the parameter adjustment is often a time-consuming and unfair analysis. MaChIAto uses Bayesian optimization to manage this problem. Bayesian optimization is one of the most famous parameter optimization methods, performing an automatic trial-and-error procedure. The method repeats a partial process of MaChIAto Classifier, while changing parameters. The results are recorded in the system and the system estimates new parameters which are expected to get a better result. Subsequently, the system decides the best parameters based on the records. This method can perform an adjustment of parameters as a person does manually, but it is faster because the internal program is designed to perform the method efficiently. Moreover, the parameter is fair because it is based on the calculable rules. In Bayesian optimization of MaChIAto, the parameters include “length of left junction,” “length of right junction,” “length of Out Left indicator,” “length of Out Right indicator,” “length of In Left indicator,” “length of In Right indicator,” “threshold of out- left alignment,” “threshold of out-right alignment,” “threshold of in-left alignment,” and “threshold of in-right alignment.” The process of optimization is split into four steps and each step is different in adjustable parameters and calculation. The detailed information on the steps is described in the section of Methods.
2. The amplicon sequencing analysis tools^1–4, 11, 25–33^ for genome editing apply various methods to identify a mutation in amplicon reads. Some of the existing tools^1, 2, 4^ use Needleman-Wunsch alignment for the identification. This reasonable method, however, can fail to identify the positional relationships which are important for the classification between knock-in and non-knock-in. This problem may be caused by the alignment score, which does not have the information on the positional relationships such as the distance between 5′ end of knock-in junction and donor. On the other hand, other tools^3^ use mapping software such as BWA MEM for the identification. Such an aligner is expected to achieve accurate variants identification, but it can miss large InDels^4^. Therefore, it is not appropriate for the first classification of amplicon reads because the loss of reads should be minimized in this step. MaChIAto Classifier offers an advanced “indicator” method for the first classification. The original indicator method is used in Cas-Analyzer^11^. The method uses a part of sequences flanking the cut site to perform character matching with amplicon reads. The positional relationships can be clear using this tool. In MaChIAto Classifier, four indicators are used for more accurate recognition of on-target sequence and knock-in sequence to recognize the relationships with higher resolution. These indicators are located around the cutting site and on knock-in donor sequence (Supplementary Fig. 3, callout). Our tool can perform the first classification using these indicators and the secondary analysis pipelines such as “Knock-in analysis” and “Mutation analysis” per amplicon read based on the rough classification. After that, MaChIAto reliably utilizes BWA MEM to get the more detailed variants profile (MaChIAto Aligner).
3. Needleman-Wunsch alignment is reasonable to detect mutations. However, when the alignment is used against the whole sequence once, it is difficult to distinguish *bona fide* mutation by genome editing from non-editing mutation caused by PCR error. To solve this problem, MaChIAto Classifier performs expansion alignment in the phase of Knock-in analysis and Mutation analysis. The method is named “Bit-by-Bit Alignment.” Bit-by-Bit Alignment is a multi-step method of Needleman-Wunsch alignment and identifies *bona fide* editing reads by detecting a mutational hotspot that is truly caused by genome editing. Bit-by-Bit Alignment detects a hotspot of mutation with the following four steps: i) the initial alignment starts at both Out Left/Right Indicators in amplicon reads. ii) The range of alignment expands in the direction of the center of the sequence. There may be two stories (a, b) after this. iii-a) When there is a single substitution, the alignment score goes down. The algorithm continues the alignment unless the score is below the alignment threshold. For example, If the small substitution only exists, it will be ignored. iii-b) When it comes across the hotspot of mutation, the alignment score goes sharply down below the alignment threshold, and the algorithm decides that the corresponding mutation is the hotspot. The algorithm ends in this case. iv-a) When both alignment ranges overlap each other, the algorithm reports that there is no mutation spot.
4. Most published tools cannot recognize reverse knock-in, in which a donor insert is integrated in a reverse-compliment direction at the cut site. MaChIAto Classifier can detect reverse knock-in using the indicators. The sequences of “Out left indicator,” “Out right indicator,” the reverse complement of “In left indicator,” and the reverse complement of “In right indicator” are used for it. The detected junction sequence of reverse knock-in is aligned using the reference sequence of potential NHEJ-assisted reverse knock-in sequence to classify. After the classification process, the ratio between correct direction knock-in and reverse direction knock-in is calculated as a value of “reversibility.”
5. The classification tools for amplicon reads generally define one standard rule for distinguishing a class from others. For example, CRISPResso^1^ identifies knock-in read when a score which comes from the alignment between a read and the expected knock-in sequence reaches the setting threshold (hdr_perfect_alignment_threshold). However, even when the alignment score is lower than the threshold, the length can be the same as that of precise knock- in, and both junction sequences are similar to that of the reference. In this case, the read is likely to be a precise knock-in read. In addition, the low alignment score is due to PCR error on the part of donor sequence which is not involved in the DSB repair between the genomic cut site and donor. MaChIAto can accept the two patterns as a precise knock-in sequence. The former pattern, which is similar to the whole precise knock-in sequence, is labeled as “left/right_perfect_ki,” whereas the other pattern, which has the same length of precise knock-in and the similar sequence to the expected left/right junction of the precise knock-in, is added the label of “left/right_nearly_ki.” The read which has “left/right_perfect_ki” or “left/right_nearly_ki” is counted into “PRECISE_KNOCK_IN” class. MaChIAto Classifier also accommodates the “degeneracy” classification in other processes such as the detection of imprecise knock-in using the “strict” labels. The strategy enables not only highly flexible and efficient classification but also more informative profiling: i.e., the output files provide the labeling record so that a user can review how a read is classified into the class. In the results of most classification tools, the decision process of the amplicon reads is not specified, which makes it a black box. However, an analysis of MaChIAto Classifier has high transparency and offers the data which can be used for an additional analysis by the user.

#### Supplementary Note 3 - The evaluation of mutation detection using 240 simulation data

We evaluated the accuracy of the tool by comparing it with exiting tools, namely CrispRVariants^3^, CRISPResso^1^, CRISPResso2^2^, CRISPRessoPooled^1^ and AmpliconDIVider^28^. For this comparison, we utilized large-scale amplicon data (N=240) known as “Synthetic dataset 2.”^3^ This dataset had been artificially generated using the NGS simulation software, ART^34^, and contained on-target and off-target reads at certain rates. We reproduced the data of CrispRVariants, CRISPResso, CRISPRessoPooled and AmpliconDIVider according to the instruction at https://github.com/markrobinsonuzh/CrispRvariants_manuscript. However, some commands of “ART” and “SAMTOOLs”^35^ were changed to the usage of the recent version so that the output can remain that of the previous version. The PAM sequences were deleted from the sgRNA target sequence used for the input of CRISPResso and CRISPRessoPooled in the previous simulation, because the original manual of CRISPResso^1^ defined that the input of sgRNA is the guide RNA sequence (usually 20 nt) adjacent to 5′ of a PAM sequence (e.g., NGG for SpCas9). The simulation script for CRISPResso2 is written based on that for CRISPResso: i.e., the specification of a window (-w) is 5 bp and the other settings were the default value. The outputs of CRISPResso2 were used for the input of MaChIAto. The setting of MaChIAto is the default value for the input of CRISPResso2. The estimated mutation efficiency of CRISPResso is the value of “Modified” in “CRISPResso_quantification_of_editing_frequency.txt,” and the estimated mutation efficiency of MaChIAto (CRISPResso2-MaChIAto) is the following value:

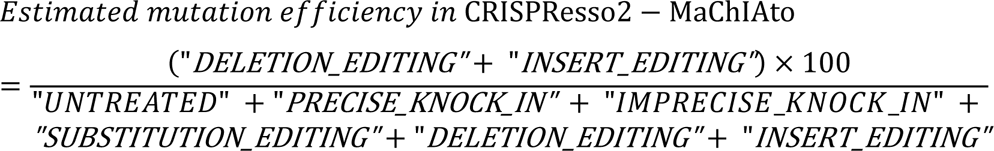

Each value is read count which is shown in “ALL_read_countdata.csv” in the output of MaChIAto.

We made a figure similar to Figure 10 in Supplementary Note 4 in a previous report^3^, which illustrated the performance of tools on “Synthetic dataset 2” with true mutation efficiency of 0%, 33.3%, 66.6% or 90% (Supplementary Fig. 4). The CRISPResso (light green) and its derivative tool, CRISPRessoPooled, (light purple) had a low performance for distinguishing true mutations from sequencing errors or off-target mutations (false mutations) (Supplementary Fig. 4, top right and bottom). CRISPResso2 (purple), however, had a higher performance of identification (Supplementary Fig. 4, bottom right), although it overestimated the mutation efficiency in the zero-mutation condition because it counted false mutations as true mutations (top left, Supplementary Fig. 4). CRISPResso2-MaChIAto (orange) succeeded in correcting the original classification of CRISPResso2 and was almost equal to CrispRVariants (light blue) and AmpliconDIVider (light orange), which achieved the top performance in this reproduction data. The most notable thing is that MaChIAto did not use mapping software in this classification. Of course, the full MaChIAto pipeline uses mapping software: BWA MEM for variant identification in MaChIAto Aligner. However, the shown classification data was pre-processing data from MaChIAto Aligner, which was the output of MaChIAto Classifier. This means that the classification performance of MaChIAto in variant classification is not dependent on a high-quality mapping tool, but that MaChIAto can reach the same performance as a tool that uses mapping. Moreover, MaChIAto does not require an additional control sample or the whole genomic sequence if only the classification function is used. Therefore, the requirement of MaChIAto is almost the same as that of CRISPResso2, which is known as a user- friendly tool.

**Supplementary Figure 4.**
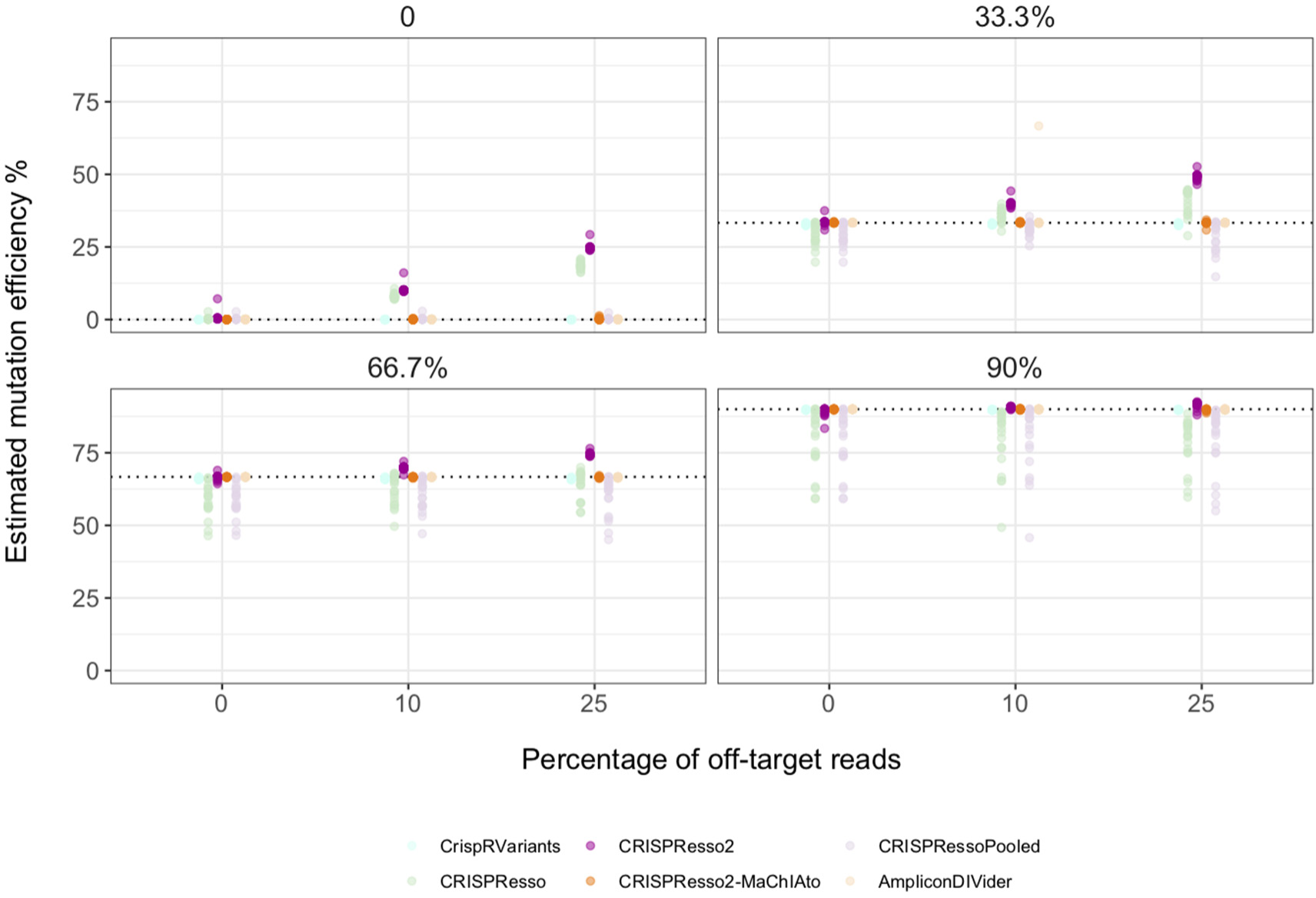
The performance comparison of various classification tools for amplicon sequencing reads. Each point indicates the estimated mutation efficiency for a single simulation. The simulation set uses synthetic amplicon data of varied proportions between unmutated sequences and ten mutation sequences. The y-axis indicates the estimated mutation efficiency as a percentage. The x-axis indicates the rate of off-target reads in the simulation data. The true mutation efficiency of the synthetic data is displayed on the top of each plot, and the horizontal dotted line on the y-axis corresponds to the percentage of that. CRISPResso and CRISPRessoPooled results were better than the simulation result in the original CrispRVariants paper^3^ because the setting of sgRNA was corrected.

#### Supplementary Note 4: Design of a series of knock-in targeting 40 gene loci

We tested the streamlined analysis of newly obtained experimental data using MaChIAto. We had previously established the PITCh (precise integration into target chromosome) and LoAD (local accumulation of DSB repair molecules) systems, which are an MMEJ-dependent knock-in strategy and its enhancement system by recruiting CtIP proteins at the CRISPR target sites, respectively^6–8^. We here show an overview of the process for a series of PITCh and LoADed PITCh knock-ins targeting 40 gene loci (Supplementary Fig. 5). The process has four main steps.

1. The choice of targeted gene was based on a previously described CRISPR knock- out screening^16^. At first, we chose non-essential genes because a lethal effect might cause bias to the knock-in efficiency. Next, we chose sgRNAs that do not have < 5 mismatch sites in the genome and a low off-target score ≦ 400. This off-target score was calculated according to a previous report^16^: 

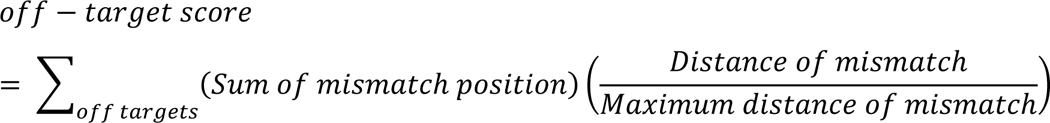

After filtering, we generated 100,000 groups containing 46 genes. We collected information regarding the distribution of DNA accessibility of microhomologies (µHs), GC content of µHs, predicted DNA accessibility, DNA methylation by program library of gkm-SVM^36^, gene expression, and chromosome number. We used the one-sample Kolmogorov-Smirnov test to check whether the distribution was unbiased. The alpha value was adopted so that the beta value could be < 0.001%. We then selected one gene set from the candidate sets. The selected gene set had wide distribution and avoided inclusion of the gene ontology (GO) annotation “biological phase” which is defined as a distinct period or stage in a biological process or cycle. Lastly, four well-documented genes, *ATP5B*, *RPL11*, *AAVS1*, and *ID1* were added to the gene set as controls.
2. The PITCh and LoAD donor vector designs including microhomology sequences were generated with PITCh designer 2.0^37^. Each CRISPR vector targets a single genomic site and two sites on the corresponding donor vector because PITCh and LoAD require linearized dsDNA donor for the targeted integration. We used 66-bp sequence fragment as a knock-in insert (Supplementary Fig. 5, callout). The insert sequence was designed without considering the change in the reading frame of the target gene. The reason why was that the integration design should be decided based on the position of PAM sequence so that the right µH should always be located in the PAM-proximal region to ensure the relationship between the PAM-proximal/distal position and the left/right µH position. There is less concern about lethal effects by integration that disrupt codons because non-essential genes had already been chosen as target loci. Therefore, theoretically, there is no concern about the survival bias caused by the integration of any donor sequence. In short, we focused on the relationship between µH and sgRNA rather than on codon context. We used a 40-bp left µH and a 40-bp right µH similar to previous studies^8^.
3. We performed knock-in experiments without any drug or fluorescence selection in HEK293T cells. After 72 h post-transfection, we performed the genomic PCR using the extracted genome samples. The NGS amplicon sequencing of the PCR product was performed using the MiSeq platform. The detailed protocol was described in Methods.
4. We analyzed the data using CRISPResso2-MaChIAto pipeline. The list of used parameters is shown in Supplementary Table 4. As a result, the 50 genes were filtered into 40 genes due because the rest of gene targeting data had bad quality. The detailed protocol was described in Methods.

**Supplementary Figure 5.**
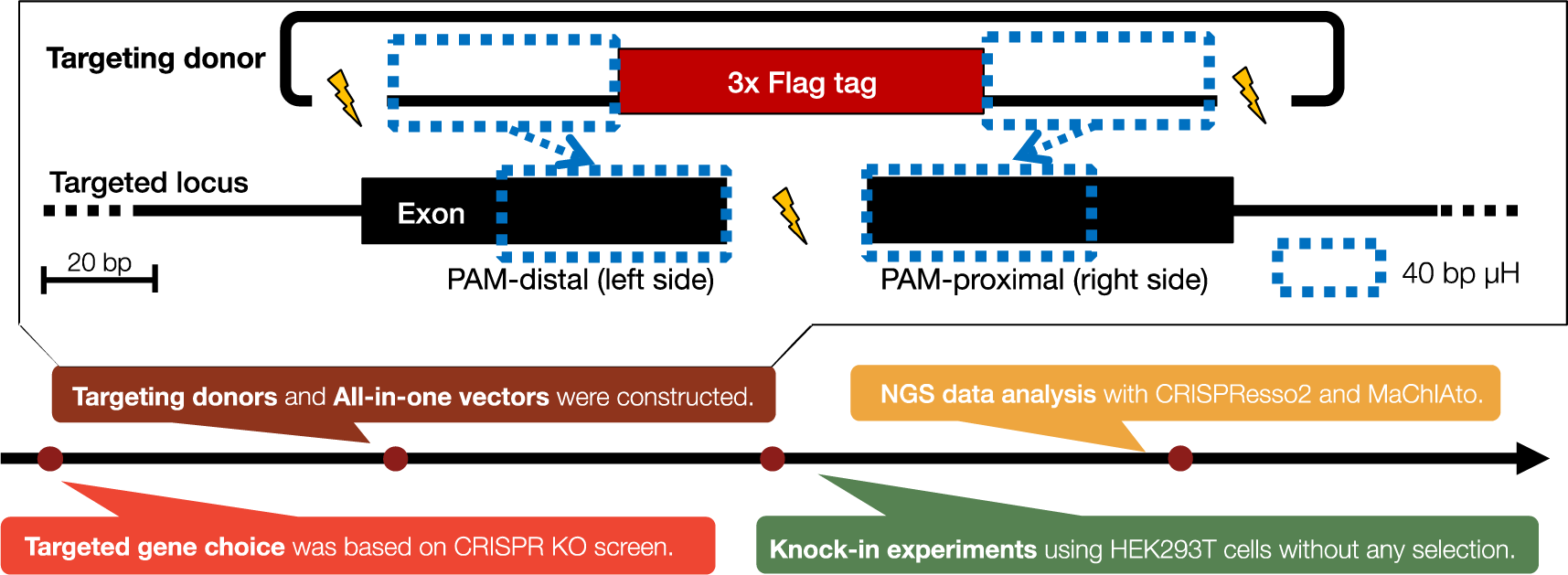
Scheme of a series of knock-in targeting 50 gene loci and design of MMEJ-assisted knock-in. The experimental process was conducted according to the black arrow. The design of PITCh and LoAD knock-in is illustrated in a white callout. The lightning bolts indicate the cut sites by Cas9-sgRNA.

#### Supplementary Note 5 - The evaluation of InDel/knock-in classification using 40 knock-in experiments data

MaChIAto could re-classify the CRISPResso2 result come from a series of knock-in targeting 40 gene loci in HEK293T cells (Fig. 1b, Supplementary Note 4). We evaluated the accuracy of classification using BWA MEM-CrispRVariants, which has a high performance regarding variant identification. First, We calculated the rate of “InDel or Knock-in” variants, with BWA MEM-CrispRVariants in the unmodified variants in CRISPResso and MaChIAto (Supplementary Fig. 6). The rate in CRISPResso2 was higher than that in MaChIAto. This result showed that CRISPResso2 tended to misclassify edited reads as unedited reads; however, MaChIAto could recover such a misclassification to build a consensus with BWA MEM–CrispRVariants, which contributes to a reduction of false negative rates in the edited reads.

**Supplementary Figure 6.**
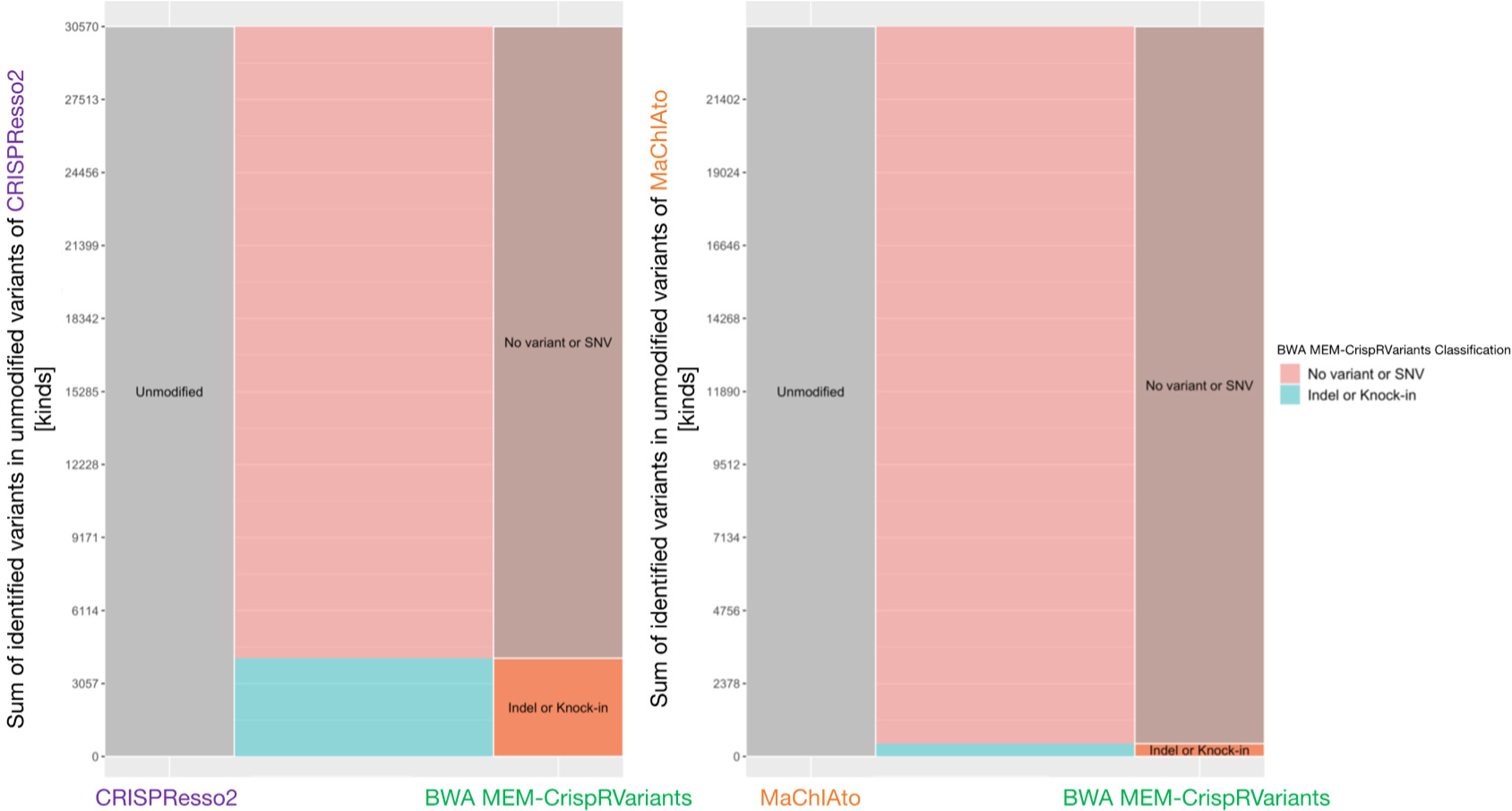
The comparison between BWA MEM-CrispRVariants and CRISPResso2 (left) or MaChIAto (right) using unmodified variants. Left parts of the left and right bar plots indicate the unmodified variants of CRISPResso2 and MaChIAto, respectively, and the y-axis is the number of identified variants in the unmodified variants. The right parts of these plots show the results of variant identification of BWA MEM-CrispRVariants using the corresponding variants. The variants are classified into “No variant or SNV” class and “InDel or knock-in” class. No variant means the unmodified sequence decided by BWA MEM-CrispRVariants. The flows indicate the change in each class. The variants with the red color flow are classified into the “No variant or SNV” class, and the variants with the blue color flow are classified into the “InDel or knock-in” class. The left plot is saved as “[h2]Alluvial_Diagrams_of_CRISPResso_Unedited_Class.png” in the output of MaChIAto Reviewer. The right plot is saved as “[i2]Alluvial_Diagrams_of_MaChIAto_Unedited_Class.png” in the output of MaChIAto Reviewer.

Finally, we focused on the difference in knock-in variants between CRISPResso2 and MaChIAto. We calculated the rate of “precise size” variants, which are insert variants having the correct length of a precise knock-in by BWA MEM– CrispRVariants in the precise knock-in variants in CRISPResso and MaChIAto (Supplementary Fig. 7). Many kinds of precise knock-in variants in CRISPResso2 do not have the precise knock-in length of the donor (66 bp). On the other hand, all of the precise knock-in variants in MaChIAto had the correct insert length. The MaChIAto algorithm recognizes the length that is the same as the sequence of the expected knock- in when the algorithm classify reads into precise knock-in class. Of course, a difference in the insert size between a knock-in read and the expected insert does not always mean the variant is an imprecise knock-in. The read might have extra InDels that are located outside the integration site. However, the rate of this more precise knock-in that MaChIAto picks up enables a discussion regarding variant identification to be clearer and more understandable. For example, severe mutations on the integrated donor can be easily searched from the output data in MaChIAto. We showed that MaChIAto can generally provide a stricter classification than CRISPResso2, thereby facilitating knock-in variant interpretation.

**Supplementary Figure 7.**
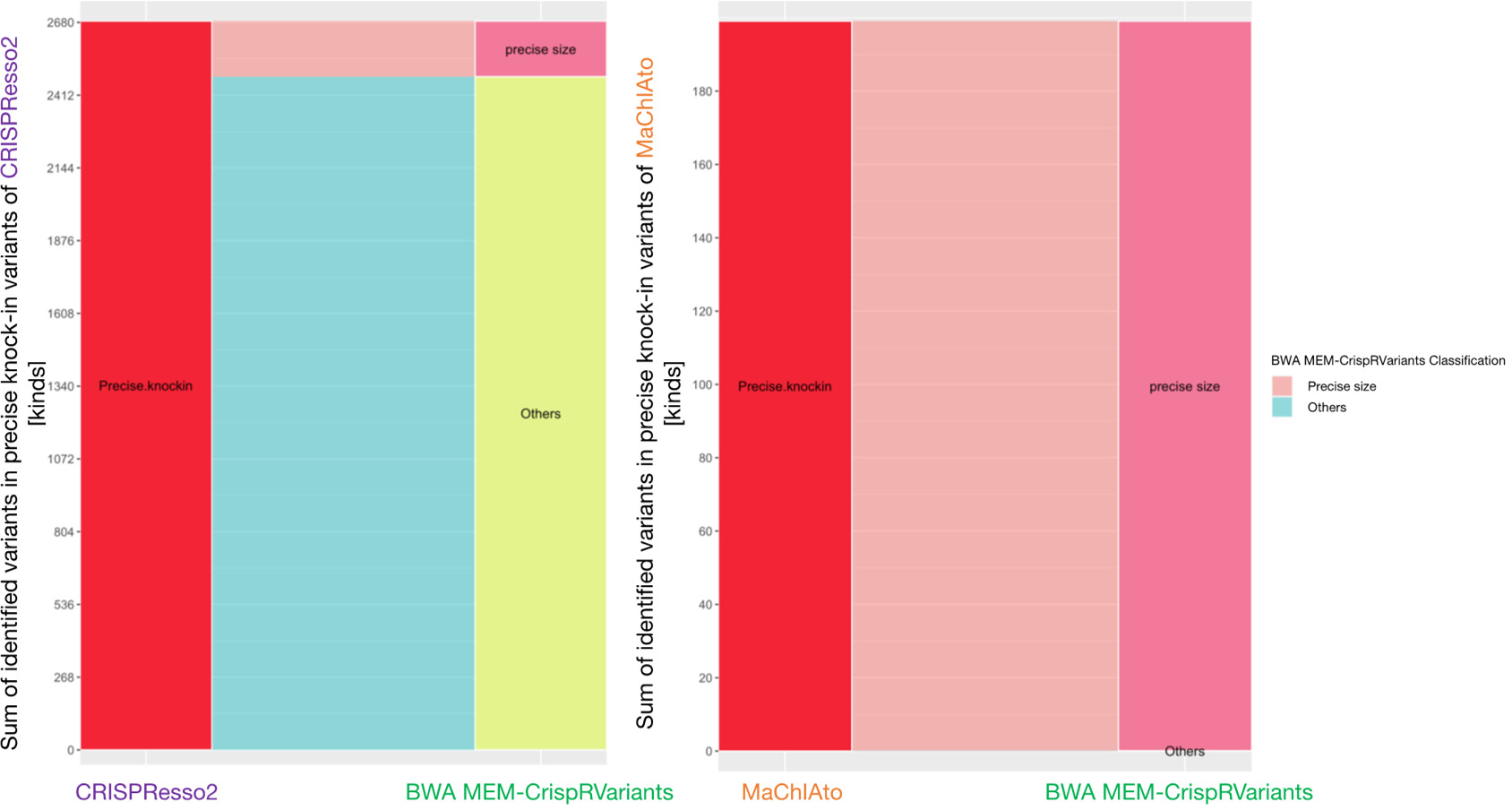
The comparison between BWA MEM-CrispRVariants and CRISPResso2 (left) or MaChIAto (right) using precise knock-in variants. Left parts of the left and right bar plots indicate the precise knock-in variants (HDR class) of CRISPResso2 and MaChIAto, respectively, and the y-axis is the number of identified variants in the precise knock-in variants. The right parts of these plots show the results of variant identification of BWA MEM-CrispRVariants using the corresponding variants. The variants are classified into a “precise size” class and an “Others” class in the variant identification of BWA MEM-CrispRVariants. The variants of the “precise size” class have an insert that is the expected length of the precise knock-in donor (66 bp) as decided by BWA MEM-CrispRVariants. The flows indicate the change in each class. The variants with the red color flow are classified into the “precise size” class, and the variants with blue color flow are classified into the “Others” class. The left plot is saved as “[j]Alluvial_Diagrams_of_CRISPResso_Knock-in_Class.png” in the output of MaChIAto Reviewer. The right plot is saved as “[k]Alluvial_Diagrams_of_MaChIAto_Knock-in_Class.png” in the output of MaChIAto Reviewer.

#### Supplementary Note 6 - Practical example for MaChIAto: a series of knock-in targeting 40 gene loci in HEK293T cells

We analyzed a series of knock-ins targeting 40 gene loci (Supplementary Note 4) using the CRISPResso2-MaChIAto pipeline to gain a deeper understanding of its applicability and features. We first estimated the precise knock-in efficiency of the samples (Supplementary Fig. 8). The precise knock-in reads were detected at all gene loci. The average efficiencies of precise knock-in (±SEM) were 7.03 (±0.77)% and 19.2 (±1.7)% for PITCh and LoAD, respectively, and the maximum efficiency, excluding unclassified classes, was 46.9%, observed with LoAD, where a 10.8-fold enhancement was achieved compared with PITCh (Supplementary Fig. 9). The result shows PITCh and LoAD knock-in can occur at various loci. However, the efficiency was dependent on the targeted locus.

**Supplementary Figure 8.**
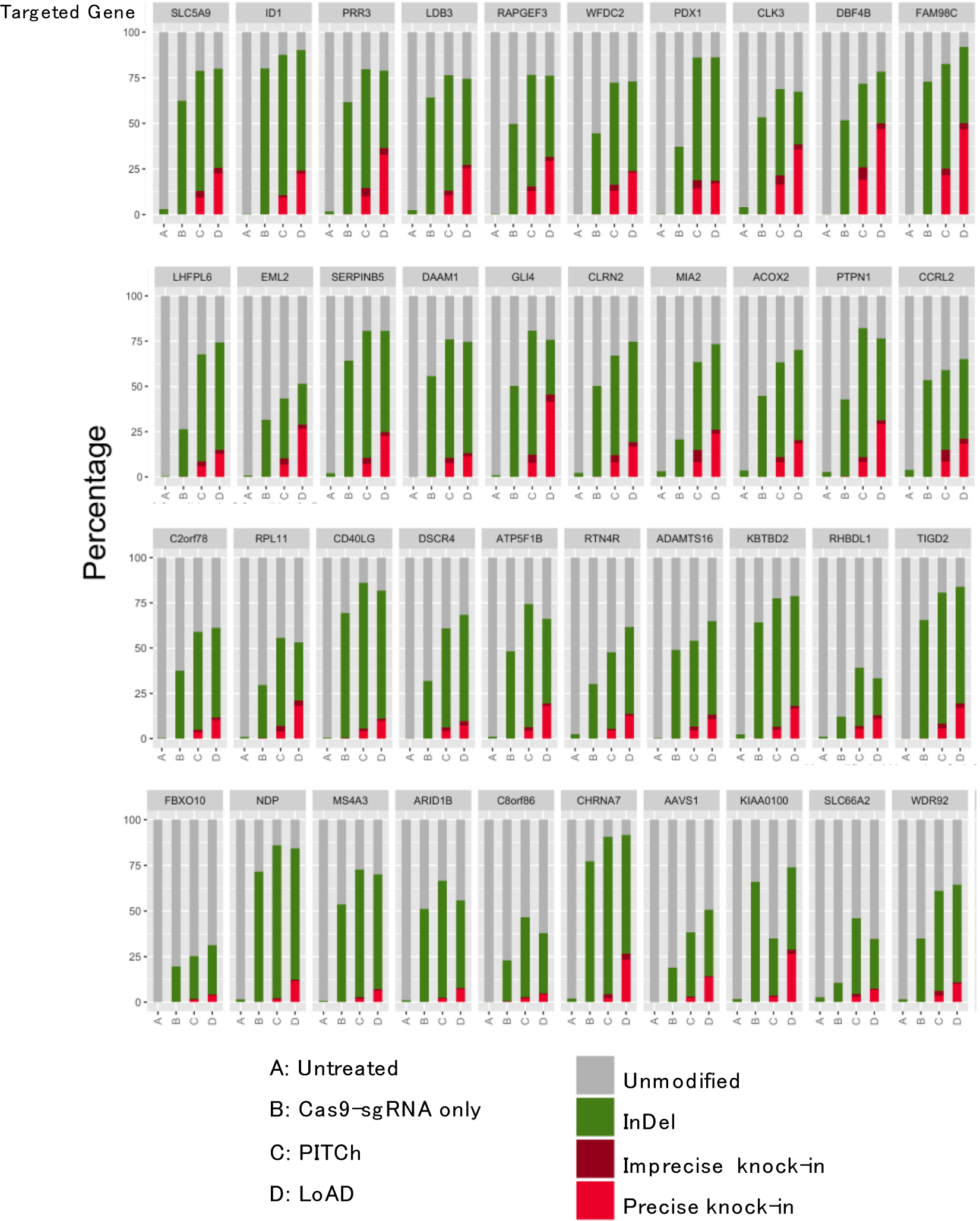
Overview of all the classification results. Each result of a targeted gene includes four sample types; The bottom labels of “A,” “B,” “C,” and “D” represent Untreated, Cas9-sgRNA only, PITCh, and LoAD samples, respectively. The bar plots are color-coded according to class; gray, green, dark red, and red boxes indicate Unmodified, NHEJ (represents InDel), Mixed HDR-NHEJ (represents imprecise knock-in), and HDR (represents precise knock-in) classes, respectively. The targeted genes are arranged in ascending order of the rate of precise knock-in in the LoAD sample. The plots are saved as “[b2]Summary_of_CRISPResso_MaChIAto_classification.png” in the output of MaChIAto Reviewer.

To understand the molecular mechanism of this variable efficiency, we investigated the relationship of various (epi-)genomic contexts of the targeted region using MaChIAto Analyzer which performed correlation analysis between knock-in efficacy and 130 kinds (epi-)genomic features as described in the Methods section. Surprisingly, sgRNA activity showed a weak correlation (|R| < 0.4) with precise knock-in efficiency or accuracy for both PITCh and LoAD knock-in (Supplementary Fig. 10a). Regarding the accuracy of PITCh knock-in, DNA accessibility on the PAM proximal (right) region, structural features such as propeller twist and duplex disrupt energy on both µHs showed relatively stronger positive correlation (R > 0.4, *p* < 0.05, Pearson correlation test) (Supplementary Fig. 10b). However, minimal energy for folding^13, 38^ and A-philicity^13, 39, 40^ on both µHs showed relatively stronger negative correlation (R < -0.4, *p* < 0.05, Pearson correlation test) (Supplementary Fig. 10c). We found that G bases in both µHs had a positive effect on accuracy (Supplementary Fig. 10b), whereas A bases in both µHs reduced accuracy (Supplementary Fig. 10c). However, interestingly, enhanced accuracy had a positive correlation with A base frequency (Supplementary Fig. 10d), which means that LoAD can recover the negative effect of A bases with respect to accuracy. The recovery effect on accuracy was also observed for minimum energy folding. We summarized the preferred features and undesirable features (|R| > 0.4, *p* < 0.05, Pearson correlation test) involved in knock-in accuracy in a table (Supplementary Fig. 11).

**Supplementary Figure 9.**
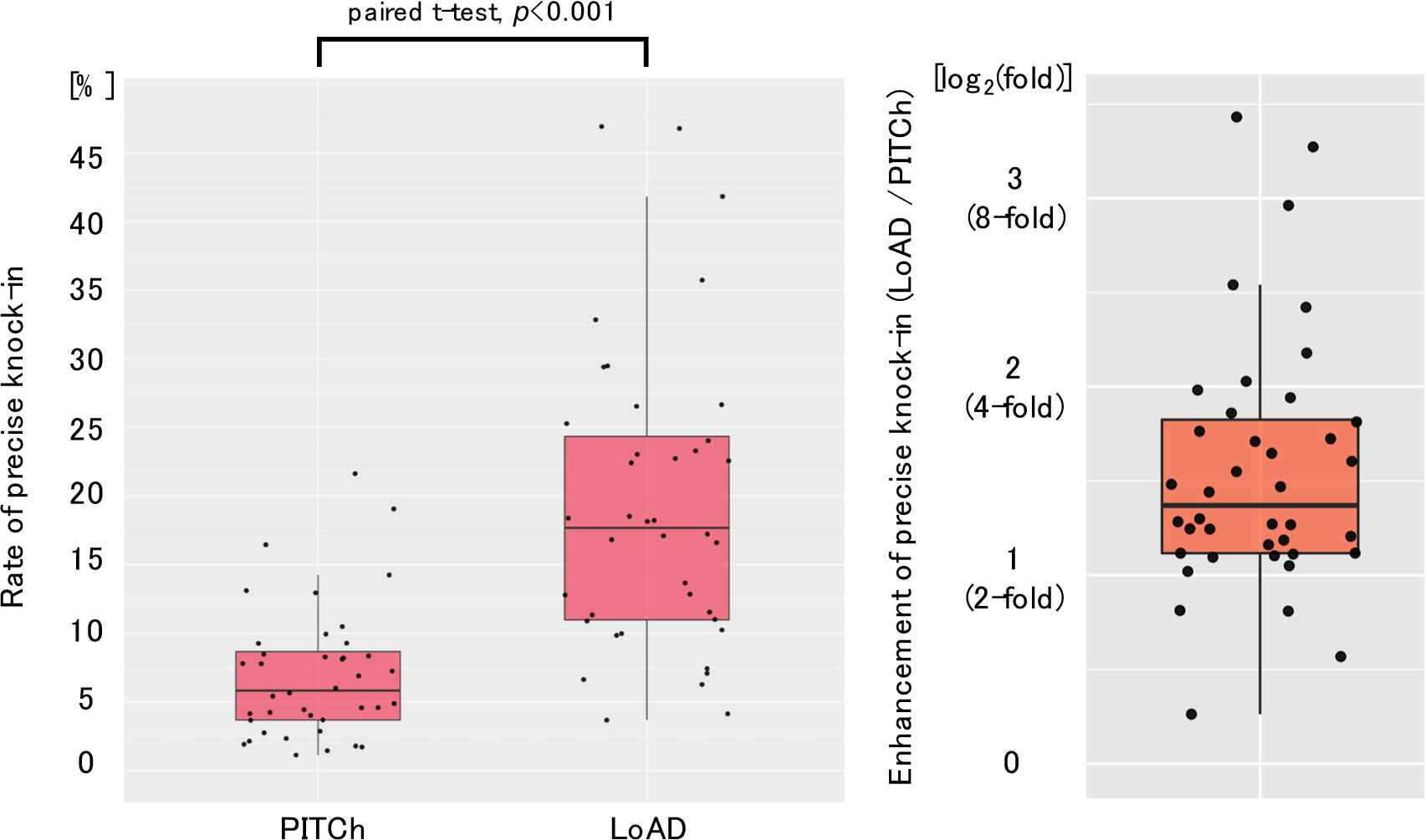
Summary of precise knock-in in the knock-in series targeting 40 gene loci. The left box plot represents the rates of precise knock-in excluding the Unclassified class. Each dot represents a result. The box plot is split by the sample type (PITCh and LoAD). The right box plot shows the enhancement of precise knock-in. The values are calculated as log2-fold change. The ratio (-fold) is (Rate of precise knock-in excluding the Unclassified class in LoAD) / (Rate of precise knock-in excluding the Unclassified class in PITCh). The left plot is saved as “[l7]Barplot_of_Rate_of_Precise_knock-in.png” in the output of MaChIAto Reviewer. The right plot is saved as “[l11]Barplot_of_Enhancement_Reduction.png” in the output of MaChIAto Reviewer.

**Supplementary Figure 10.**
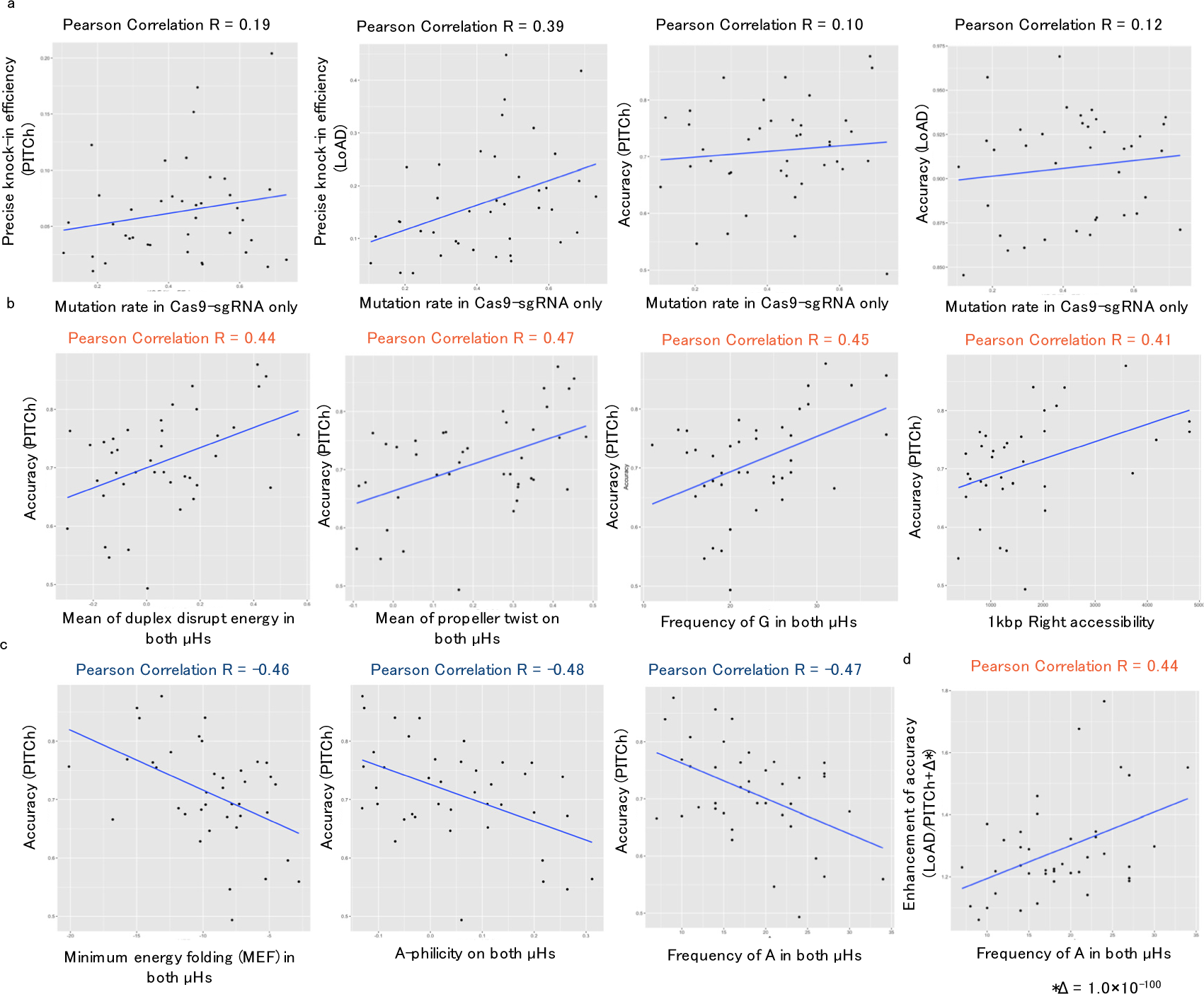
Results of correlation analysis between knock-in efficiency (precise knock-in efficiency, accuracy, and enhancement) and (epi-)genomic contexts. The y-axis of the scatter plot is the knock-in efficiency, and the x-axis of that is the value of the feature. The line in the scatter plot is a simple linear regression using the least squares method. The label above the plot represents the Pearson correlation. If the correlation (R) is R > 0.4 or R < 0.4, the label color is orange or blue, respectively. a. Scatter plots between a mutation rate in a Cas9-sgRNA only sample and knock-in efficiency of PITCh and LoAD. The mutation rate is “Editing efficiency” in the output of MaChIAto Classifier. b. Scatter plots showing a stronger positive correlation with accuracy in PITCh samples. c. Scatter plots showing a stronger negative correlation with accuracy in PITCh samples. d. Scatter plot between the frequency of A in both µHs and the enhancement of accuracy [loge(Accuracy in LoAD / Accuracy in PITCh)]. All correlations are statistically significant (*p* < 0.05, correlation test). The plots are saved in the directory of “knock-in_1_Accuracy,” “knock-in_2_Accuracy,” “knock- in_1_Precise.Knock.in.Efficiency,” and “knock-in_2_Precise.Knock.in.Efficiency” in the output of MaChIAto Analyzer.

Some features in the table have been known as the factors related to MMEJ repair. For example, Tm (melting temperature) value, which is one of the significant features regarding knock-in accuracy, is involved in the MMEJ activity in the yeast study^41^. The bendability of nucleotides has never been known as a knock-in factor, whereas the factor has been featured as one of the factors for sgRNA efficiency in the data of a previous machine learning study^13^. However, as we mentioned before, sgRNA activity (mutation rate in the samples of Cas9-sgRNA only) was not correlated with knock-in accuracy (Supplementary Fig. 10a). There may be an unknown mechanism, which is independent of sgRNA activity, inducing MMEJ knock-in without InDels. The folding energy^42, 43^ is also the feature affecting sgRNA activity, but the value of this study was dependent on both microhomologies. Therefore, the structural stability of homology arms might contribute to knock-in accuracy. The propeller twist is one of the structural factors affecting protein-DNA binding^42^. Perhaps, the binding activity of MMEJ-associated proteins have preference for DNA structure in microhomologous regions.

The accuracy of PITCh knock-in depended on various (epi-)genomic features, whereas that of LoAD was dependent on fewer features. The number of significant features decreased at least eight-fold (Supplementary Fig. 12). Some box plots, which collected the values of correlation by feature group, showed that dependency regarding accuracy can be decreased by LoAD (Supplementary Fig. 13, left). For example, regarding the accuracy, the median of DNA accessibility got close to no correlation under LoAD (Supplementary Fig. 13, left). Moreover, regarding the precise knock-in efficiency, LoAD changed the dependencies of many features. As a result, the distribution of all correlations significantly converged to the zero correlation (Supplementary Fig. 13, right, All features).

**Supplementary Figure 11.**
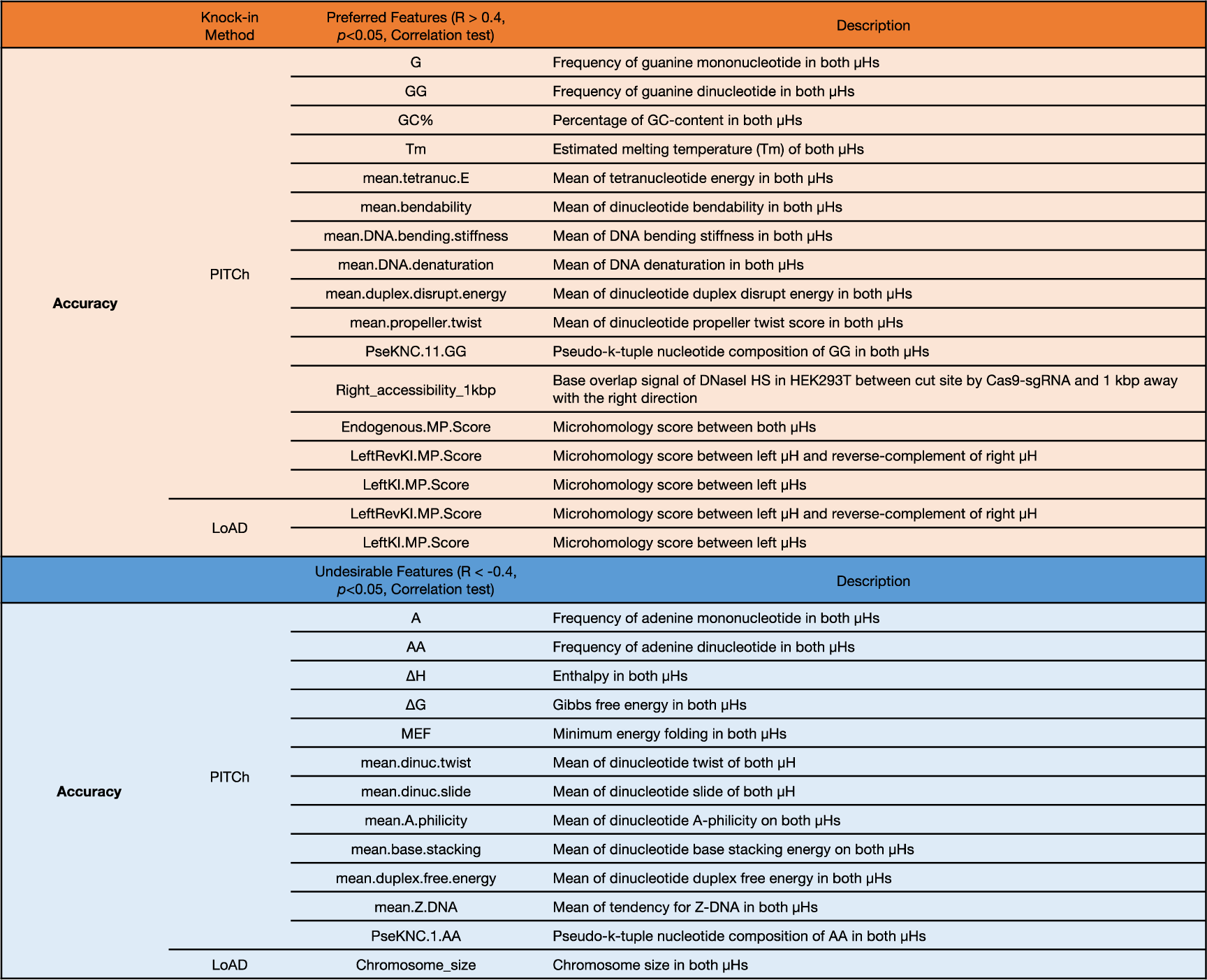
List of preferred and undesirable features involved in knock- in accuracy.

**Supplementary Figure 12.**
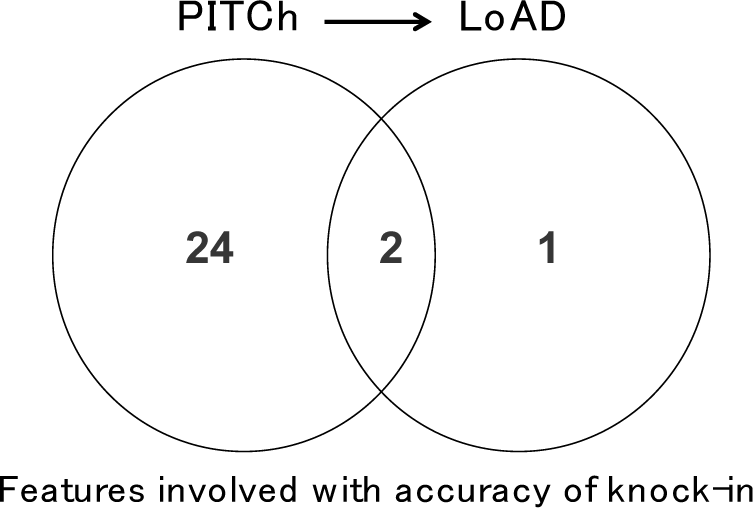
Comparison of the number of preferred features and undesirable features involved in knock-in accuracy of two knock-in methods, PITCh and LoAD. LoAD decreased the number of features.

By the way, the InDel class that had endogenous InDels without knock-in sequence seemed to have a more important role in knock-in efficiency because the scale of InDel reads was generally larger than that of imprecise knock-ins (Supplementary Fig. 14). Moreover, the rank of the rate of imprecise knock-in was relatively the same as that of the rate of precise knock-in, whereas the rank of the rate of InDel was more different from that of the rate of precise knock-in (Supplementary Fig. 15). The change in rank between precise knock-in and InDel was more dynamic, indicating that the frequency and pattern of InDels controlled the knock-in outcome. To investigate this effect, we analyzed the mutation pattern of InDel and imprecise knock-in using MaChIAto Aligner and MaChIAto Reviewer. MaChIAto Aligner used the unmodified amplicon sequence as the reference of alignment for the InDel reads. MaChIAto Aligner also used the expected sequence of the precise and imprecise knock-ins as the reference of alignment for the imprecise knock-in reads. The mutation information was extracted from the alignment data and they were analyzed using the grouping analysis of MaChIAto Reviewer. The group with precise knock-in efficiency in PITCh is shown in Supplementary Fig. 15. We observed that there were relatively fewer inserts in the PAM-proximal position in the group of low precise knock-in efficiency in PITCh (Supplementary Fig. 16). Regarding InDel size of reads in the InDel class, more reads in the low group have a +1-bp InDels compared with those in the high group (Supplementary Fig. 17, top). In the high group, there was no gene with a +1-bp InDels as the most frequent mutant (Supplementary Fig. 17, bottom, I1).

**Supplementary Figure 13.**
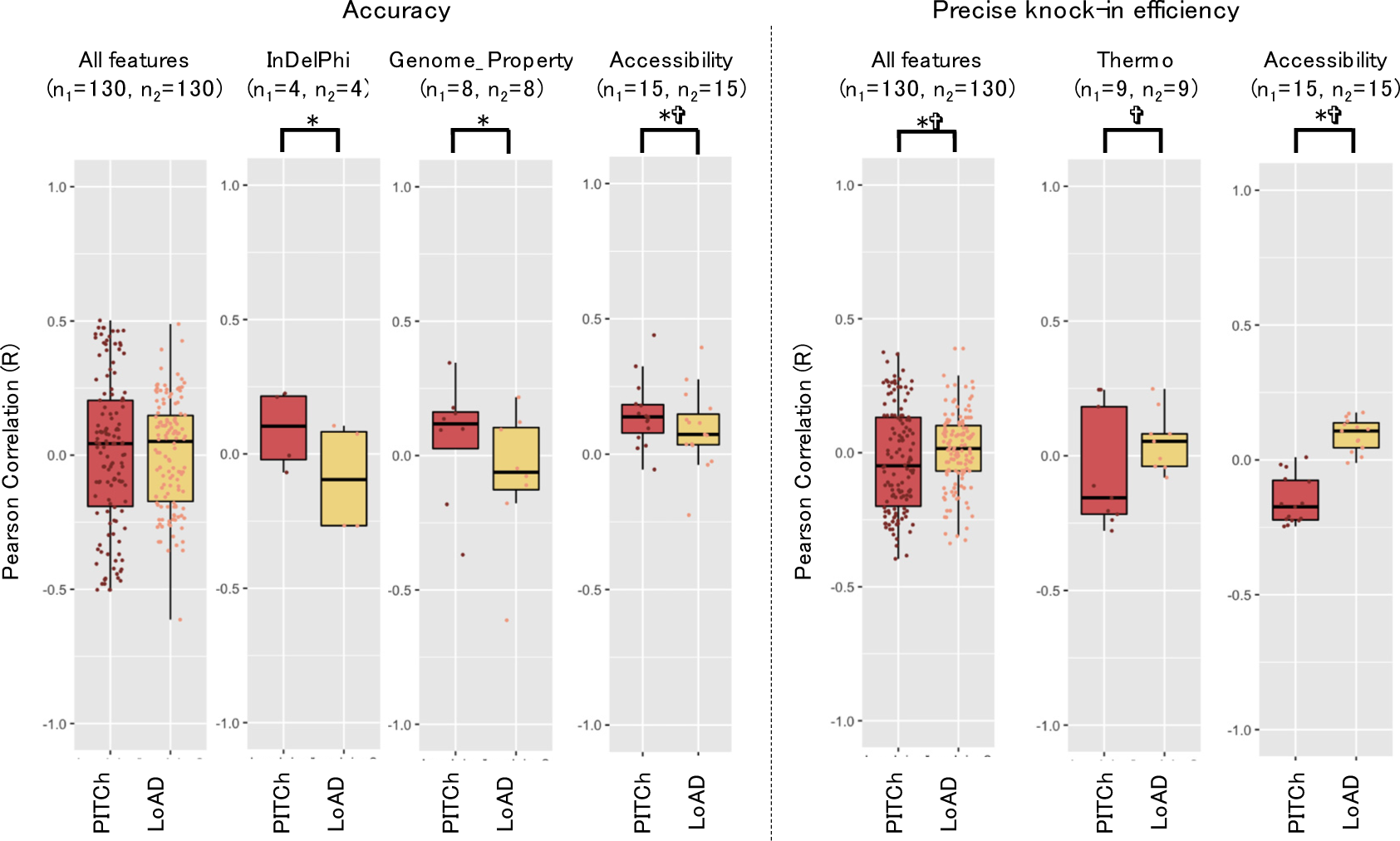
Boxplots of Pearson correlation values by feature group. The left plots are for accuracy, and the right ones are for precise knock-in efficiency. The left box plot of each plot area is the result of PITCh, and the right box plot of that is LoAD. The y-axis is the Pearson correlation with the knock-in efficiency, which is shown in the above plots. The asterisk and cross indicate the significant difference between PITCh and LoAD; *p* < 0.05, paired *t*-test and *p* < 0.05, Kolmogorov– Smirnov test, respectively. The n1 is the number of features used for calculating the correlation with knock-in efficiency of PITCh, and the n2 is that of LoAD. The n1 is different from the n2 in the plot of all features regarding accuracy because there is an incalculable condition in a part of the process. The plots are saved in the directory of “comparison_Accuracy” and “comparison_Precise.Knock.in.Efficiency” in the output of MaChIAto Analyzer.

**Supplementary Figure 14.**
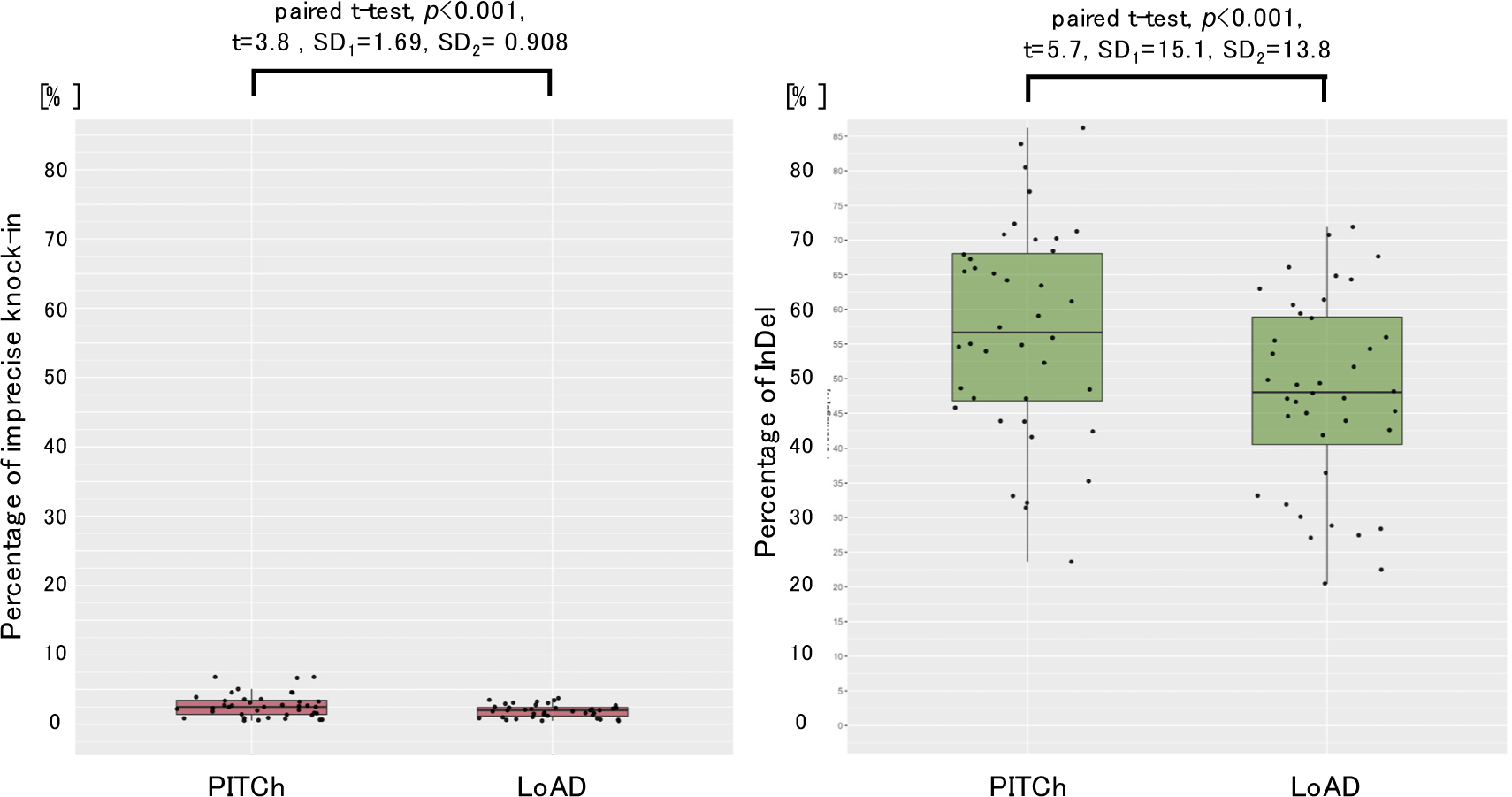
Summary of imprecise knock-in and InDel in the knock-in series targeting 40 gene loci. The left box plot represents the rates of imprecise knock- in excluding the Unclassified class, and the right box plot represents the rates of InDels excluding the Unclassified class. Each dot represents one result. The box plot is split by the sample type (PITCh and LoAD). t and SD represent t-statistic and standard deviation, respectively. The left plot is saved as “[l8]Barplot_of_Rate_of_Imprecise_knock-in.png” in the output of MaChIAto Reviewer. The right plot is saved as “[l9]Barplot_of_Rate_of_InDel.png” in the output of MaChIAto Reviewer.

In addition, we distinguished MMEJ-assisted deletion with NHEJ-assisted deletion using MaChIAto Aligner and MaChIAto Reviewer (Supplementary Fig. 18, top). -2-bp MMEJ-assisted InDels occurred more frequently in the low group, while - 2-bp NHEJ-assisted InDels were more frequently observed in the high group. Interestingly, multiples of -3-bp MMEJ-assisted InDels, which resulted in frame-in mutations, often occurred in the high group. To further check the frequency, the periodograms that MaChIAto Reviewer provides for MMEJ-assisted deletion and NHEJ-assisted deletion per group were assessed (bottom, Supplementary Fig. 18). As a result, the obvious frequency peak was observed at the period of 3 bp (frequency = 0.33) only in the high group of the MMEJ-assisted knock-in. Next, we focused on the sequence involved in the endogenous MMEJ deletions, not MMEJ knock-in (Supplementary Fig. 19). The high group had many G and CG bases in “deletion-related µH (DµH)” which caused endogenous MMEJ deletions (Supplementary Fig. 19, left). The MMEJ deletion with a -1-bp DµH showed a stronger tendency with the frequency of G bases (Supplementary しゅ Fig. 19, center). Regarding the MMEJ deletion with certain CG DµHs often existed in the high group (Supplementary Fig. 19, right). The guanine and cytosine near the target site reportedly promote the pathway activity of MMEJ^19, 20^ between the two cut ends in the genomic locus. Thus, the MMEJ activity in the endogenous site might propagate the activity of the MMEJ repair between the chromosome and the donor. Additionally, we focused on the intervening sequence between two DµHs that were supposedly not related to wthe DSB repair process because it was removed from the genomic sequence along with one unit of µH through the DSB repair (top, Supplementary Fig. 20). The low group showed a lot of MMEJ- assisted InDels without the intervening sequence (bottom, Supplementary Fig. 20), and the group had fewer InDels with G and CG intervening sequence (Supplementary Fig. 21). Other features also showed outstanding values in the low or high group in the plot of the cumulative ratio; however, the most of them did not show a significant difference between the high and low precise PITCh knock-in efficiency groups in *t*-test (*p* > 0.05), which did not seem to have an obvious tendency.

**Supplementary Figure 15.**
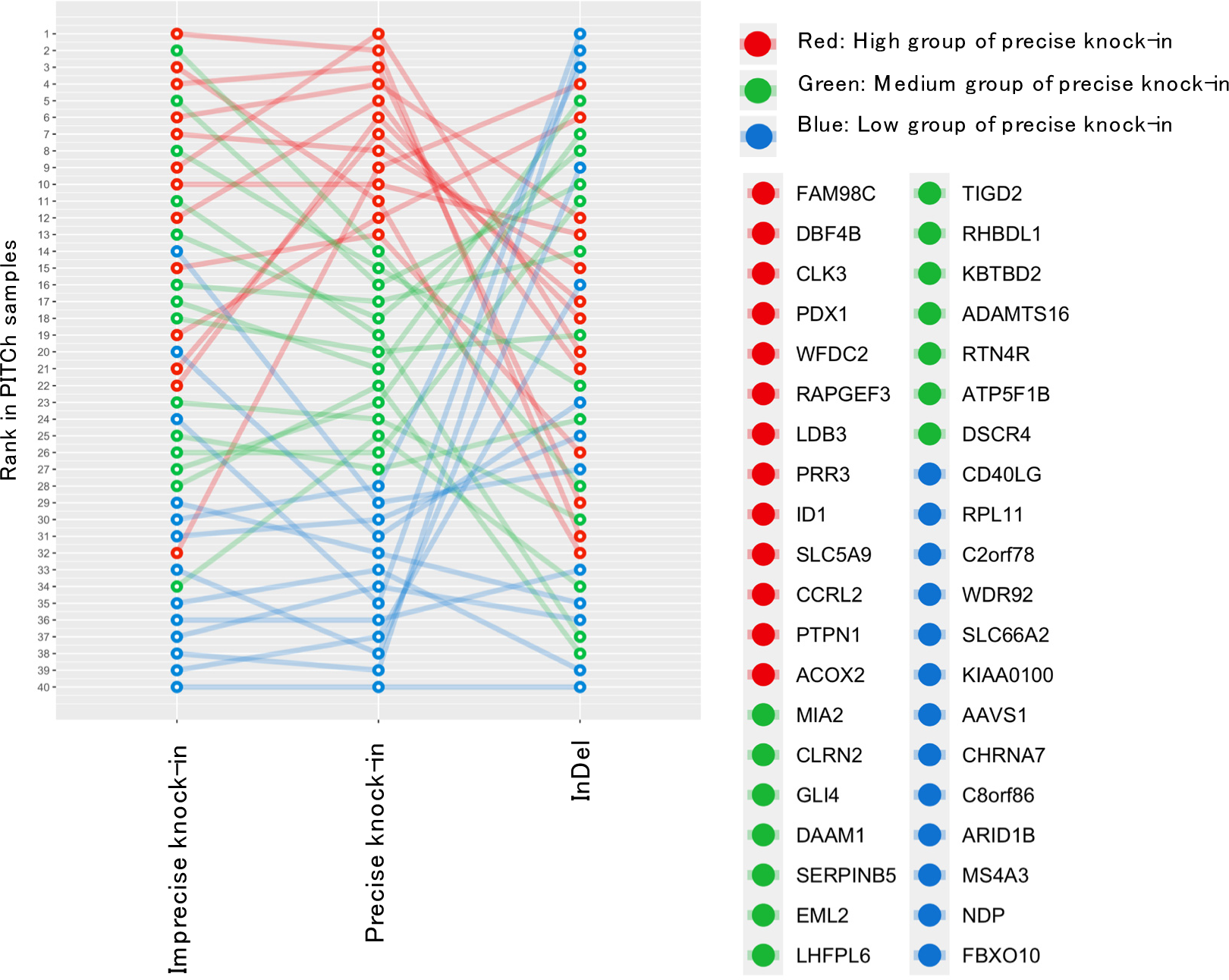
Rank relationship among the rates of imprecise knock-in, precise knock-in, and InDel in PITCh. The rank is arranged in ascending order per class. The lower rank numbers represent the higher value of the rate. Gene groups showing high, medium, and low rates of precise knock-in are indicated in red, green, and blue circles, respectively. The points joined with the colored line represent the same gene. The plot is saved as “[l4]Relation_of_Rank_of_Class_on_MaChIAto_in_condition1.png” in the output of MaChIAto Reviewer.

**Supplementary Figure 16.**
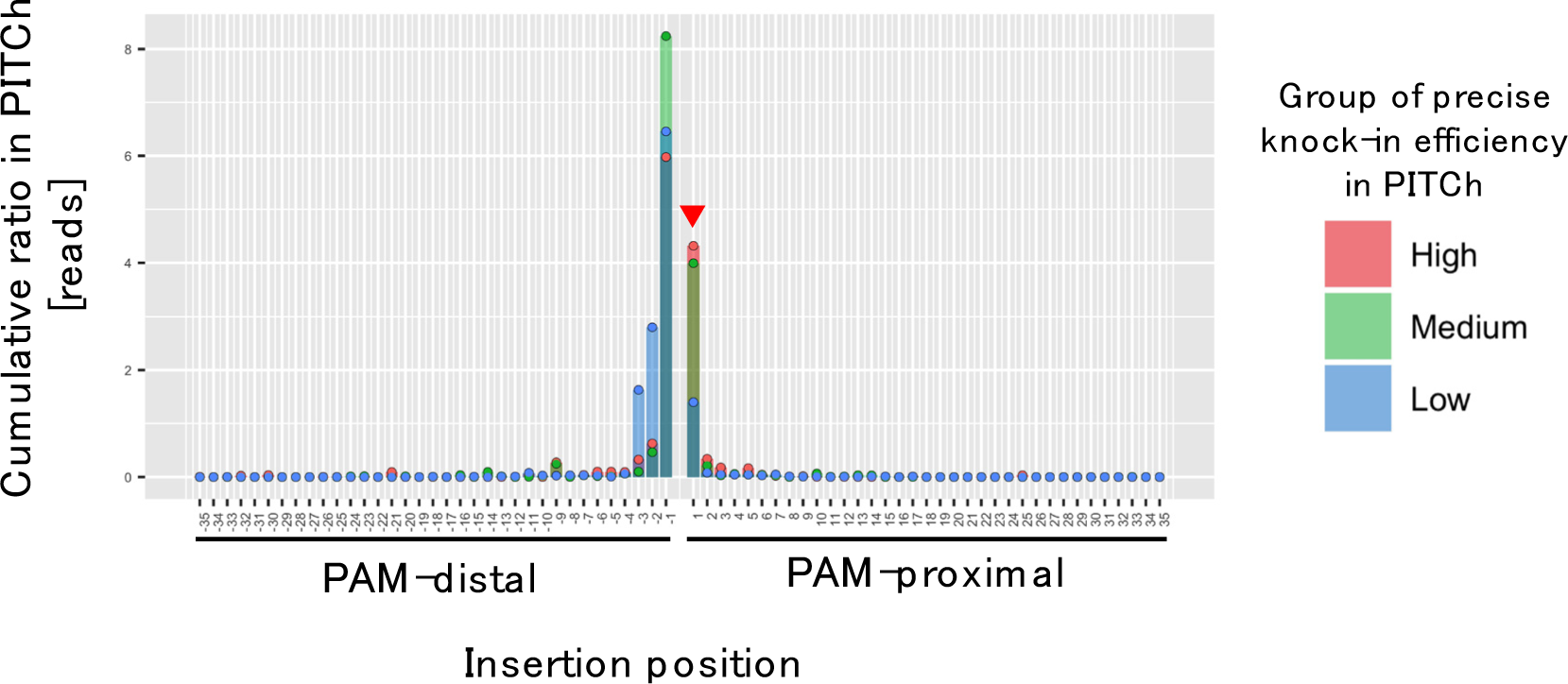
Distribution of insert position in the InDel class. The value is the cumulative ratio of variant labels which MaChIAto Aligner provides. The cumulative ratio is calculated per group. The groups overlap on the plot, and the red, green, and blue points indicate the cumulative ratio of the high, medium, and low groups, respectively, regarding the rate of precise knock-in in PITCh. The red arrowhead represents the preferred features of precise knock-in in PITCh. The plot is saved as “[a1- 1]Barplot_of_insert_position_of_condition1_reads_by_condition1_group.png” in the output of MaChIAto Reviewer.

**Supplementary Figure 17.**
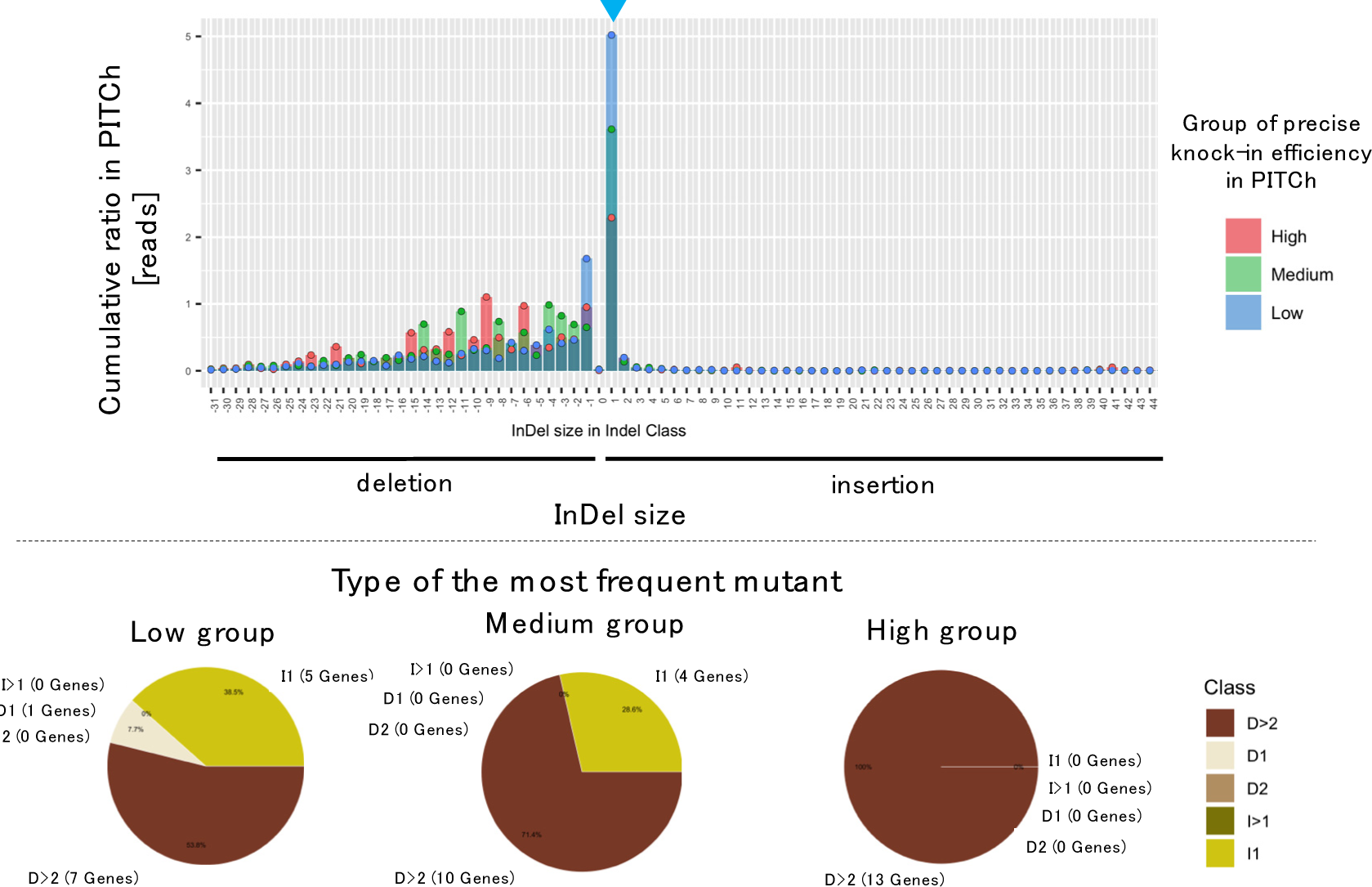
Distribution of InDel size (top) and type of the most frequent mutant per group (bottom) in the InDel class. Top. The value is the cumulative ratio of variant reads identified with MaChIAto Aligner. The cumulative ratio is calculated per group. The groups overlap on the plot and the red, green, and blue points indicate the cumulative ratio of high, medium, and low groups, respectively, regarding the rate of precise knock-in in PITCh. The blue arrowhead represents the undesirable feature of precise knock-in in PITCh. Bottom. The most frequent mutant per group is expressed as a pie chart. The labels D1, D2, D>2, I1 and I>1 represent mutations of 1-bp deletion, 2-bp deletion, > 2-bp deletion, 1-bp insertion, and > 1-bp insertion, respectively. The top plot is saved as “[b- 1]Barplot_of_mutation_indel_size_of_condition1_reads_by_condition1_group.png” in the output of MaChIAto Reviewer. The bottom plots are saved as “[d- 1]Piechart_of_max_indel_size_class_in_condition1_reads_among_condition1_low_g roup.png,” “[d-1]Piechart_of_max_indel_size_class_in_condition1_reads_among_condition1_mediu m_group.png,” and “[d- 1]Piechart_of_max_indel_size_class_in_condition1_reads_among_condition1_high_ group.png” in the output of MaChIAto Reviewer.

**Supplementary Figure 18.**
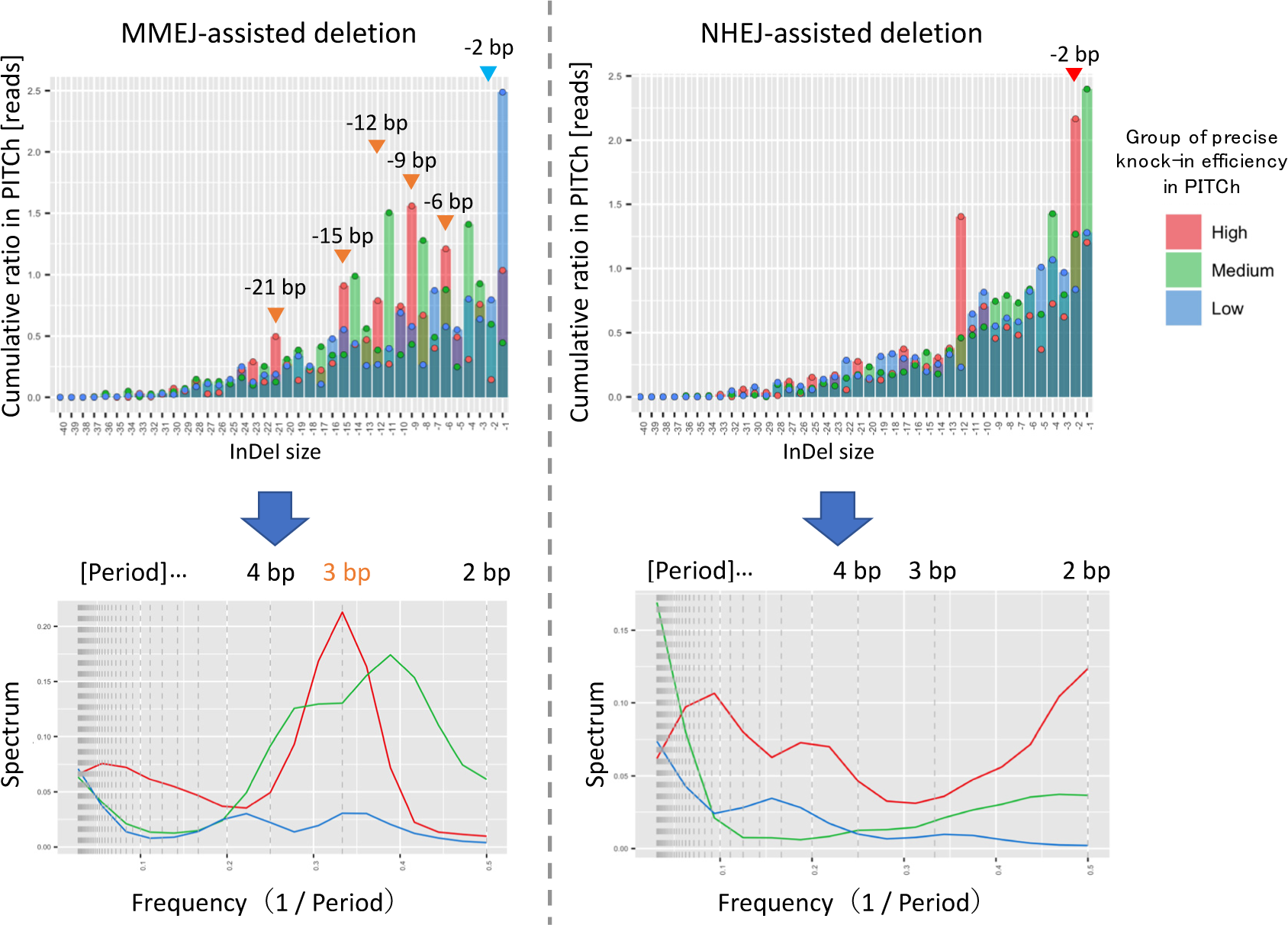
Distributions of deletion size by MMEJ-assisted mutation (top left) and NHEJ-assisted mutation (top right), and periodograms of MMEJ-assisted deletion (bottom left) and NHEJ-assisted deletion (bottom right) in the InDel class. Top left and top right. The value is the cumulative ratio of variant reads identified with MaChIAto Aligner. The cumulative ratio is calculated per group. The groups overlap on the plot, and the red, green, and blue points indicate the cumulative ratio of the high, medium, and low groups, respectively, regarding the rate of precise knock-in in PITCh. The red and blue arrowheads represent the preferred and undesirable features of precise knock-in in PITCh, respectively. Bottom left and bottom right. The y-axis is the estimated spectrum, and the x-value is the frequency. The period [bp] can be obtained by taking the reciprocal of the frequency. The red, green, and blue lines represent high, medium, and low group values, respectively, regarding the rate of precise knock-in in PITCh. The top plots are saved as “[j-1]Barplot_of_mutation_MMEJ- mediated_indel_size_of_condition1_reads_by_condition1_group.png” and “[k- 1]Barplot_of_mutation_NHEJ- mediated_indel_size_of_condition1_reads_by_condition1_group.png” in the output of MaChIAto Reviewer. The bottom plots are saved as “[j- 1]Barplot_of_mutation_MMEJ- mediated_indel_size_of_condition1_reads_by_condition1_group_periodogram.png” and “[k-1]Barplot_of_mutation_NHEJ-mediated_indel_size_of_condition1_reads_by_condition1_group_periodogram.png” in the output of MaChIAto Reviewer.

**Supplementary Figure 19.**
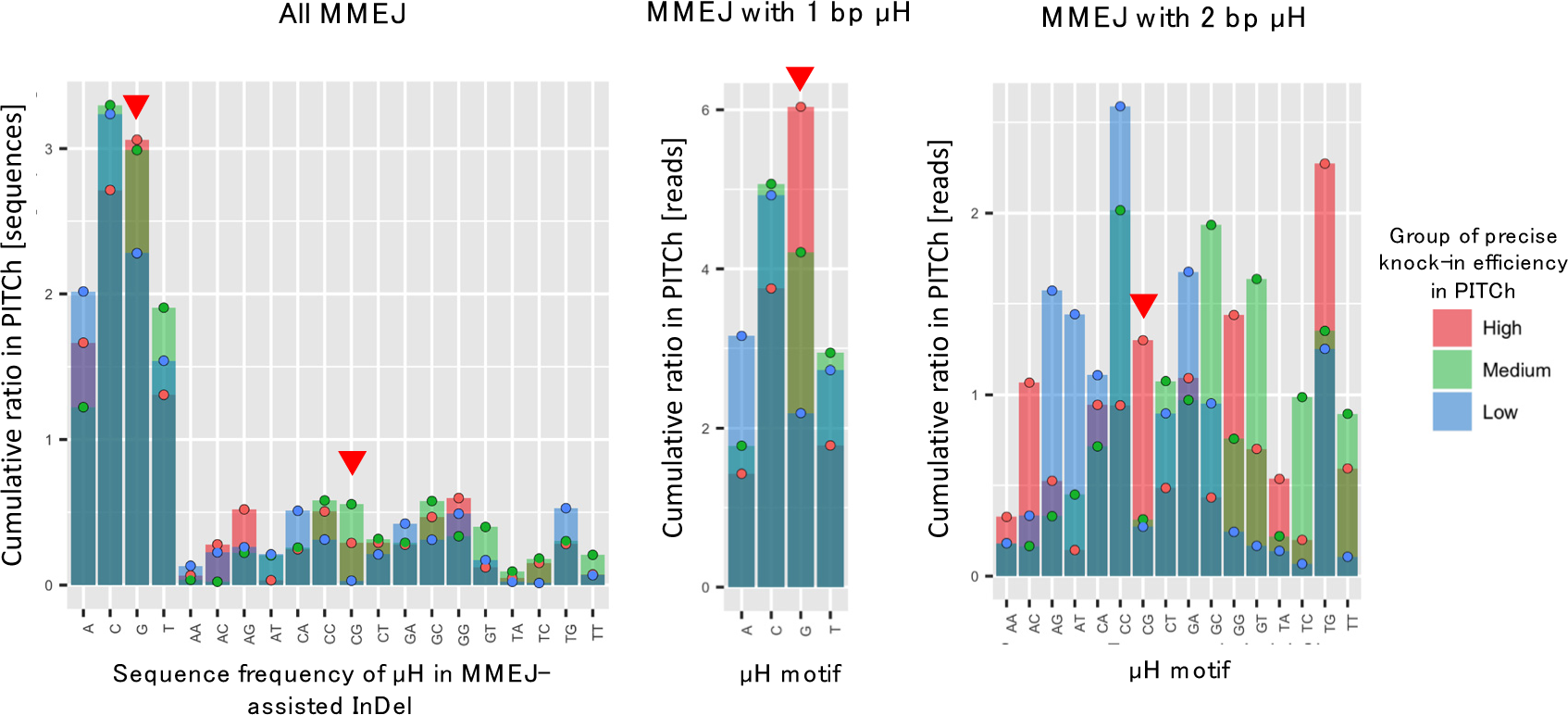
Distributions of mono- and di-nucleotide frequency in microhomologies involved in MMEJ-assisted mutation (left) and distributions of microhomology sequence involved in MMEJ-assisted mutation with 1-bp microhomology (center) and 2-bp microhomology (right) in the InDel class. The cumulative ratio is calculated per group. The groups overlap on the plot, and the red, green, and blue points indicate the cumulative ratio of the high, medium, and low groups, respectively, regarding the rate of precise knock-in in PITCh. The red and blue arrowheads represent the preferred and undesirable features of precise knock-in in PITCh, respectively. Left. The value is the cumulative ratio of sequence frequency identified with MaChIAto Aligner. Center and right. The value is the cumulative ratio of variant reads identified with MaChIAto Aligner. The left plot is saved as “[p- 1]Barplot_of_microhomology_sequence_frequency_of_condition1_reads_by_conditi on1_group.png” in the output of MaChIAto Reviewer. The center plot is saved as “[q- 1]Barplot_of_mono- microhomology_sequence_frequency_of_condition1_reads_by_condition1_group.pn g” in the output of MaChIAto Reviewer. The right plot is saved as “[r-1]Barplot_of_di- microhomology_sequence_frequency_of_condition1_reads_by_condition1_group.pn g” in the output of MaChIAto Reviewer.

**Supplementary Figure 20.**
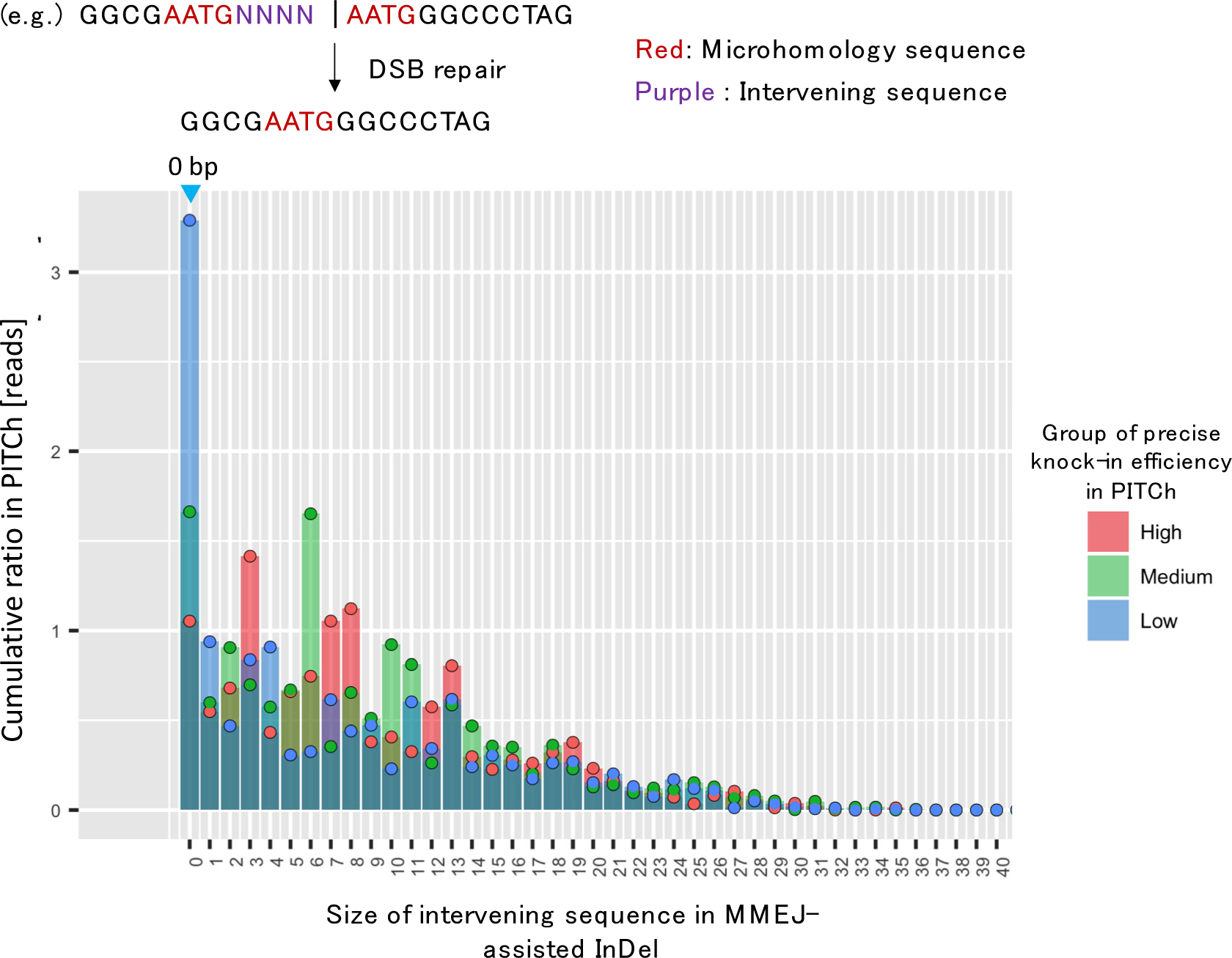
Distributions of the size of intervening sequence involved in MMEJ-assisted mutation. The cumulative ratio is calculated per group. The groups overlap on the plot, and the red, green, and blue points indicate the cumulative ratio of the high, medium, and low groups, respectively, regarding the rate of precise knock-in in PITCh. The blue arrowhead represents the undesirable features of precise knock-in in PITCh. The value is the cumulative ratio of variant reads identified with MaChIAto Aligner. The plot is saved as “[m-1]Barplot_of_mutation_MMEJ- mediated_intervening_size_of_condition1_reads_by_condition1_group.png” in the output of MaChIAto Reviewer.

**Supplementary Figure 21.**
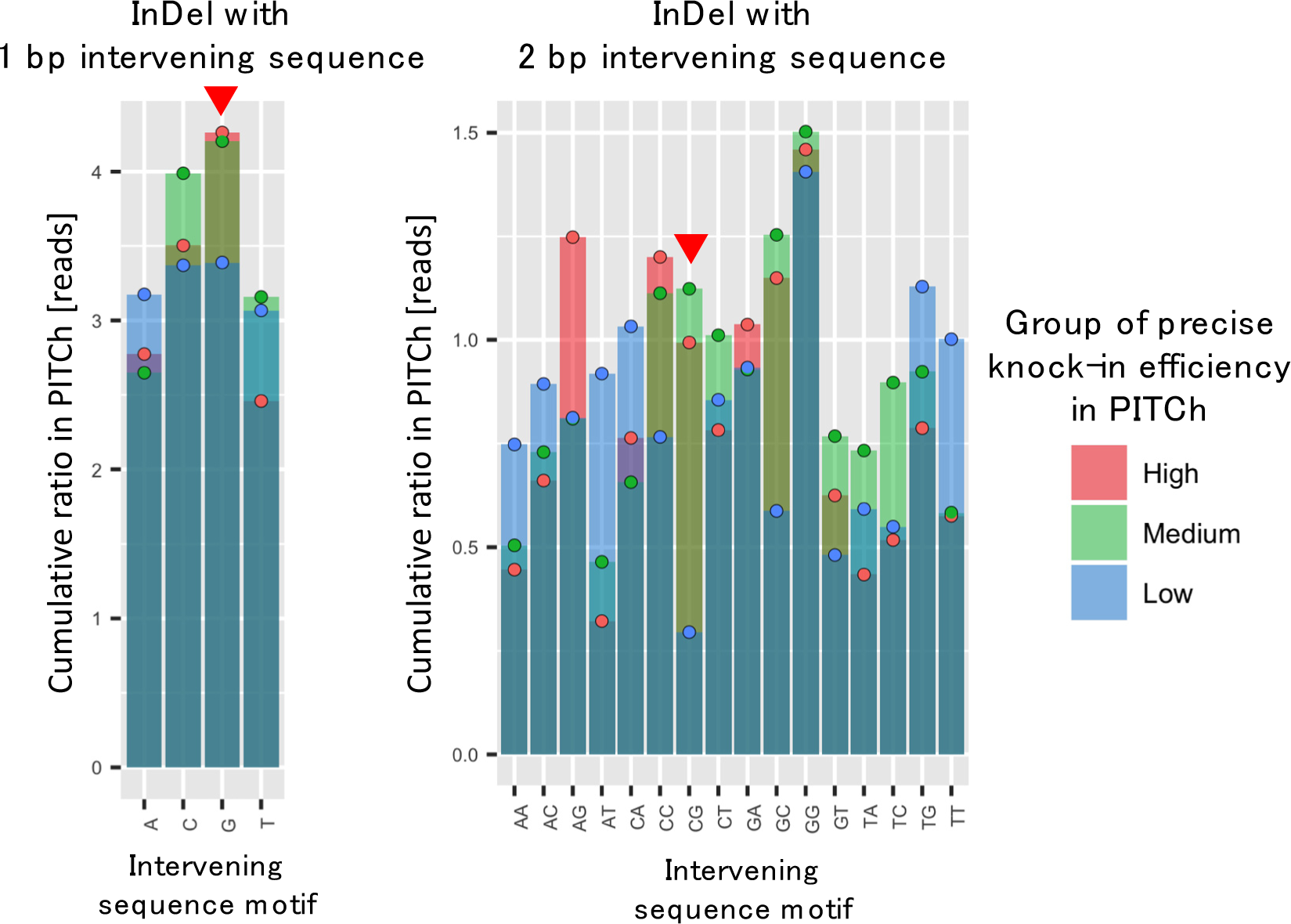
Distributions of mono- and di-nucleotide intervening sequence involved in mutation with 1-bp intervening sequence (left) and 2-bp intervening sequence (right) in the InDel class. The cumulative ratio is calculated per group. The groups overlap on the plot, and the red, green, and blue points indicate the cumulative ratio of the high, medium, and low groups, respectively, regarding the rate of precise knock-in in PITCh. The red arrowheads represent the preferred features of precise knock-in in PITCh. The value is the cumulative ratio of variant reads identified with MaChIAto Aligner. The left plot is saved as “[v-1]Barplot_of_mono- intervening_sequence_frequency_of_condition1_reads_by_condition1_group.png” in the output of MaChIAto Reviewer. The right plot is saved as “[w-1]Barplot_of_di- intervening_sequence_frequency_of_condition1_reads_by_condition1_group.png” in the output of MaChIAto Reviewer.

We then looked at the pattern of mutations in knock-in junctions in reads that belonged to the imprecise knock-in class. The most frequent sizes of the left knock-in junction in the imprecise knock-in class were equal to the expected imprecise left junction generated by NHEJ (Supplementary Fig. 22), which meant that, in a case of NHEJ-assisted integration, most knock-in inserts joined the 5′ end of the cut site without any InDels. However, in the right junction of the low group, many NHEJ- assisted integrations with +1-bp InDels were observed (Supplementary Fig. 23). We have summarized the results of the significant mutation patterns as a table (Supplementary Fig. 24).

**Supplementary Figure 22.**
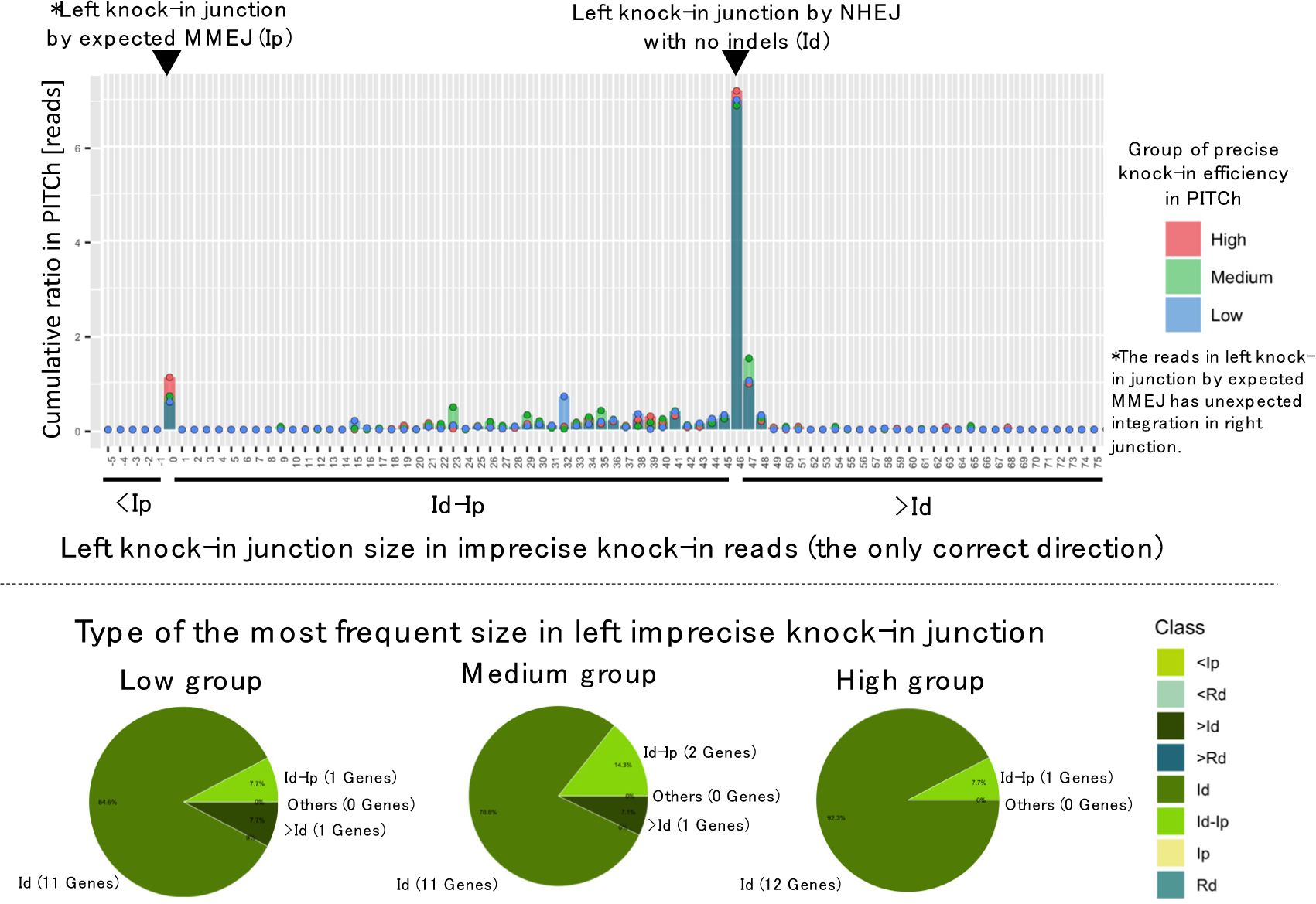
Distribution of left knock-in junction size (top) and the most frequent size in the left imprecise knock-in junction per group (bottom) in the imprecise knock-in class. Top. The value is the cumulative ratio of variant reads identified with MaChIAto Aligner. The cumulative ratio is calculated per group. The groups overlap on the plot, and the red, green, and blue points indicate the cumulative ratio of the high, medium, and low groups, respectively, regarding the rate of precise knock-in in PITCh. The labels “<Ip,” “Ip,” “Id-Ip,” “Id,” and “>Id” mean the size class in the junction. The “Ip” reads showed the same size as the precise knock-in reference. Some have precise junction sequences; however, they do not have that of the right junction. The values do not include the reads of the left junction of the reverse knock-in. Bottom. The most frequent left junction size per group is expressed as a pie chart. The classes include “Ip” (imprecise junction sequence with the same length as that of the precise junction reference), “Id” (imprecise junction sequence with the same length as that of the imprecise junction reference, which has duplicated left/right homology arm and left/right extra sequence), “<Ip” (imprecise junction sequence with a shorter length than that of the precise junction reference), “Id-Ip” (imprecise junction sequence with a longer length than that of the precise junction reference but shorter than that of the imprecise junction reference), “Rd” (reverse imprecise junction sequence with the same length as that of the reverse imprecise junction reference, which has combined left and right homology arms and right/left extra sequence), “<Rd” (reverse imprecise junction sequence with a shorter length than that of the reverse imprecise junction reference) and “>Rd” (reverse imprecise junction sequence with a longer length than that of the reverse imprecise junction reference). The top plot is saved as “[a- 1]Barplot_of_left_junction_of_indel_size_of_condition1_reads_by_condition1_group .png” in the output of MaChIAto Reviewer. The bottom plots are saved as “[c- 1]Piechart_of_max_indel_size_class_on_left_junction_in_condition1_reads_among_ condition1_low_group.png,” “[c-1]Piechart_of_max_indel_size_class_on_left_junction_in_condition1_reads_among_ condition1_medium_group.png,” and “[c- 1]Piechart_of_max_indel_size_class_on_left_junction_in_condition1_reads_among_ condition1_high_group.png” in the output of MaChIAto Reviewer.

**Supplementary Figure 23.**
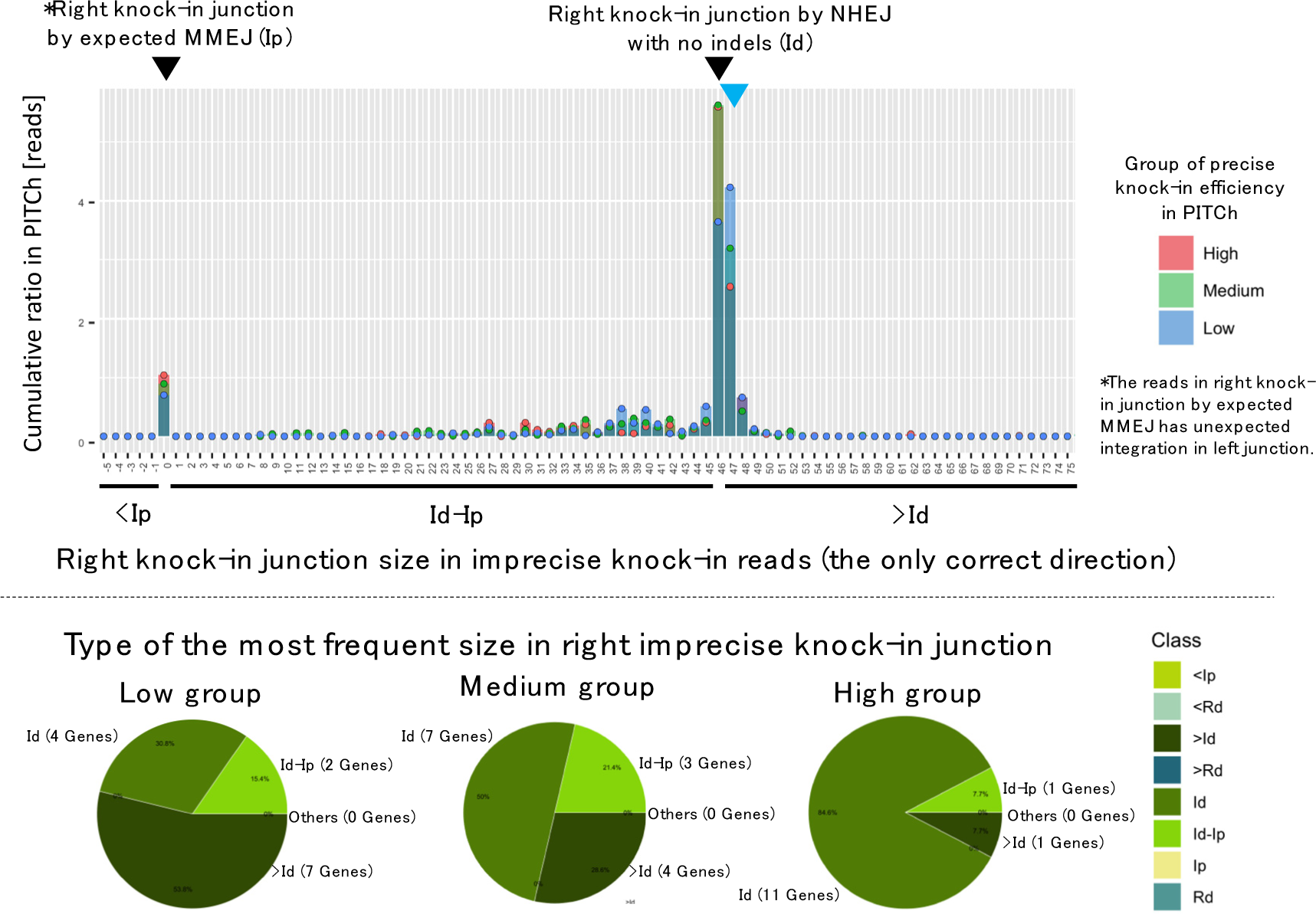
Distribution of right knock-in junction size (top) and the most frequent size in the right imprecise knock-in junction per group (bottom) in the imprecise knock-in class. The description is the same as that for Supplementary Figure 22. The blue arrowhead represents the undesirable feature of precise knock-in in PITCh. The top plot is saved as “[a- 1]Barplot_of_right_junction_of_indel_size_of_condition1_reads_by_condition1_group.png” in the output of MaChIAto Reviewer. The bottom plots are saved as “[c- 1]Piechart_of_max_indel_size_class_on_right_junction_in_condition1_reads_among _condition1_low_group.png,” “[c-1]Piechart_of_max_indel_size_class_on_right_junction_in_condition1_reads_among_condition1_medium_group.png,” and “[c- 1]Piechart_of_max_indel_size_class_on_right_junction_in_condition1_reads_among_condition1_high_group.png” in the output of MaChIAto Reviewer.

**Supplementary Figure 24.**
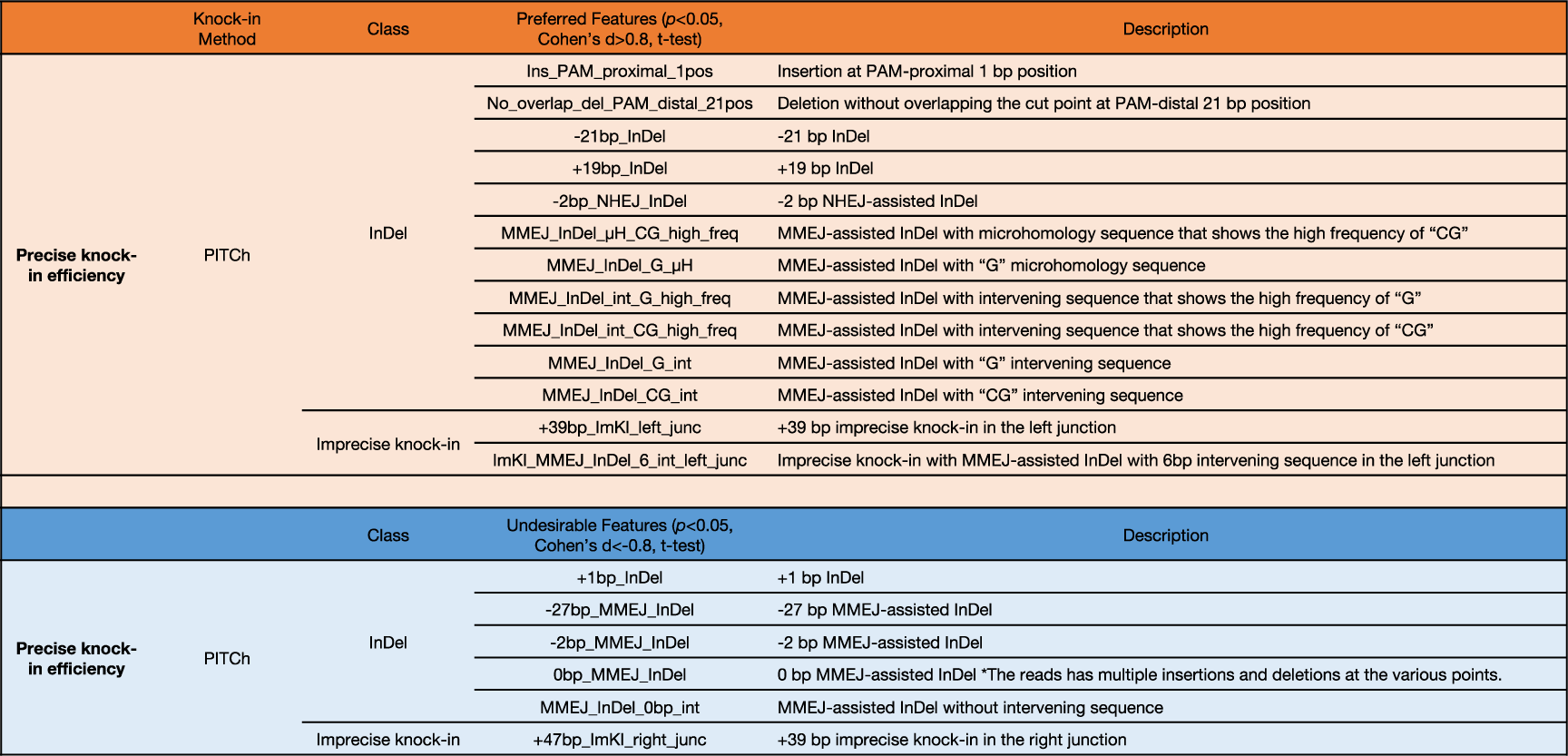
List of preferred mutation features and undesirable mutation features involved in the precise knock-in in InDel and imprecise knock-in classes that is generated in the PITCh knock-in sample.

The fact suggested that the activity of MMEJ knock-in depends on the endogenous mutation pattern. Moreover, we revised the alignment results of MaChIAto Aligner to confirm the mutation tendency in the genes that showed the best or worst precise knock-in efficiency (Supplementary Fig. 25, 26). There are plenty of significantly preferred features in the top ten frequent reads of the top genes (Supplementary Fig. 25). However, the most frequent reads of the genes that showed the worst efficiency had a +1-bp InDel in the PAM-distal position, otherwise, there was no preferred feature in the top ten frequent reads of the genes that presented inefficient knock-in results (Supplementary Fig. 26). It is important to note that the top genes had a G base on the 5′ end of the cut site because the G base is known for preventing a +1- bp InDel at the PAM-distal position^12, 20^. The +1-bp InDel at the PAM-distal position might be a crucial factor in deciding the outcome of MMEJ-assisted knock-in (Supplementary Fig. 24).

**Supplementary Figure 25.**
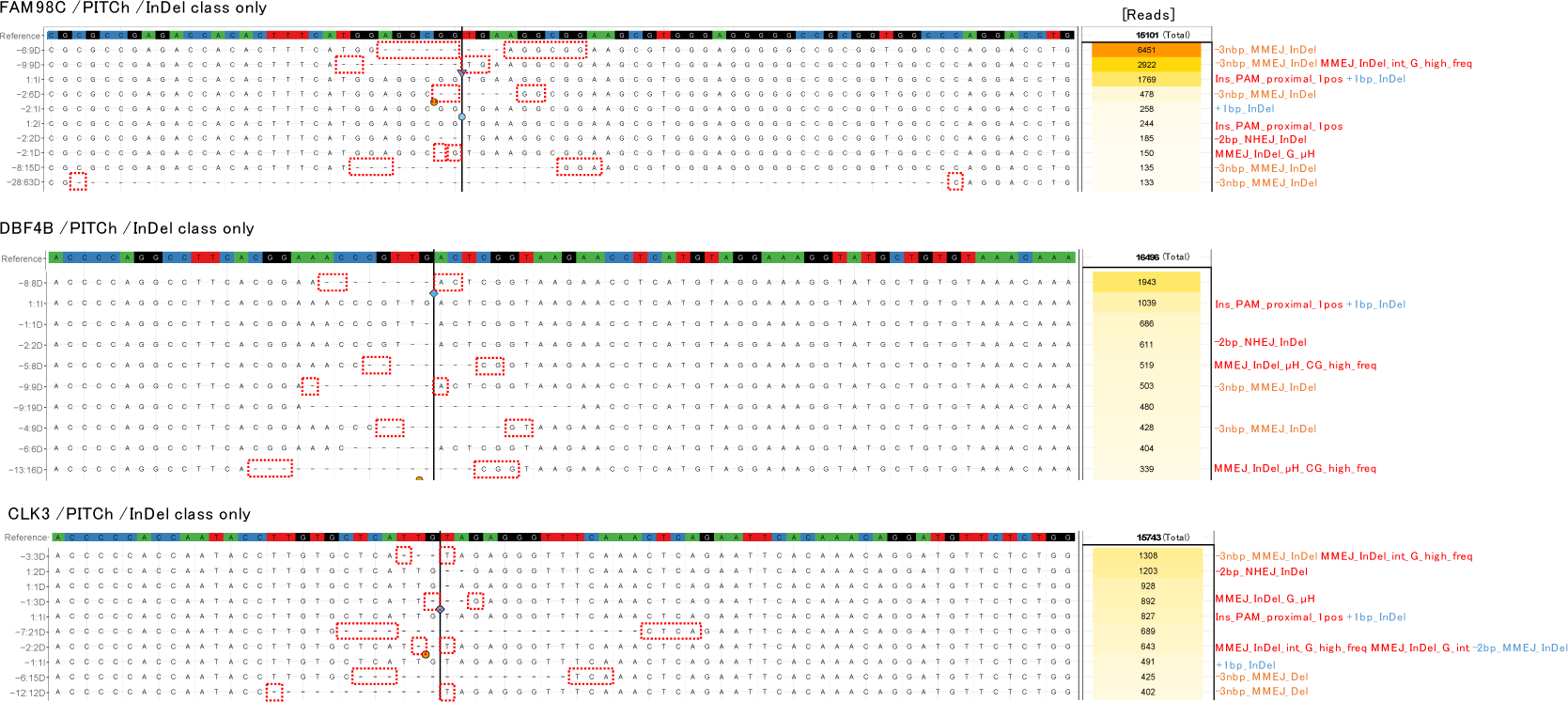
The ten most frequent mutations in the top three genes regarding the rate of precise knock-in in the InDel class. The numbers located beside the alignment map represent read number. The red and blue labels indicate the significantly preferred and undesirable features of precise knock-in in PITCh, respectively. The multiples of -3-bp MMEJ-assisted InDels (-3nbp_MMEJ_Del) represent the orange label. The plots are saved as “result/[1]comparison_with_original_crispresso_data/[Ⅱ]machiato_alignment_data/[A c]machiato_nhej/aligned.variants.pdf” in each of the output of MaChIAto Aligner.

We investigated the effect of LoAD against the pattern of reads in the InDel and imprecise knock-in classes (Supplementary Fig. 27). The distribution of InDel sizes in reads of the InDel class was not changed very much. The size distributions of MMEJ-assisted and NHEJ-assisted deletions were also not modified dramatically. However, in reads of the imprecise knock-in class, the right knock-in junction of the precise integration increased and the tendency of NHEJ-assisted integration with +1-bp InDels was decreased. These observations suggested that LoAD could modify the mutation pattern between the endogenous cut site and the knock-in donor; however, the mutations in the endogenous cut site could less be interfered with by LoAD.

**Supplementary Figure 26.**
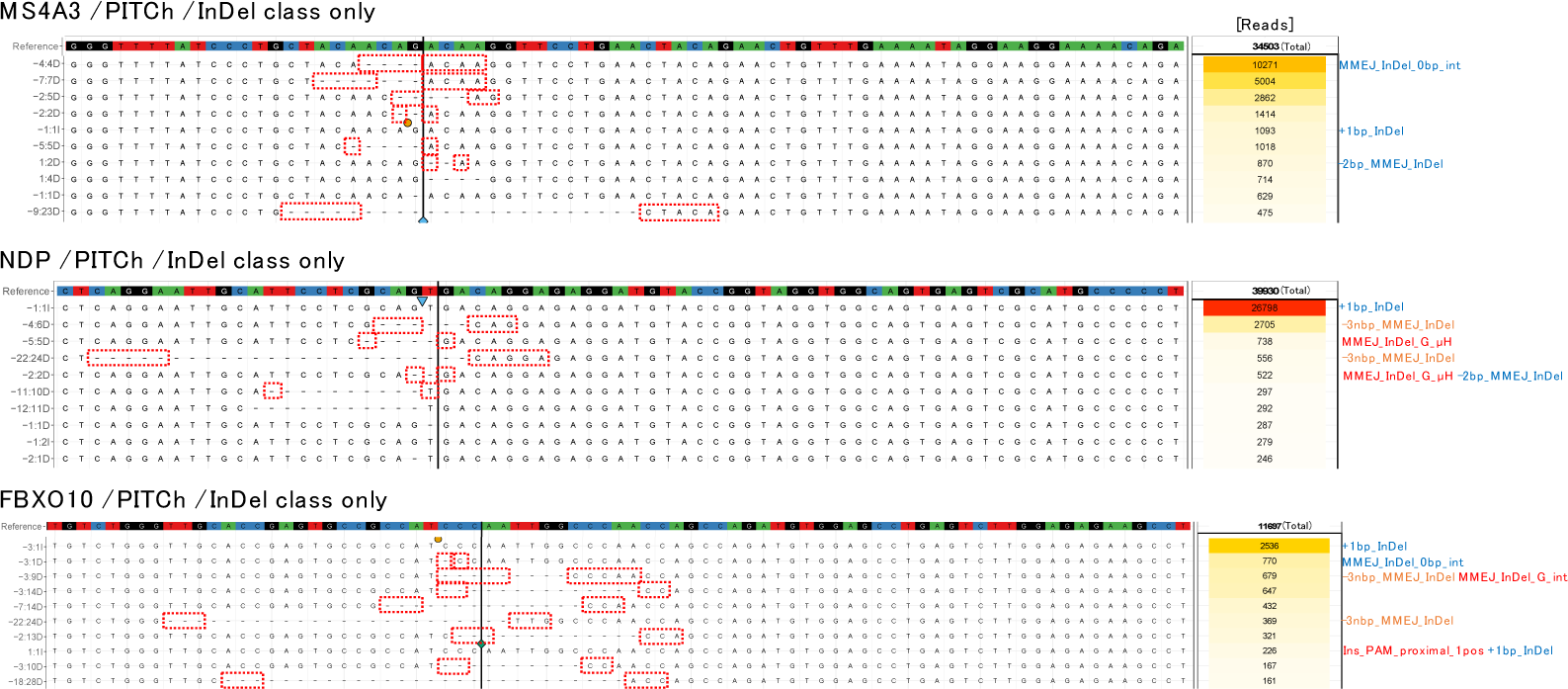
The ten most frequent mutations in the bottom three genes regarding the rate of precise knock-in in the InDel class. The numbers located beside the alignment map represent read number. The red and blue labels indicate the significantly preferred and undesirable features of precise knock-in in PITCh, respectively. The multiples of -3-bp MMEJ-assisted InDels (-3nbp_MMEJ_Del) represent the orange label. The plots are saved as “result/[1]comparison_with_original_crispresso_data/[Ⅱ]machiato_alignment_data/[A c]machiato_nhej/aligned.variants.pdf” in each of the output of MaChIAto Aligner.

**Supplementary Figure 27.**
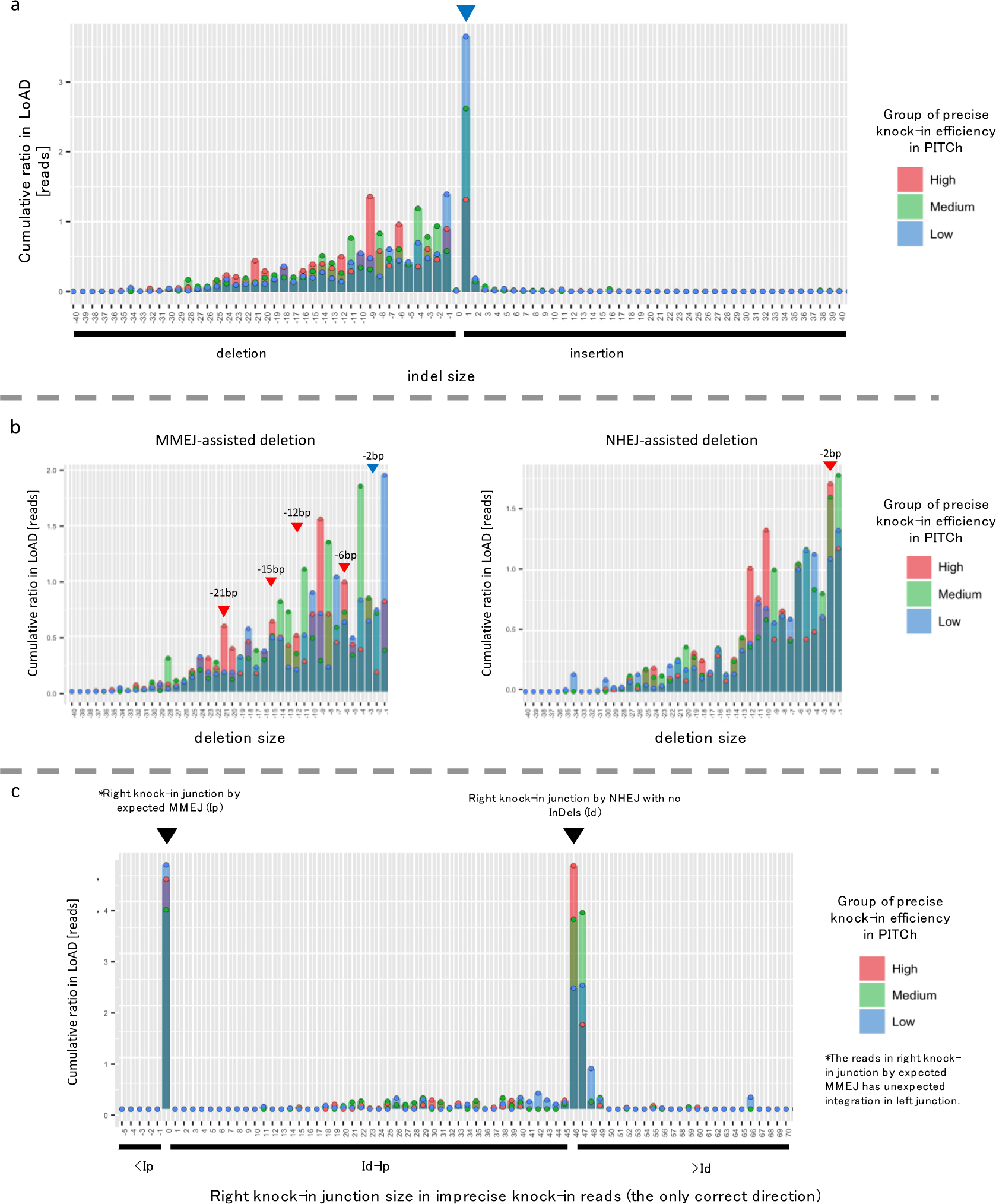
Distribution of InDel size (a), distributions of deletion size by MMEJ-assisted mutation (b, left) and NHEJ-assisted mutation (b, right), and distribution of right knock-in junction size (c) in LoAD. The group category is about the precise knock-in rate in PITCh. The reads of LoAD are plotted as the distributions according to this category. The red and blue arrowheads represent the preferred and undesirable features of precise knock-in in PITCh, respectively. The graph is helpful to check the change between PITCh and LoAD. The top plot is saved as “[b- 4]Barplot_of_mutation_indel_size_of_condition2_reads_by_condition1_group.png” in the output of MaChIAto Reviewer. The center plots are saved as “[j-4]Barplot_of_mutation_MMEJ- mediated_indel_size_of_condition2_reads_by_condition1_group.png” and “[k- 4]Barplot_of_mutation_NHEJ- mediated_indel_size_of_condition2_reads_by_condition1_group.png” in the output of MaChIAto Reviewer. The bottom plot is saved as “[a- 4]Barplot_of_right_junction_of_indel_size_of_condition2_reads_by_condition1_grou p.png” in the output of MaChIAto Reviewer.

To support the discussion, we performed the additional profiling of mutation pattens of the Cas9-sgRNA only (knock-out) samples using MaChIAto. The high, medium, and low groups were generated in terms of the precise knock-in efficiency in PITCh knock-in. We confirmed that 10 of 20 significant mutation patterns were the same as the mutation types that were significant in the above knock-in analysis (Supplementary Fig. 28). The fact implied that many mutation features affecting knock- in efficiency emerged from the existence of donor vector. The consistent mutations could be a native pattern of the (epi-)genomic context of the target site.

**Supplementary Figure 28.**
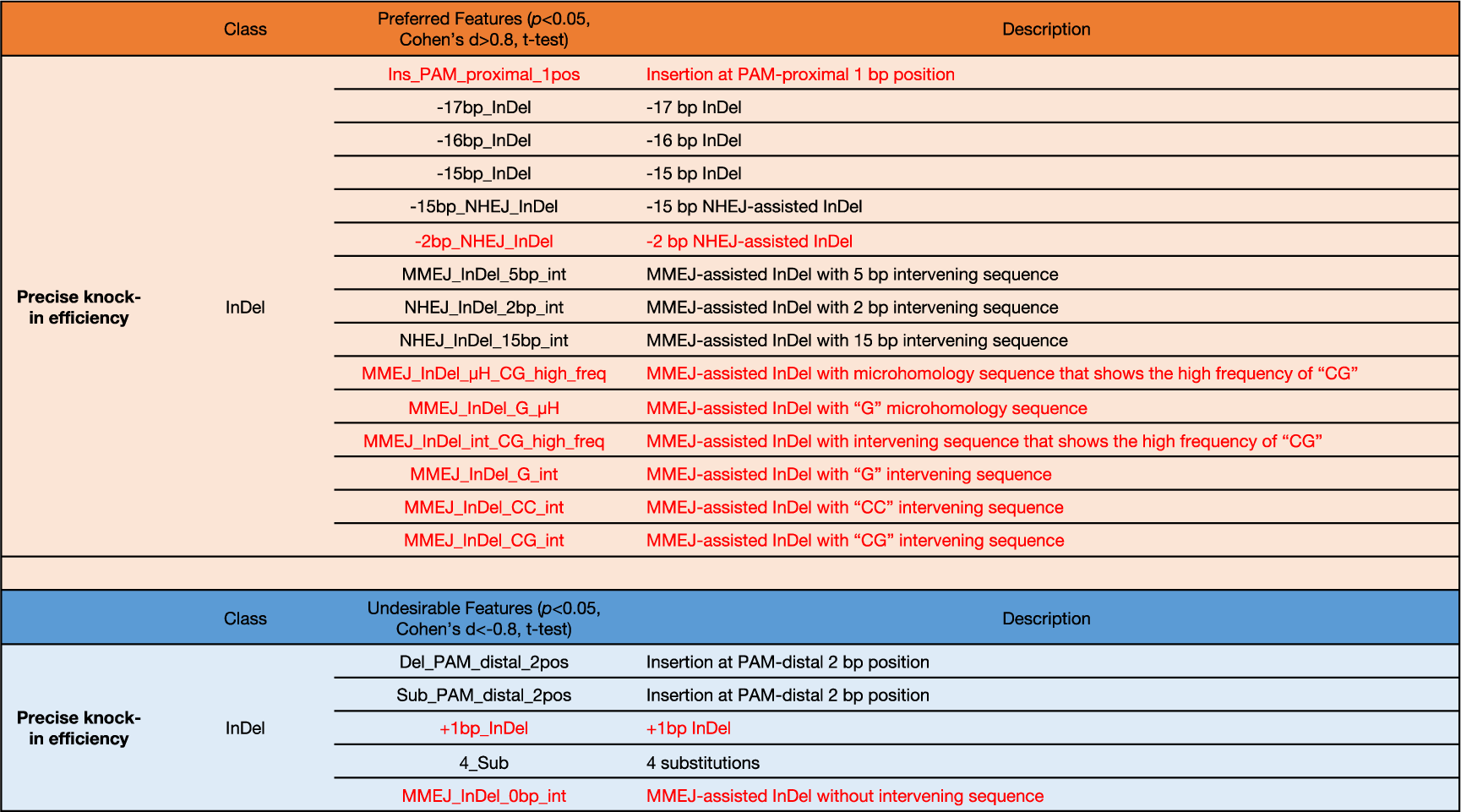
List of preferred mutation features and undesirable mutation features involved in the precise knock-in in InDel class generated in the Cas9-sgRNA only (knock-out) sample. The features show significance in the *t*-test (*p* < 0.05 and Cohen’s d > 0.8 in the preferred features, and *p* < 0.05 and Cohen’s d < -0.8 in the undesireble features) that compares the gene group of high precise knock-in efficiency in PITCh knock-in with the low group. The red-colored features are common with Supplementary Figure 24.

As described above, MaChIAto can provide a practical and comprehensive profile to gain a deeper understanding of knock-out and knock-in efficiency. We used PITCh and LoAD knock-in methods in this analysis; however, MaChIAto can be applied to other short homology-based methods and beyond^10^.

#### Supplementary Note 7 - Practical example for MaChIAto: a series of knock-out targeting 40 gene loci in HEK293T cells

Along with the investigation of the (epi-)genomic features affecting knock-in efficiency using the data from a series of targeting at 40 gene loci (Supplementary Note 6), we also performed a series of targeting at 40 gene loci without donor (Cas9-sgRNA only). MaChIAto also accommodates knock-out analysis to find features controlling the editing efficiency in the same way as the knock-in analysis by MaChIAto (Supplementary Note 6). Here, we used the in-house data of a series of targeting at 40 gene loci as an input of the MaChIAto Analyzer to investigate the features affecting the editing efficiency of sgRNAs.

As a result, MaChIAto Analyzer found some significant features. In the in-house data of a series of 40 gene loci, the change of enthalpy, the minimum energy of energy, and TA motif had a positive effect on the editing efficiency, while disrupt duplex energy and CG motif showed a negative one (Supplementary Fig. 29). Interestingly, TT di-nucleotide frequency was not a significant feature, although it has been reported as a critical feature in the lentiviral CRISPR screen^13, 44, 45^. When we used the sgRNA activity data of ribosomal genes compiled by Xu and colleagues^46^ as the input of the MaChIAto analyzer, the tool showed that the frequency of TT di-nucleotide is significant (*p* < 0.001) (Supplementary Fig. 30). We assume that there are two probable reasons causing the conflict. First, sequence diversity is not enough to evaluate the feature. In our data, the range of TT number was 0–7, although the theoretical range is 0–19. Second, the TT effect may be specific in the lentiviral CRISPR screen. CRISPR screen is an efficient method to get numerous sgRNA activity data, although the methodology has some sources of error^47^.

**Supplementary Figure 29.**
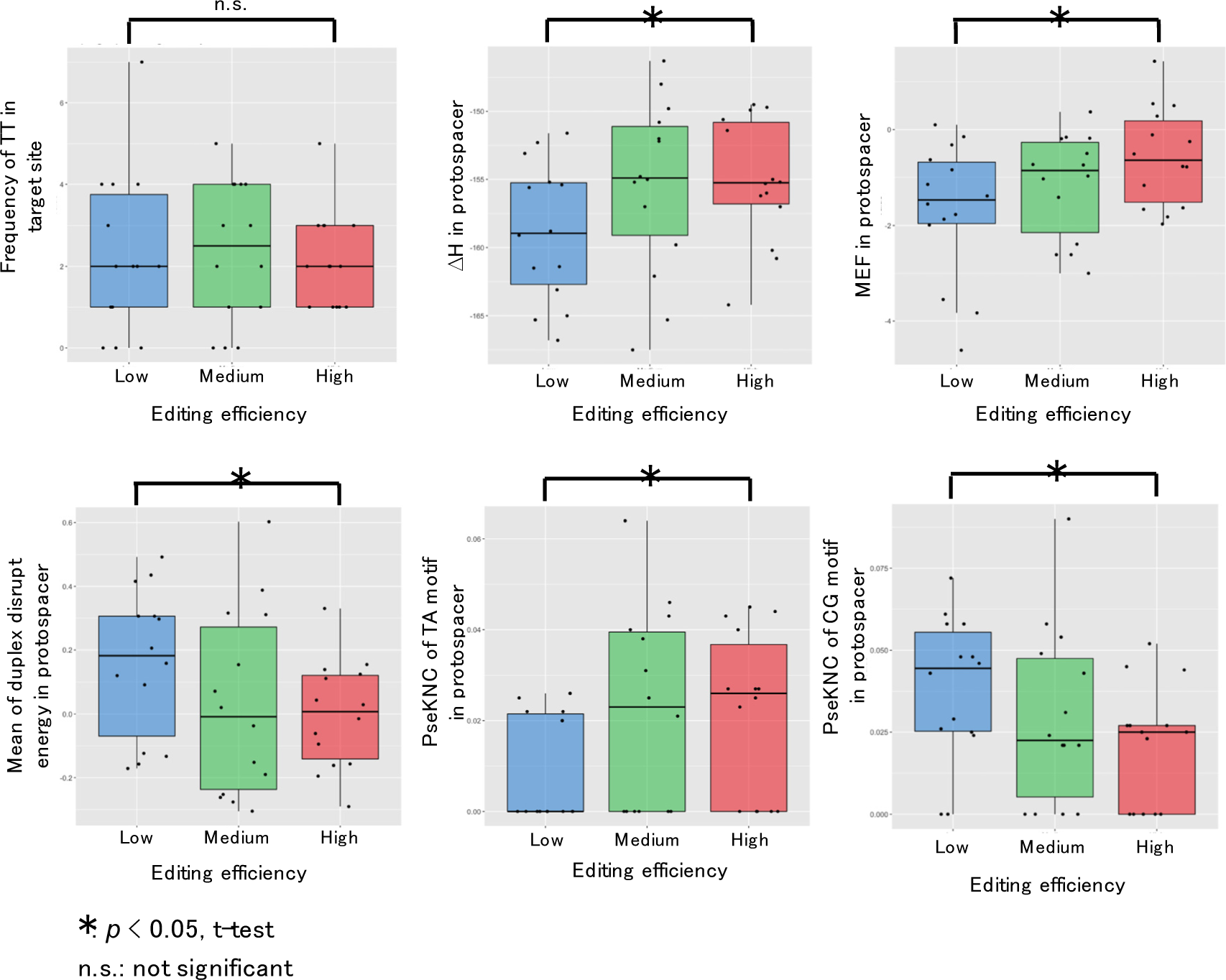
Comparison of features among the high, medium, and low groups in terms of the editing efficiency of a series of knock-out at 40 gene loci in HEK293T cells. The values of each feature are summarized in the box plots. The plots are split by the high, medium, and low groups regarding the editing efficiency. Each dot represents one value of the targeting site. The asterisks and n.s. mean significant at *p* < 0.05 and not significant in the *t*-test, respectively.

Next, we analyzed the mutation patterns using MaChIAto Reviewer. In knock- out analysis mode, it makes the high, medium, and low groups in terms of the editing efficiency. We found some mutation types that showed a significant difference between the low and high groups (Supplementary Fig. 31). The result suggested that the G and GA DµH-dependent MMEJ might contribute to the efficient InDel formation.

**Supplementary Figure 30.**
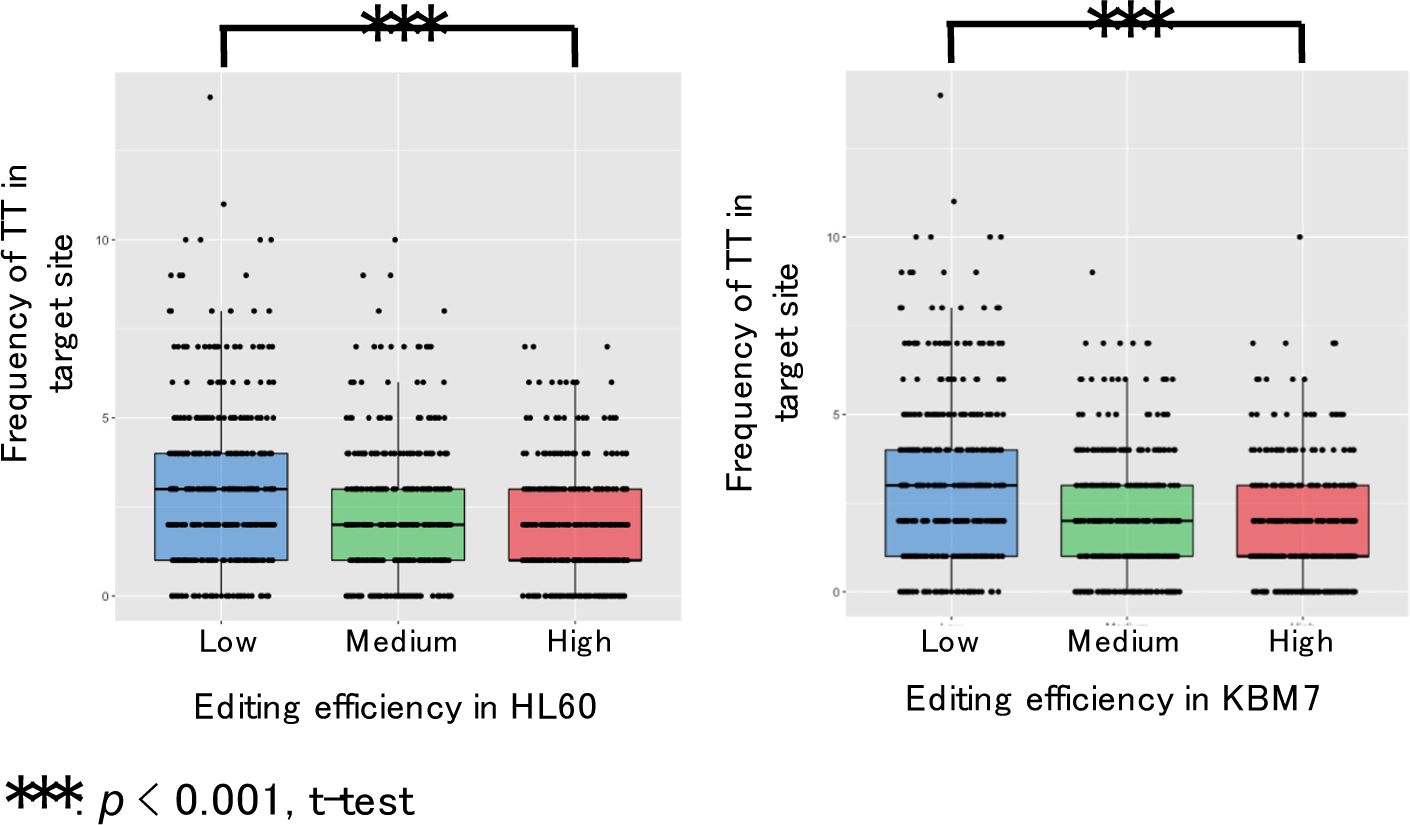
Comparison of TT frequency among the high, medium, and low groups in terms of the editing efficiency of ribosomal genes of Xu *et al*., 2015^46^ (n=1,169). The values of each feature are summarized in the box plots. The plots are split by the high, medium, and low groups in terms of the editing efficiency. Each dot represents one value of the targeting site. The triple asterisks mean significant at *p* <0.001 in the *t*-test.

**Supplementary Figure 31.**
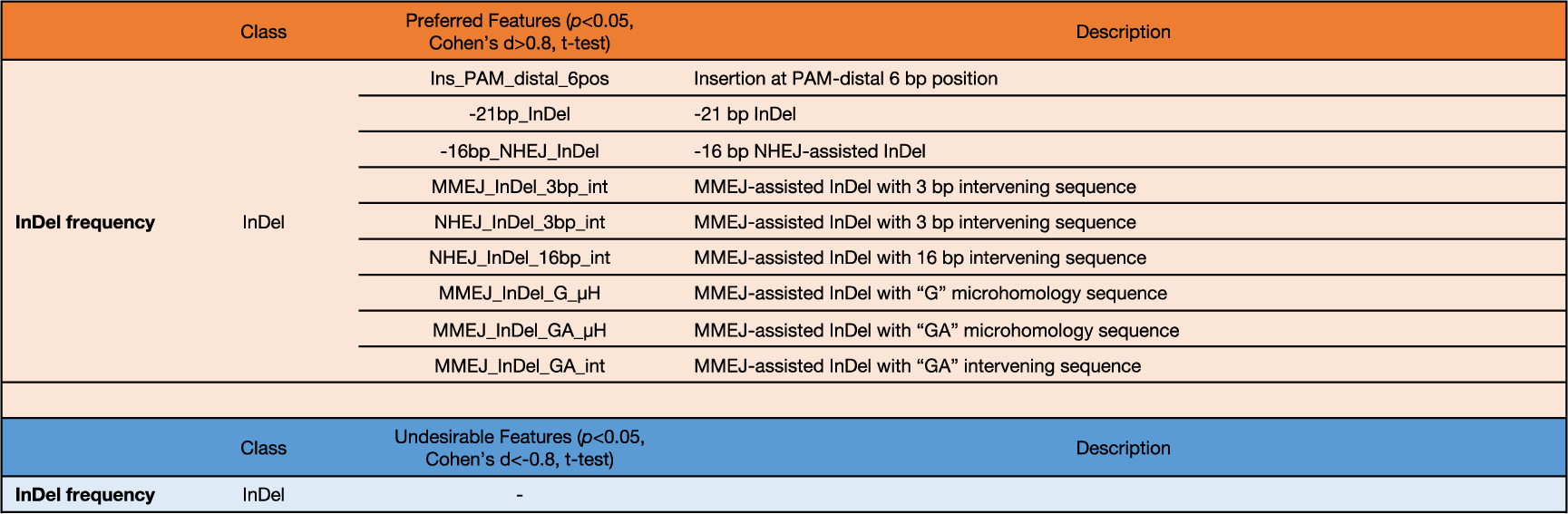
List of preferred mutation features and undesirable mutation features involved in the editing efficiency in InDel class generated in the Cas9-sgRNA only (knock-out) sample. The features show significance in the *t*-test (*p* < 0.05 and Cohen’s d > 0.8 in the preferred features, and *p* < 0.05 and Cohen’s d < -0.8 in the undesireble features) that compares the gene group of high editing efficiency in knock- out with the low group.

This section shows the capability of knock-out analysis using MaChIAto, which can find significant features affecting the editing efficiency using plasmid vector-based in-house data and publicly available CRISPR screen data^46^. The result of the plasmid vector-based data conflicts the previous knowledge retrieved from lentiviral pooled screening. The reason seems to be a small sample size (n=40) or some technical problems^47^. The assessment of the difference will require a large amount and broad diversity of data on the editing efficiency using various experimental methods such as lentiviral transduction^44, 45^, plasmid vector-based transfection, and sgRNA injection *in vivo*^48^. MaChIAto Analyzer can analyze any type of experimental data if researchers have the efficacy data and the target sites’ sequences. MaChIAto would be a common platform to find the significant features

#### Supplementary Note 8 - InDel prediction tool is helpful for knock-in prediction

The analysis of a knock-in series targeting 40 gene loci provided a lot of curious facts for the understanding of MMEJ-assisted knock-in. We found the endogenous InDel tendency directly leaded to precise knock-in tendency (Supplementary Note 6). Notably, the sequence features such as a G base on the 5′ end of the cut site can be used to estimate the InDel tendency. However, it is still challenging to utilize the finding for knock-in design because we cannot always find the sequence features for the prediction. To solve this issue, we investigated whether the existing tools for InDel prediction helped researchers to predict that the InDel tendency on a template-free knock-out is appropriate for a knock-in. We predicted the endogenous variants using the tools such as inDelphi^20^ and FORECasT^12^, and then the distribution of estimated variants was compared with the real distribution of untreated, Cas9-sgRNA only, PITCh and LoAD samples (Supplementary Fig. 32). The corresponding combination (i.e., same genes) showed more substantial similarity in Cas9-sgRNA only, PITCh, and LoAD samples. We summarized the result as a box plot by the corresponding combination and non-corresponding combination (i.e., different genes) (Supplementary Fig. 33). The box plot proved the similarities were significant in these samples. Thus, we can propose that the researchers can predict the precise knock-in efficiency of PITCh and LoAD using the InDel prediction tools even if they do not have the information on InDel distribution in advance.

**Supplementary Figure 32.**
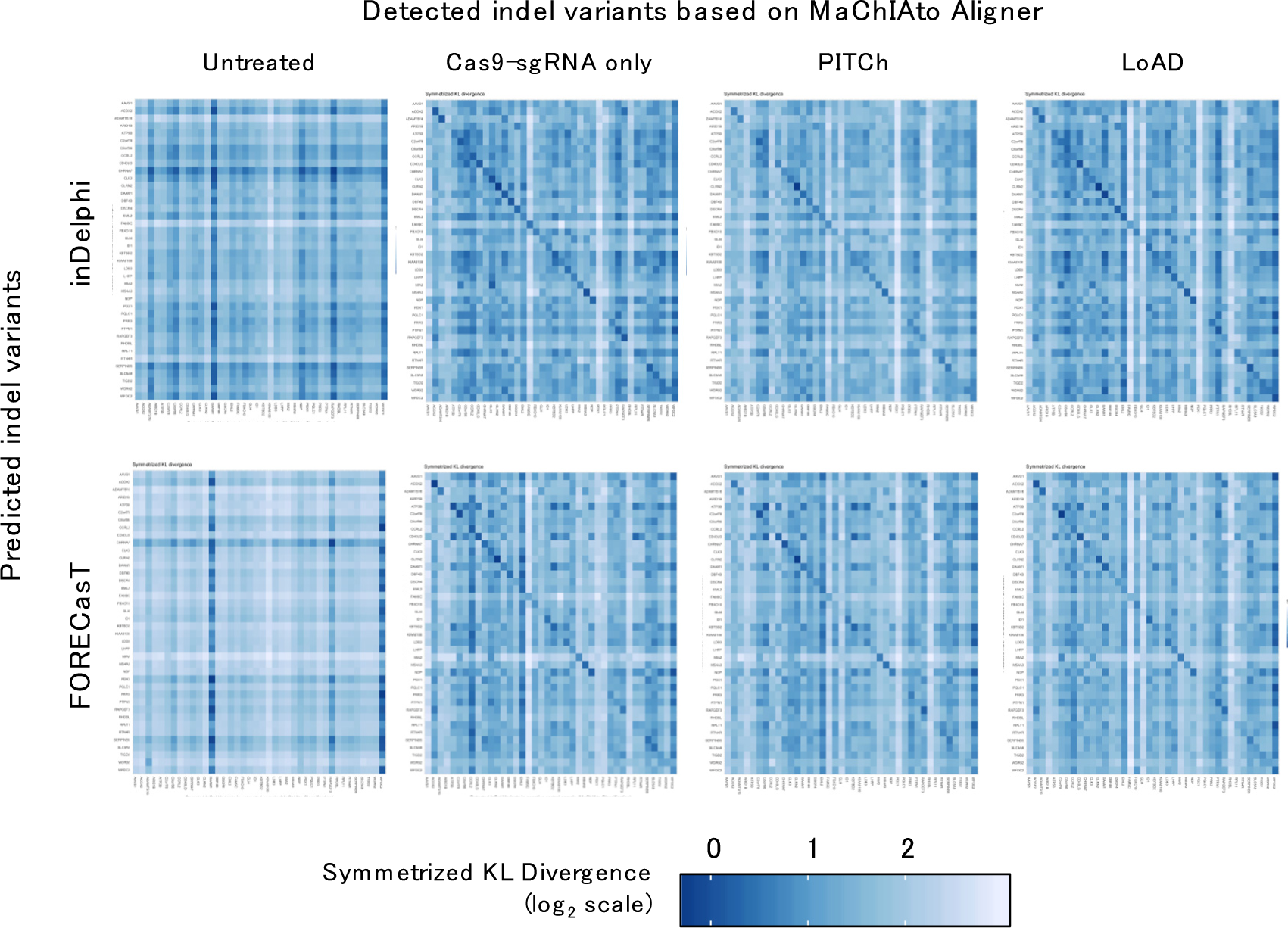
Comparison between the detected InDel variants which MaChIAto Aligner provided and the predicted variants by inDelphi or FORECasT. Symmetrized KL divergence (blue to white color scale) between the detected InDel variants of a gene in a sample type (x-axis) and the predicted variants of a gene (y-axis). The genes are arranged so that the same genes (the corresponding combination) can be located on a diagonal from the top left to bottom right. Sample types are untreated, Cas9-sgRNA only, PITCh, and LoAD. The lower Symmetrized KL divergence represents a higher similarity.

#### Supplementary Note 9 – Analysis of Prime Editing samples

We finally tested whether the tool could apply the sample of other precise editing methods because most of the precise editing system has left/right homologous component (HC) like homology arms. To check it, we utilized MaChIAto to analyze the amplicon sequencing data of Prime Editing^5^. Prime Editing is a reverse-transcriptase-based CRISPR editing method in which an engineered M-MLV reverse-transcriptase fused with Cas9 nickase (Prime Editor) introduces a nick on the editing strand and then starts reverse transcription to generate the desired genomic sequence. For the precise modification, Prime Editing system harnesses a 3′-extended sgRNA, prime editing guide RNA (pegRNA), to provide a binding site against the 3′-end of the nicked site (primer binding site) and act as a template for reverse transcription (RT template). In the high-efficiency version of Prime Editing system, called PE3, the system additionally introduces a nick on the non-edited strand using an additionally expressed sgRNA (nicking gRNA; ngRNA) along with Prime Editor so that the editing efficiency can increase by the preferential removal of the sequence on the non-edited strand. According to the previous report^8^, the PE3 achieved the precise modification with typically 20–50% editing efficiency in the HEK293T cells.

To analyze the Prime Editing system using MaChIAto, we considered the sequences of primer binding site and RT template as left and right HC, respectively (Supplementary Fig. 34), and then built an optimized mode of MaChIAto for Prime Editing. Our prominent concept of the mode is a “wide perspective.” Prime Editing utilizes two CRISPR targeting sites on both strands, which can introduce InDels by double nicking. Therefore, we must prove that the low sequencing coverage and the severe mutagenesis do not occur outside the editing site. However, most TASA tools only focus on the editing site and do not sufficiently take care of the other region. We implemented the Prime Editing mode of MaChIAto so that it can survey the editing site (pegRNA binding site) and the additional target site (ngRNA binding site) using the indicator method (see Methods) and pattern matching. To demonstrate the capacity, we analyzed the amplicon sequencing dataset of ref 5 using CRISPResso2 and MaChIAto (Supplementary Table 5). The dataset has diversity in terms of sequencing coverage. Some sequencing data have low coverage on the ngRNA binding site (Supplementary Fig. 36, all samples). First, as a positive control, we analyzed the data whose coverage at the target site and ngRNA binding site is enough to analyze (Supplementary Fig. 35, a). The estimation of editing rate was similar to that of CRISPResso2 (Supplementary Fig. 35, b: the difference of perfect editing was 4.13%). In contrast, when the data whose coverage on the ngRNA binding site is less than that of cut site (Supplementary Fig. 36, all samples), MaChIAto successfully removed the sequence having lack and mutations on the ngRNA binding site from perfect editing class (Supplementary Fig. 37, all samples: the differences of perfect editing were 14.12%, 16.96%, 79.86, and 37.34%). MaChIAto is to put it into an “Ambiguous editing” class or “Other” class if a read does not have a sequence at the ngRNA binding site, because it is difficult to discuss the qualification of InDels without the information on the ngRNA binding site, which mainly causes the unexpected InDels in the PE3 system. MaChIAto classification is more accurate compared to CRISPResso2 in even Prime Editing. However, we do not mean that CRISPResso2 does not have the advantage compared with MaChIAto (Classifier) and a mapping-based tool like a BWA MEM- CrispRVariants. For example, the BWA MEM could not map reads at the *HEK3* locus in the substitution editing #2 (Supplementary Fig. 36), although we observed the target region’s sequence in the reads using ClustalW alignment^49^ (Supplementary Fig. 38). CRISPResso2 did not miss the reads and then classified most of them into “Perfect editing” or “Unmodified” classes (Supplementary Fig. 37). MaChIAto did not mostly count these classes because most of them lacked the sequence at the ngRNA binding site (Supplementary Fig. 38). We propose that the user of amplicon sequencing should use CRISPResso2 as the first rough tool that detects reads to the fullest extent, and then they get the more exact and detailed classification and profile of the Prime Editing using MaChIAto.

**Supplementary Figure 33.**
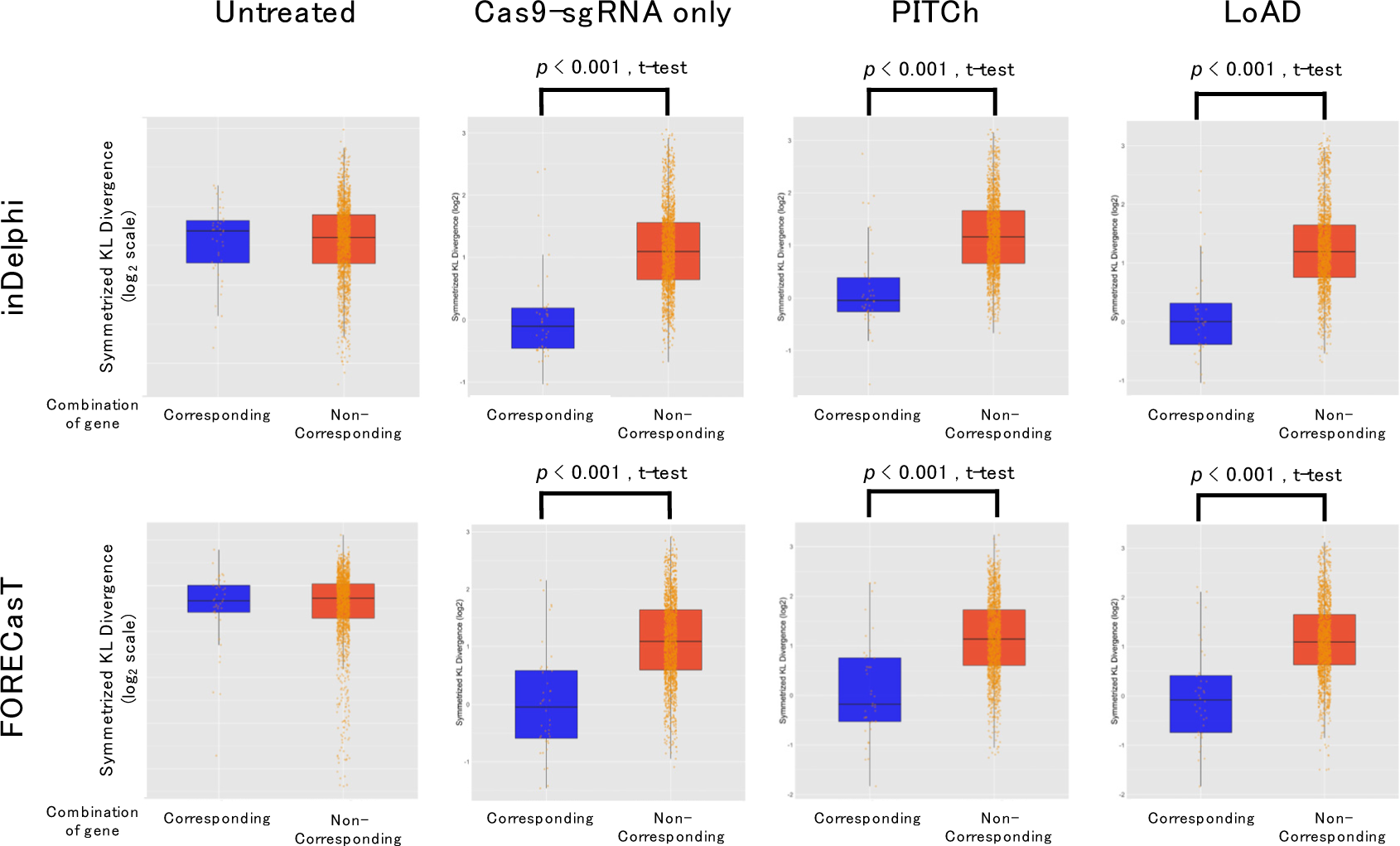
The comparison between the values of the corresponding set and values of the non-corresponding set. Symmetrized KL divergence (log2 scale) is shown on the y-axis.

**Supplementary Figure 34.**
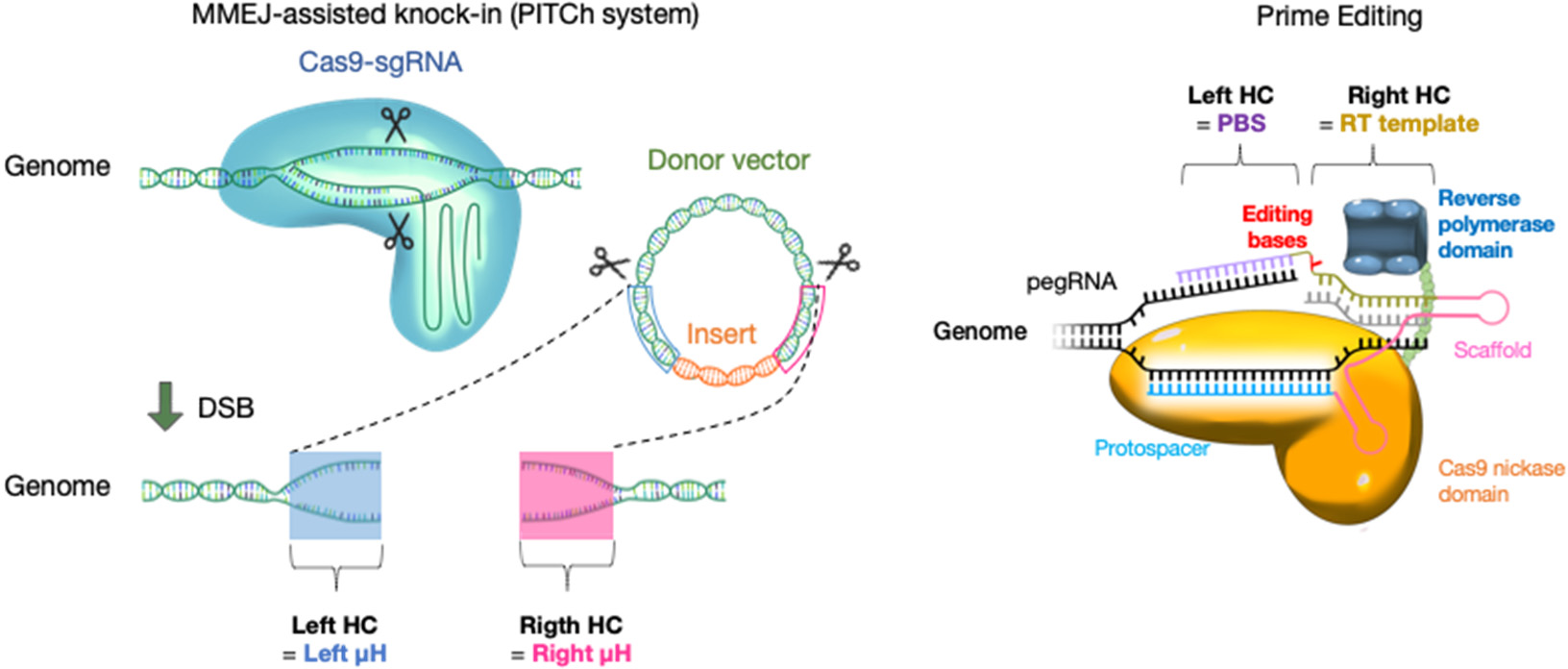
Schematic of the MMEJ-assisted knock-in (PITCh system) and Prime Editing. Left. PITCh system harnesses homologous components (HCs) to induce the MMEJ between the cut sites of donor and target locus. Right. Prime Editing utilizes the HCs as the primer and template in reverse transcription.

MaChIAto cannot only care about the ngRNA binding site but also checks the conservation of the reverse-transcript (RT) derived from the RT template. MaChIAto classifies the reads with an imperfect RT into the “Ambiguous editing” class even if the reads have the expected editing bases without apparent insertion and deletion. When the reads with reverse-transcription phase-dependent mutation are not eligible for the perfect editing reads, MaChIAto could recognize the reads with only the expected editing bases as perfect editing reads in the best case (Supplementary Fig. 39, bottom), whereas CRISPResso2 counted the reads with the unexpected reverse-transcription phase-dependent mutation and obvious InDels among the “Perfect editing” class (Supplementary Fig. 39, top). When we defined the misclassification rate as the rate of the reads with the unexpected mutations in the perfect editing class, the misclassification rates of MaChIAto were significantly lower in comparison with the results of CRISPResso2 in the *RNF2*, *RUNX1*, and *VEGFA* loci (Supplementary Fig. 40).

**Supplementary Figure 35.**
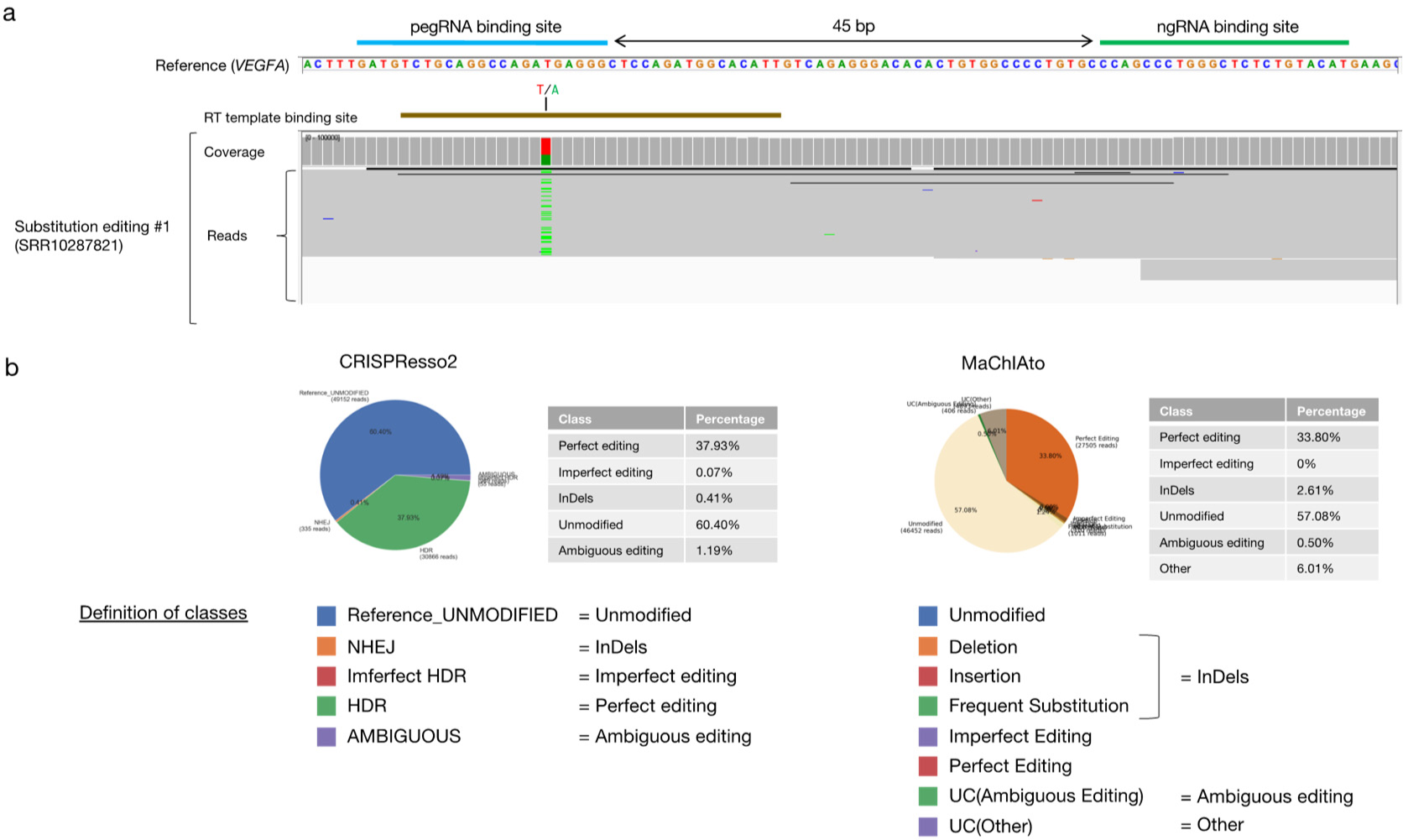
Integrative Genomics Viewer (IGV) visualization of a fragment of the *VEGFA* gene on chromosome 6 encompassing the target sequence comparing the unmodified *VEGFA* (Reference) and the amplicon sequencing data of substitution editing #1 (SRR10287821) (a), and the classification result processed with CRISPResso2 and MaChIAto (b). The binding sites of pegRNA, ngRNA, and RT template are outlined in light blue, green, and brown, respectively. The range of coverage is from 0 to 100,000. The classification labels are renamed to compare each other.

**Supplementary Figure 36.**
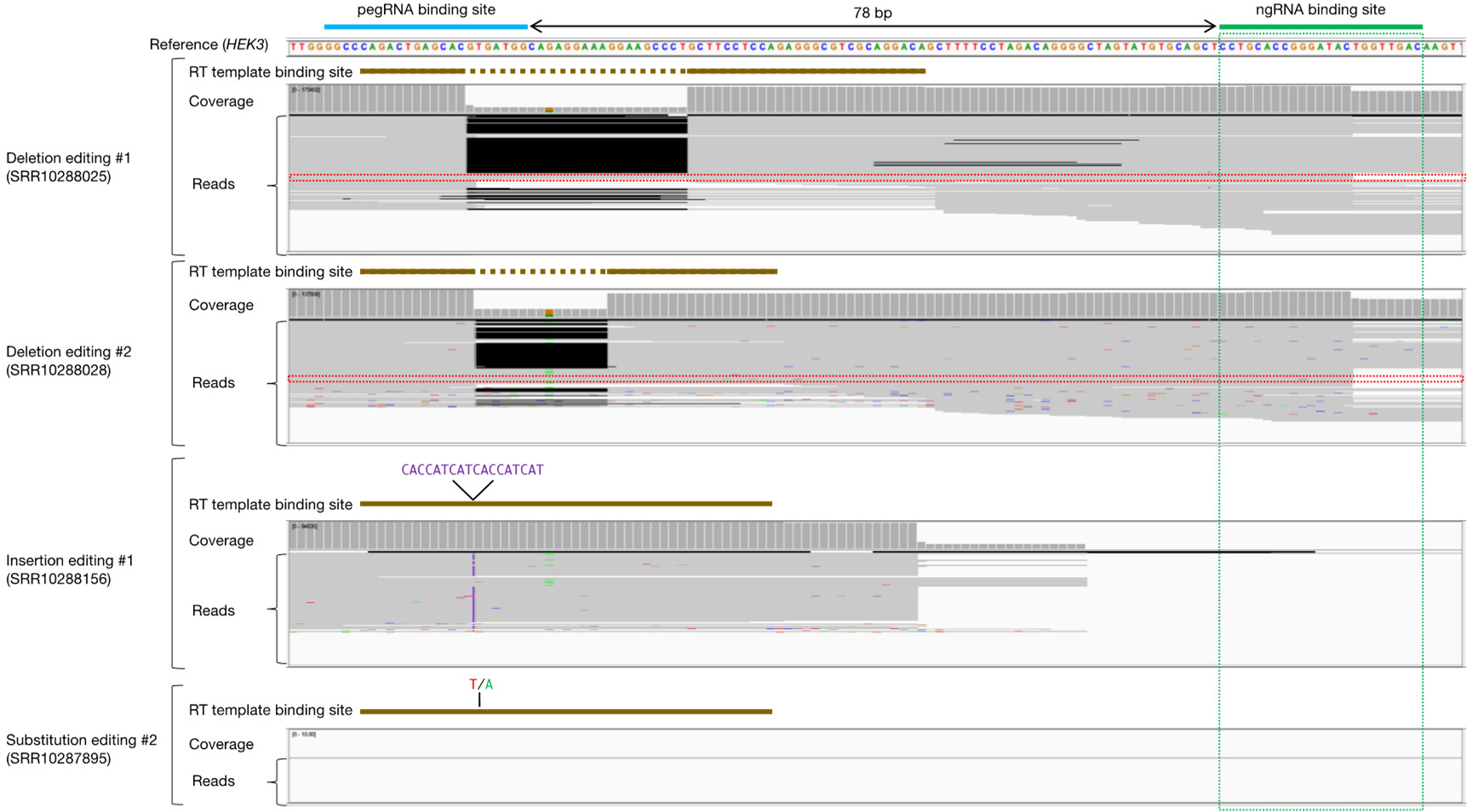
Integrative Genomics Viewer (IGV) visualization of a fragment of the *HEK3* gene on chromosome 6 encompassing the target sequence comparing the unmodified *HEK3* (Reference) and the amplicon sequencing data of deletion editing #1 (SRR10288025), deletion editing #2 (SRR10288028), insertion editing #1 (SRR10288156), and substitution editing #2 (SRR10287895). The binding sites of pegRNA, ngRNA, and RT template are outlined in light blue, green, and brown, respectively. The brown dotted lines indicate the excluded sequence in the RT template. The insertion sequence in the RT template represents the purple characters. The ranges of coverage are from 0 to 173,402, from 0 to 137,938, from 0 to 94,535, and from 0 to 10.00. The region of the binding site of ngRNA is highlighted with the green dotted box. The unmodified reads of deletion editing #1, #2 seem not to have the full sequence of the ngRNA binding site (red dotted box).

**Supplementary Figure 37.**
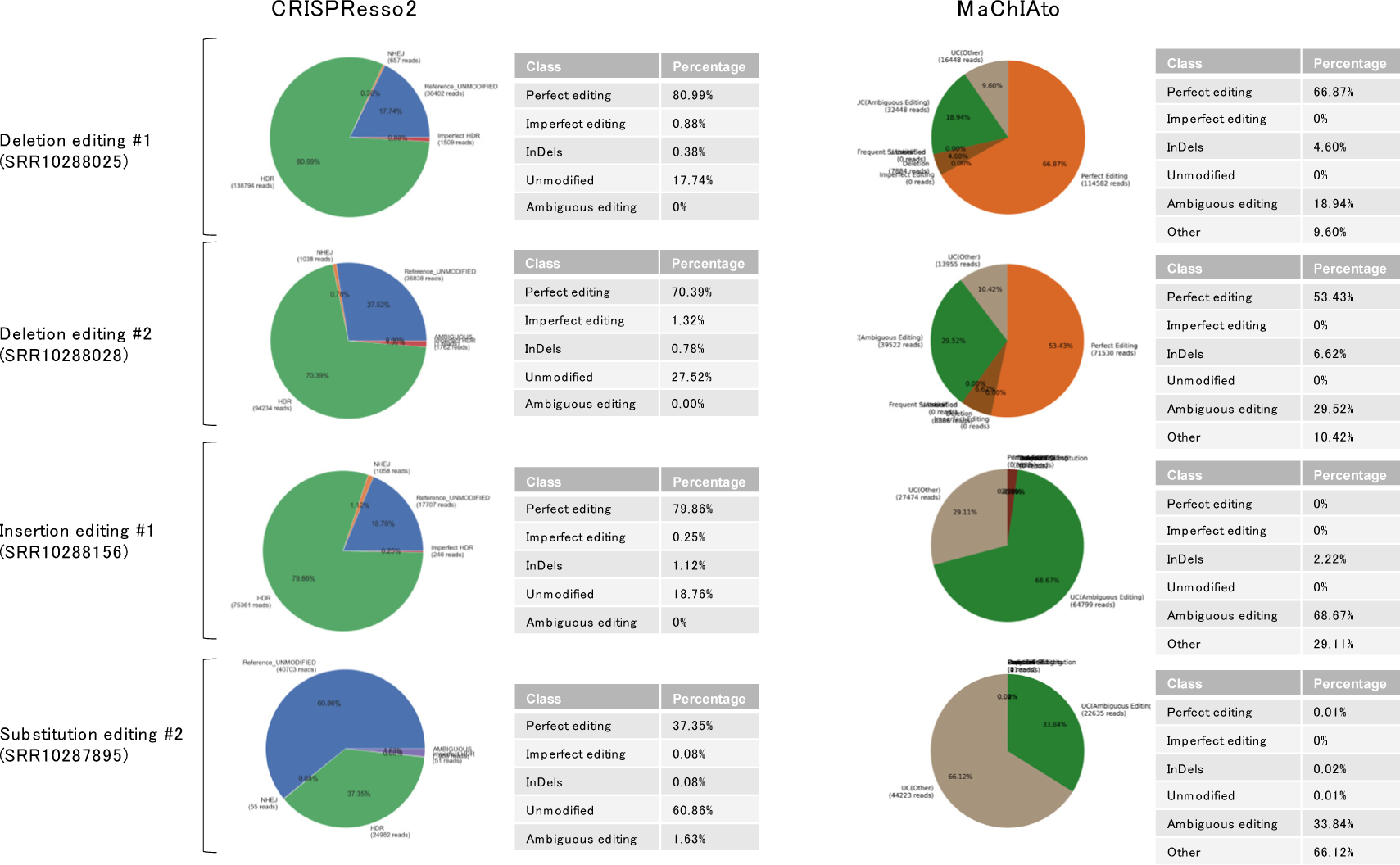
The classification result of amplicon sequencing samples of Supplementary Figure 36 processed with CRISPResso2 and MaChIAto. The classification labels are the same as Supplementary Figure 35. The MaChIAto results of insertion editing #1 and substitution editing #2 are incredibly different from the CRISPResso2 ones because the most reads of these samples are unclear sequences that did not have the ngRNA binding site.

**Supplementary Figure 38.**
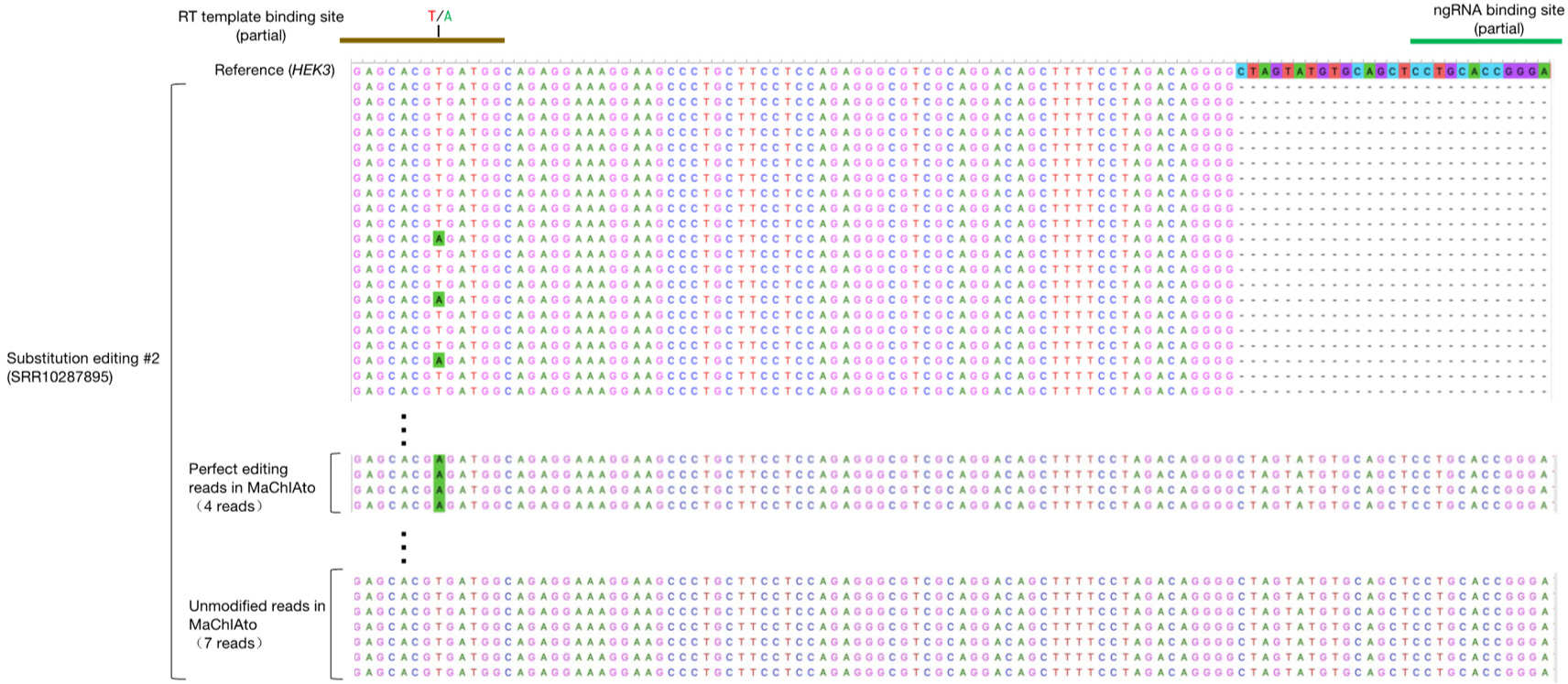
The multiple sequence alignment of substitution editing #2 processed with ClustalW. The green boxes within the RT template binding site represent the edited bases. The top of the sequence is the reference sequence of the *HEK3* amplicon sequence. The binding sites of ngRNA and RT template are outlined in green and brown, respectively. Most reads did not have the sequence of the ngRNA binding site. However, a few reads recognized as perfect editing or unmodified reads with MaChIAto have the ngRNA binding site.

**Supplementary Figure 39.**
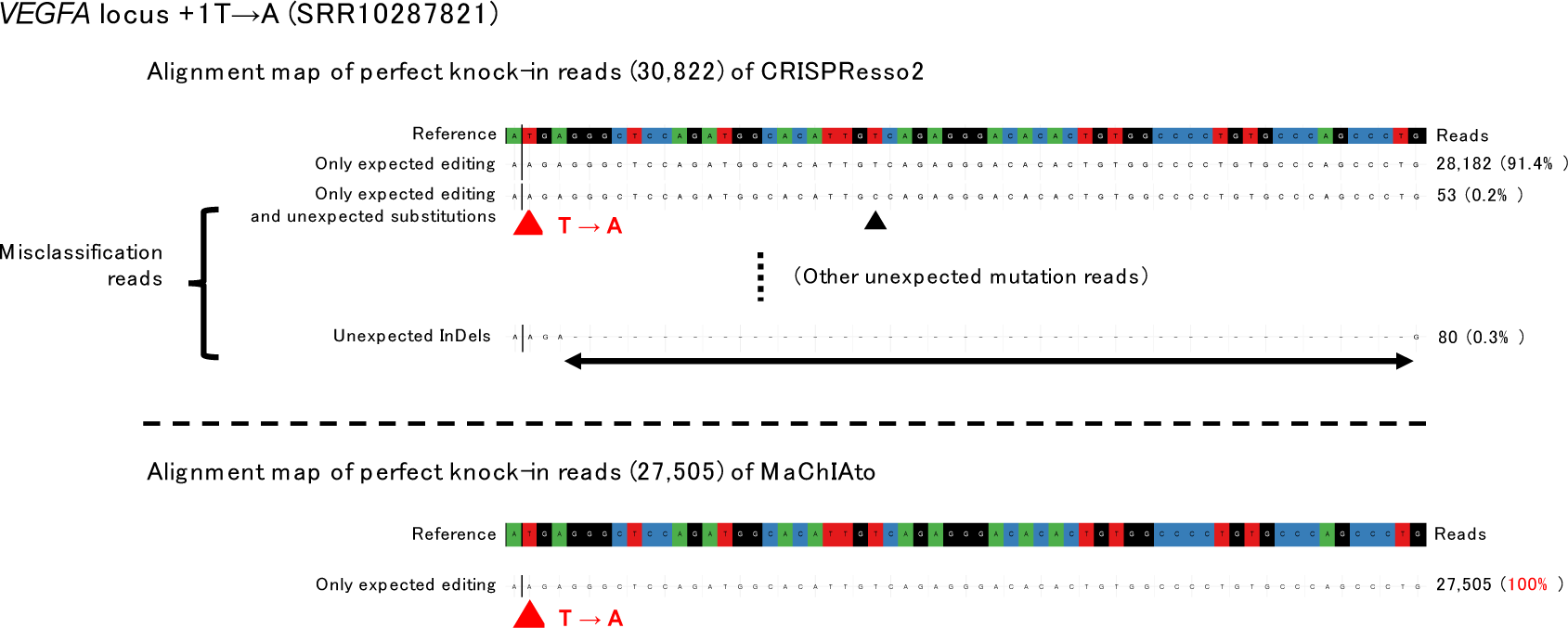
The Alignment map of the perfect editing class of CRISPResso2 and MaChIAto in *VEGFA* +1TA. Top. Perfect editing class of CRISPResso2 had not only the expected sequence but also unexpected mutation reads. Bottom. The perfect editing class of MaChIAto included only the expected editing sequence. These alignments are based on the BWA MEM-CrispRVariants (local alignment).

**Supplementary Figure 40.**
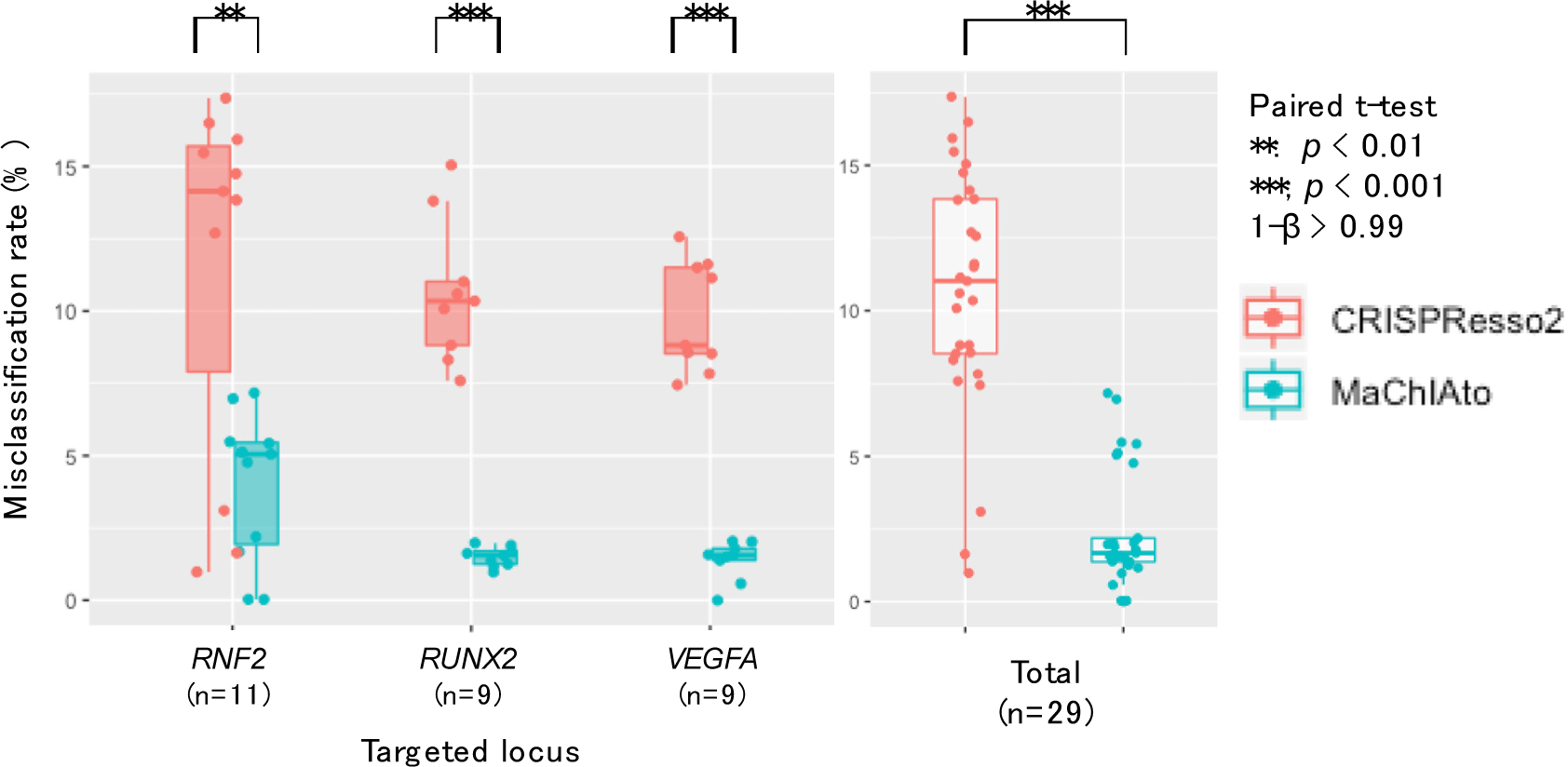
The rate of misclassification in the *RNF2*, *RUNX1*, and *VEGFA* loci. The red and blue points indicate the rate of misclassification in CRISPResso2 and MaChIAto, respectively. The misclassification rate was considered as the rate of reads that has unexpected mutations.

Furthermore, we aggregated the data and then classified them by the size of variants. Notably, all variants in the perfect editing class of MaChIAto are the same length as the expected editing sequence, whereas those of CRISPResso2 are not the same in some variants. The result indicates that MaChIAto could ideally eliminate the obvious InDel reads from the perfect editing class (Supplementary Fig. 41).

**Supplementary Figure 41.**
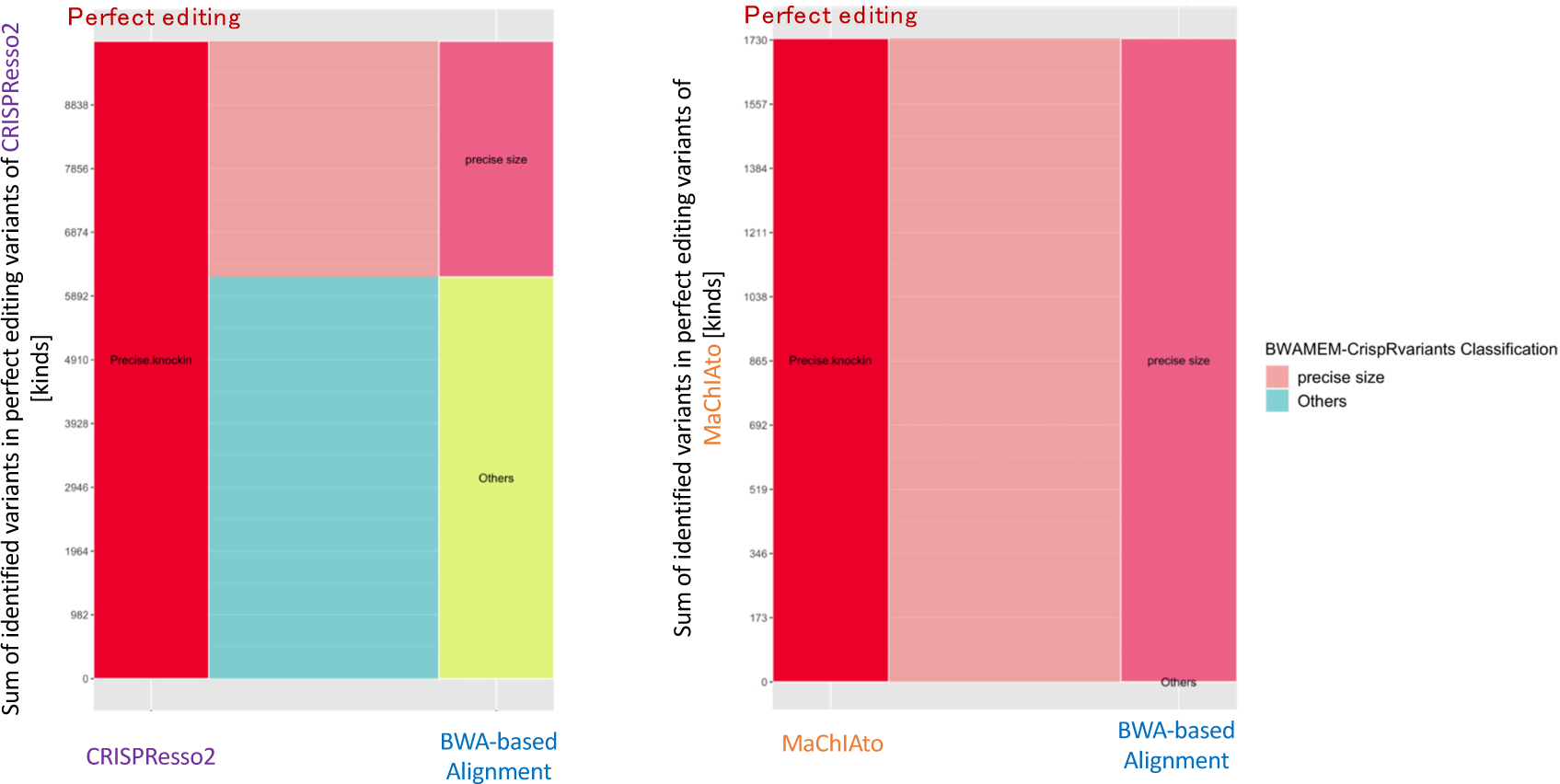
The comparison between BWA MEM-CrispRVariants and CRISPResso2 (left) or MaChIAto (right) using precise knock-in variants in Prime Editing. Left parts of the left and right bar plots indicate the precise knock-in variants (Perfect editing class) of CRISPResso2 and MaChIAto, respectively, and the y-axis is the number of identified variants in the precise editing variants. The right parts of these plots show the variant identification of BWA MEM-CrispRVariants using the corresponding variants. The variants are classified into an “expected length” class and an “Others” class in the variant identification of BWA MEM-CrispRVariants. The variants in the “expected length” class have a sequence that is the expected length of the perfect editing as decided by BWA MEM-CrispRVariants. The flows indicate the change in each class. The variants with the red color flow are classified into the “expected length” class, and the variants with the blue color flow are classified into the “Others” class.

Lastly, we profiled the 29 kinds of amplicon sequencing samples of Prime Editing at the *RNF2*, *RUNX1*, and *VEGFA* loci using MaChIAto Analyzer and MaChIAto Reviewer. MaChIAto Reviewer showed the difference between the groups with high, moderate, and low efficiency in the distributions of the deletion position (Supplementary Fig. 42). The editing construct in the high efficiency group tended to generate the unexpected deletion on the PAM-proximal side, whereas one of the low efficiency groups made many unexpected deletions on the PAM-distal side. MaChIAto Analyzer found some features involved in the precise editing efficiency (Supplementary Fig. 43). Interestingly, the CG motif can increase the editing efficiency, while the GG motif, which has a low free energy parameter for DNA duplex hybridization similar to the CG motif^50, 51^, makes decrease the editing efficiency. Perhaps, the secondary structure of di-nucleotide is more important than the activity for hybridization. For example, the DNA-protein interaction depends on the secondary structure of di-nucleotide. The preferences of GG and CG for kinds of amino acid residue are reportedly different according to a computational study^52^. There might be a di-nucleotide preference in the proteins associated with reverse transcription and/or DNA repair.

**Supplementary Figure 42.**
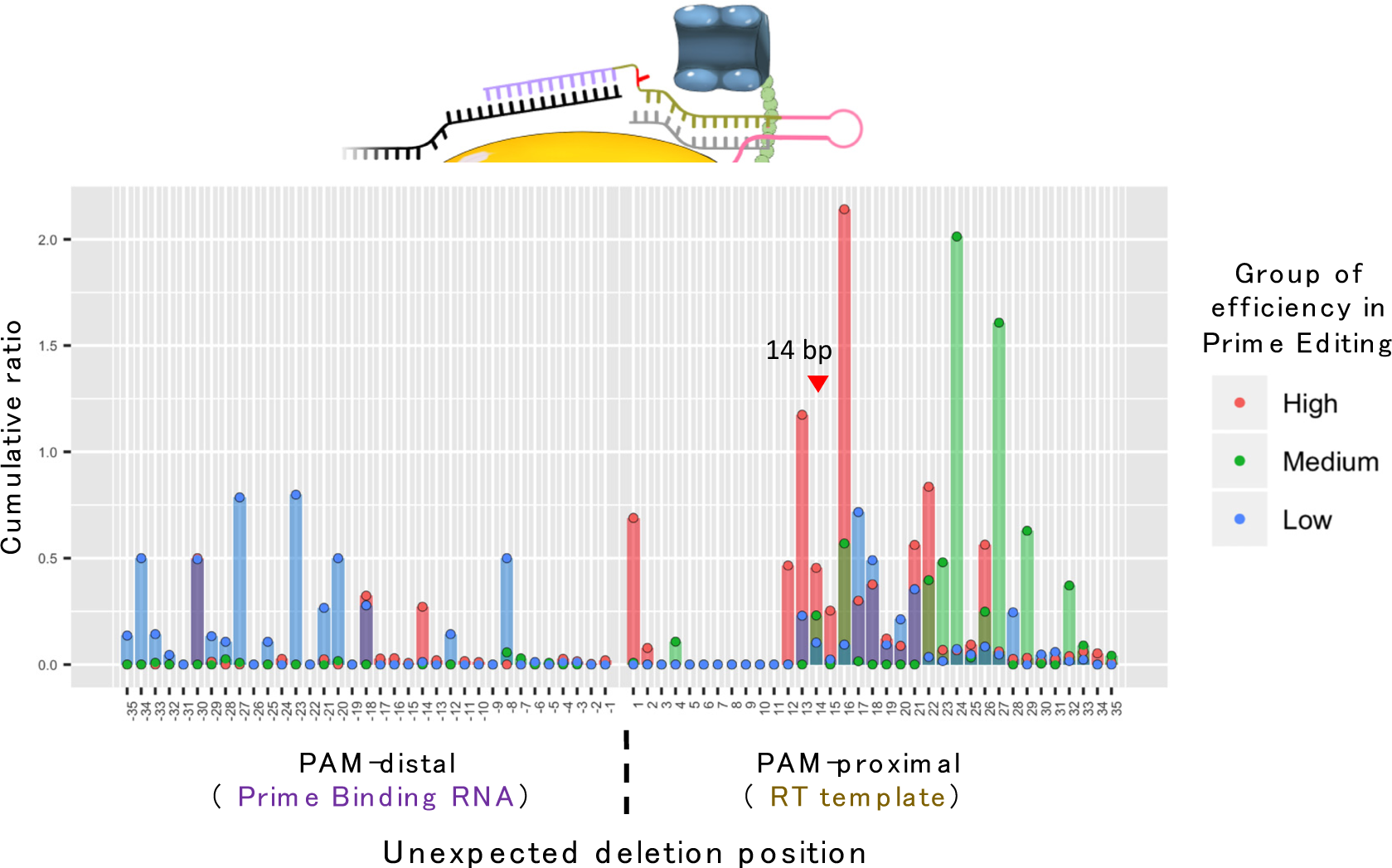
Distribution of deletion position in the InDel class in the Prime Editing of the *RNF2*, *RUNX1*, and *VEGFA* loci. The value is the cumulative ratio of variant labels that MaChIAto Aligner provides. The cumulative ratio was calculated per group. The groups overlap on the plot, and the red, green, and blue points indicate the cumulative ratio of the high, medium, and low groups, respectively, regarding the rate of perfect editing in Prime Editing.

**Supplementary Figure 43.**
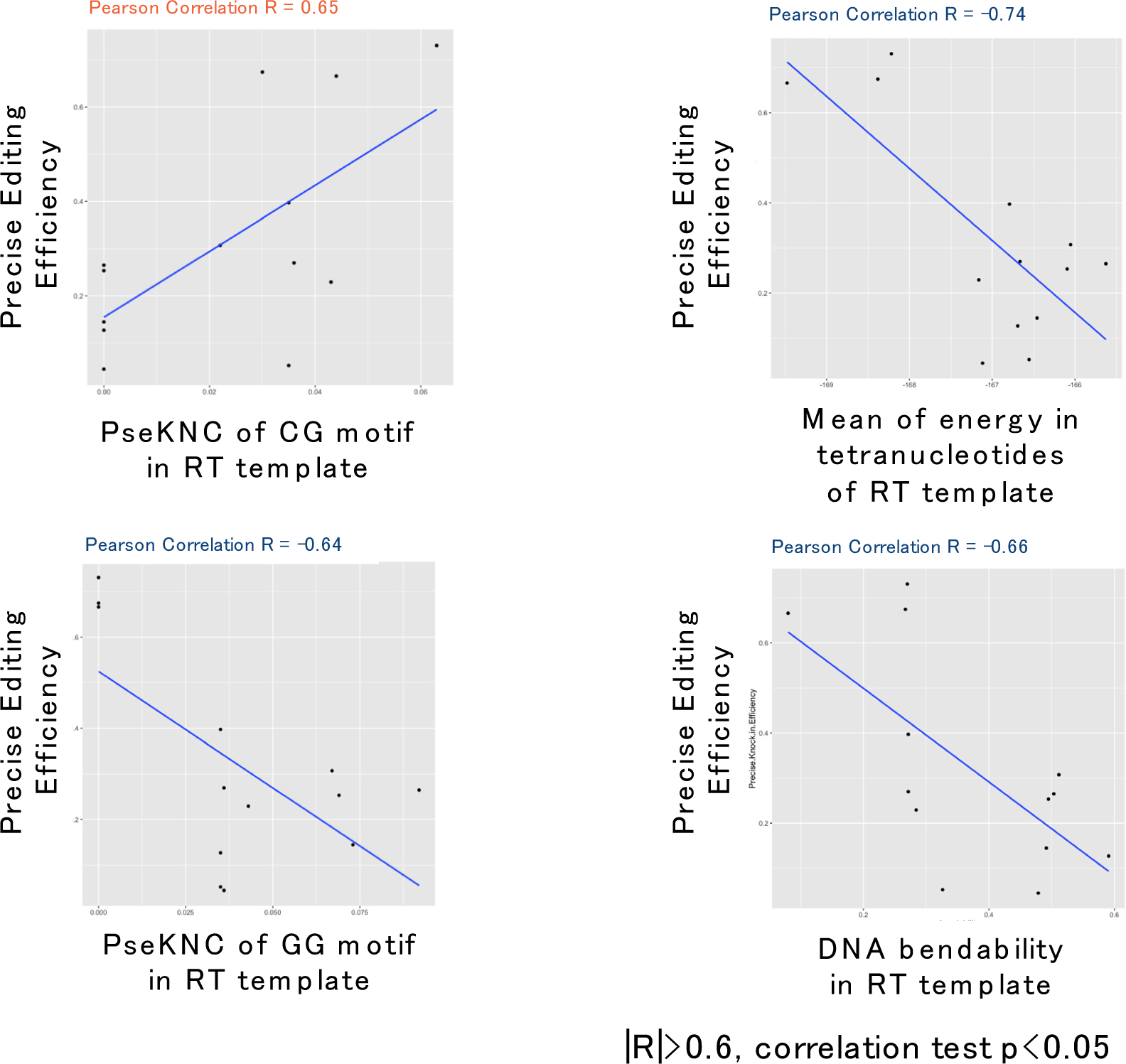
Results of correlation analysis between precise knock-in efficiency (Perfect editing efficiency) and genomic contexts. The y-axis of the scatter plot is the editing efficiency, and the x-axis of that is the value of the feature. The line in the scatter plot is a simple linear regression using the least squares method. The label above the plot represents the Pearson correlation. If the correlation (R) is R > 0.6 or R < 0.6, the label color is orange or blue, respectively. All correlations are statistically significant (p<0.05, correlation test).

In summary, we introduced the analysis mode for Prime Editing. This mode accommodates the function of accurate classification and relationship analysis with mutation patterns and genomic context that could be useful for predicting editing efficiency. MaChIAto analyzer can also analyze the correlation between editing efficiency and user-defined features. MaChIAto would be the comprehensive platform for the quantification of amplicon sequencing data, which enables to quickly find critical characteristics related to the efficacy of Prime Editing.

#### Supplementary Note 10 - Background to the data and tools used

All analyses and development regarding MaChIAto were processed using Anaconda environments. We showed the environment name and the packages in Supplementary Table 6. The execution of MaChIAto Classifier, MaChIAto Analyzer, MaChIAto Aligner and MaChIAto Reviewer were performed under MaChIAto_env, MaChIAtoAnalyzer_env, MaChIAtoAligner_env and MaChIAtoReviewer_env, respectively. The execution of CRISPResso and CRISPResso2 were performed under CRISPResso_env and CRISPResso2_env or in Docker container provided by the author’s GitHub (https://github.com/pinellolab/CRISPResso2#docker). The global alignment of Prime Editing data was performed under the bwa_map_env.

## Notes

https://github.com/KazukiNakamae/MaChIAto

https://machiatopage.github.io

### Competing Interest Statement

The authors have declared no competing interest.

